# A multimodal foundation model to integrate histology with spatial molecular programs

**DOI:** 10.64898/2026.06.01.729028

**Authors:** Zongxu Zhang, Bowen Qin, Yahui Zhao, Zekun Qi, Hanlin Xu, Yunfeng Wang, Wenjin Zheng, Jiateng Dai, Anxin Chen, Nachuan Wang, Lanxi Nie, Peng Zhang, Haorui Zhang, Yanping Zhao, Tian Xu, Siyu Lin, Pengfei Ren, Zhe Zhang, Liyan Xue, Xuemin Xue, Zhaoyang Yang, Jiaqi Xu, Jiahao Ma, Deng Pan, Cunyu Wang, Zhihua Liu, Yuanguang Meng, Zexian Zeng

## Abstract

Histology and spatial transcriptomics provide complementary views of cancer morphology and molecular organization, yet their spatial correspondence has not been learned at scale. We developed SQUALL, a multimodal foundation model pretrained on *histMol*, comprising 1.93 billion spatially aligned histology– transcriptomics spots/bins from 3,446 sections across 33 tissues and 12 platforms. By learning spatial morphology–molecular correspondences, SQUALL enables virtual spatial transcriptomics, tumor microenvironment characterization, and molecularly informed histology representations for clinical prediction. SQUALL generalized virtual spatial gene expression profiling across cohorts, cancers, and platforms. Its representations resolve tumor microenvironments including progression-associated tumor trajectories in breast cancer and recurrence-associated immune-excluded regions in ovarian tumor, together with underlying transcriptional programs. Applied to whole-slide histology, SQUALL enabled outcome and treatment-response prediction while revealing interpretable features associated with risk. Together, these results establish large-scale spatially aligned histology–transcriptomics pretraining as a general framework for connecting tissue morphology, molecular programs, microenvironmental organization, and clinical phenotypes.

## Introduction

Cancer diagnosis and treatment rely heavily on tissue-based assessments. Among available clinical modalities, histopathological review of hematoxylin and eosin (H&E)-stained sections remains a cornerstone of cancer diagnosis, grading, and risk stratification.^1,2^ By resolving tissue architecture, cellular morphology, and tumor-stroma organization, histology provides a direct view of disease states in their native tissue context and guides routine clinical diagnostics and progress evaluation. Yet histological interpretation remains primarily morphology-based, providing only indirect access to the molecular programs that shape tumor progression, therapeutic response, and immune interactions. Bridging histological phenotypes with their underlying molecular states therefore remains an important challenge in cancer biology and pathology.

Advances in molecular profiling have enabled systematic characterization of gene expression programs, signaling pathways, and cellular composition. Bulk and single-cell sequencing studies have identified biomarkers associated with tumor progression,^3^ metastatic potential,^4^ treatment response,^5^ and patient outcome.^6^ However, these approaches often require tissue homogenization or dissociation, disrupting the spatial organization of tumors and their surrounding microenvironments.^7^ Consequently, molecular measurements are frequently separated from the histological structures.^8^

Spatial transcriptomics technologies, as a conceptual convergence of histological imaging and molecular sequencing,^9,10^ provide a means to bridge this gap by measuring molecular states while preserving their tissue context. These technologies enable gene expression programs to be mapped onto histological architecture, revealing spatially organized tumor states,^11^ stromal programs,^12,13^ and immune functions.^14,15^ As spatial transcriptomics continues to expand in scale and resolution,^16,17^ growing collections of paired histology and spatial molecular profiles create an opportunity to systematically learn the relationships between molecular states and the image-based phenotypes routinely observed in pathology.

In parallel, foundation models trained on large biomedical datasets have emerged as a powerful framework for representation learning.^18–21^ In computational pathology, large vision models trained on histology images have improved tumor classification, subtyping, and prognostic prediction from whole-slide images.^22–25^ Most existing pathology foundation models, however, are trained primarily on histology alone or through alignment with text. Without direct learning from spatially resolved molecular measurements in matched tissue sections, these models are not explicitly optimized to capture the molecular programs associated with local histological phenotypes, thus limiting their ability to recover the biological programs underlying histological phenotypes or to generalize mechanistic insight across datasets and platforms.

Here, we present SQUALL, a 555-million-parameter multimodal foundation model trained on paired histology and spatial transcriptomics to integrate tissue morphology with molecular states. To enable large-scale multimodal pretraining, we assembled *histMol*, a corpus of 1.93 billion spatial transcriptomics spots/bins from 3,446 tissue sections across 33 tissues and 12 spatial profiling platforms, spanning diverse imaging and sequencing resolutions. We reasoned that learning the correspondence between histological morphology and spatially resolved gene expression profiles could provide representations that connect tissue phenotypes with their associated molecular programs. After pretraining, SQUALL supports various downstream analyses across diverse cancer types using either joint histology-transcriptomics representations or histology alone. From histology alone, SQUALL enables virtual spatial biomarker profiling of up to 15,757 genes across independent cohorts and platforms. Using joint multimodal representations derived from measured histology and spatial transcriptomics, SQUALL resolves spatially aware tumor microenvironment clusters, reconstructs a progression-associated molecular trajectory in breast cancer across 267,945 spots from 198 sections, and identifies a recurrence-associated immune-excluded spatial region in ovarian tumor from Visium HD sections of 52 patients. Using representations from the pretrained histology encoder alone, SQUALL further improves prognostic prediction from whole-slide images across 770 patients and provides more balanced prediction of platinum-based chemotherapy resistance in two independent cohorts. Together, these findings support multimodal spatial representation learning as a scalable framework for connecting histopathology with tumor biology and extending spatial molecular insights to routine tissue imaging at scale.

## Results

### Collection and preprocessing of the histMol corpus

To enable multimodal pretraining integrating tissue histological morphology and spatial molecular programs, we assembled *histMol*, a large-scale corpus of paired histology and spatial transcriptomics data spanning 33 tissues, 12 platforms, 3,446 tissue sections, and various spatial resolutions (**Figures 1A and S1A-S1H**; **Tables S1-S8**; **Methods**). *histMol* aggregates 1.93 billion sequencing spots/bins from 15 public repositories (308 studies; **Table S1**) together with our in-house generated datasets (n = 198 sections).^26–28^ The corpus incorporates major spatial transcriptomics technologies, including Visium,^10^ ST,^29^ Visium HD,^30^ OpenST,^31^ Nova-ST,^32^ Seq-Scope,^33^ and Stereo-seq,^34^ each paired with matched H&E images from the same tissue sections (**Table S2**). Overall, *histMol* encompasses diverse anatomical sites, experimental protocols, and human, mouse, and xenograft specimens (**Tables S3-S8**), providing a broad resource for multimodal representation learning (**Figure 1B**).

**Figure 1.**
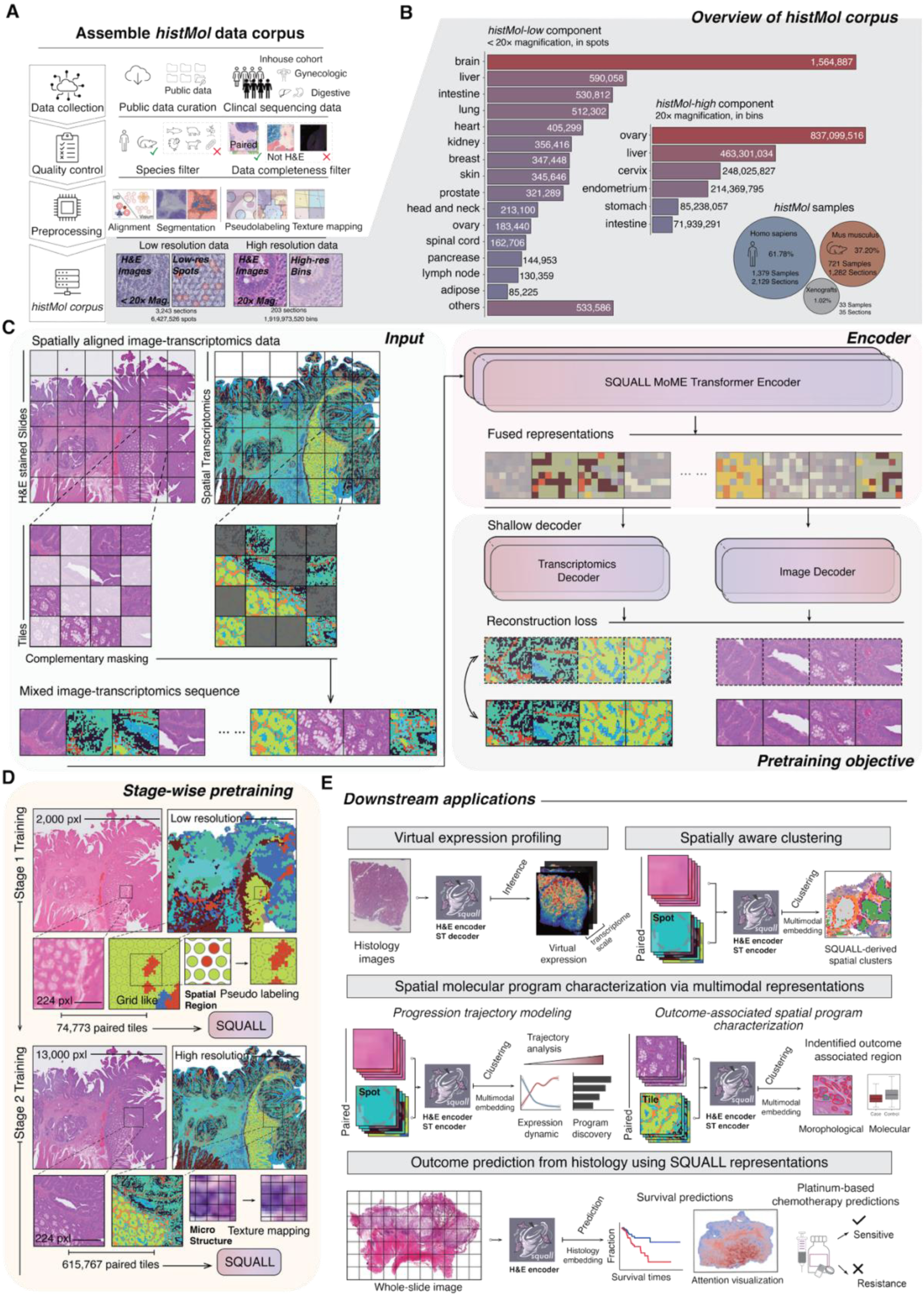
SQUALL is a multimodal foundation model that integrates histology with spatial molecular programs. **(A)** Assembly of the *histMol* corpus for multimodal pretraining. Publicly available spatial transcriptomics (ST) datasets were integrated with newly collected in-house data. Only samples containing matched hematoxylin and eosin (H&E)–stained histology images and ST profiles from the same tissue section were retained. Following quality control, preprocessing, image–transcriptome alignment, segmentation, pseudo-labeling, and texture mapping, the resulting corpus comprised approximately 1.93 billion sequencing spots or bins. See also Figures S1 and S2. **(B)** Overview of the *histMol* corpus. The dataset comprises 3,446 sections spanning 33 anatomical tissues from human, mouse, and xenograft samples and integrates multiple ST technologies. Fraction indicates the proportion of sections. See also Figure S1. **(C)** Overview of the SQUALL architecture and pretraining objective. Spatially aligned H&E images and transcriptomic profiles are partitioned into paired tiles and subjected to complementary masking. The resulting mixed image–transcriptomics sequences are processed by the SQUALL mixture-of-modality-experts (MoME) Transformer encoder to learn shared representations of tissue morphology and gene expression programs. Modality-specific shallow decoders reconstruct the masked histological and transcriptomic inputs. See also Figure S3. **(D)** Stage-wise multimodal pretraining of SQUALL on the *histMol* corpus across spatial resolutions. In the first stage, SQUALL is trained on low-resolution data to capture broad relationships between tissue morphology and molecular organization. In the second stage, high-resolution data are used to learn fine-grained correspondences among cellular and microanatomical structures, histological textures, and gene expression patterns. **(E)** Downstream applications of SQUALL across spatial scales. At the spot- and tile-level, SQUALL supports virtual gene expression profiling, spatially aware clustering, progression-trajectory modeling, and characterization of outcome-associated spatial programs. At the whole-slide level, SQUALL representations enable clinical outcome prediction directly from histology, including survival and treatment-resistance prediction, with attention-based visualization of predictive tissue regions.

To support cross-species learning, we defined a shared gene space of 15,757 one-to-one human-mouse orthologs (**Figure S1A**) that was uniformly distributed across cytobands (**Figure S1B**). Compared with large-scale single-cell datasets such as Genecorpus-30M,^35^ spatial transcriptomics profiles were markedly sparse (**Figure S1C**). Across platforms, library size correlated with the number of detected genes per spot/bin significantly (mean Pearson’s r = 0.98; **Figure S1D**), indicating that gene-detection status retained substantial biological information. Similar data binarization has previously been shown to preserve biologically meaningful signals for transcriptomic analysis,^36–42^ and has also served as a reconstruction target for foundation model pretraining.^43^ We therefore binarized gene expression to preserve informative presence-absence patterns while improving modeling efficiency across platforms and resolutions.

To capture complementary spatial scales, we partitioned *histMol* into low-resolution (*histMol*-low; **Tables S5-S7**) and high-resolution (*histMol*-high; **Table S8; Figure S1E**) subsets according to imaging and sequencing resolution (**Table S2**). *histMol*-low provided broad coverage of tissue and cancer types for learning global morphology-molecular relationships across various resolutions (**Figure S1F**), whereas *histMol*-high captured fine-grained microanatomical correspondence at higher resolution. We developed resolution-specific preprocessing pipelines (**Methods**) incorporating background white-balance correction,^44^ pseudo-labeling (**Figures S2A-S2E**), and texture mapping (**Figures S2F-S2L**).

The final corpus comprised 6,427,526 spots from 3,243 *histMol*-low sections representing 1,952 samples and 1,919,973,520 bins from 203 *histMol*-high sections representing 181 samples (**Figure 1B**; **Table S1**). This scale substantially exceeded that of HEST-1k and STimage-1K4M (**Figure S1G**).^45,46^ After preprocessing, *histMol* contained 74,773 low-resolution and 615,767 high-resolution tiles (**Figure S1H**), establishing a large-scale resource for multimodal learning in spatial biology.

**Figure S1.**
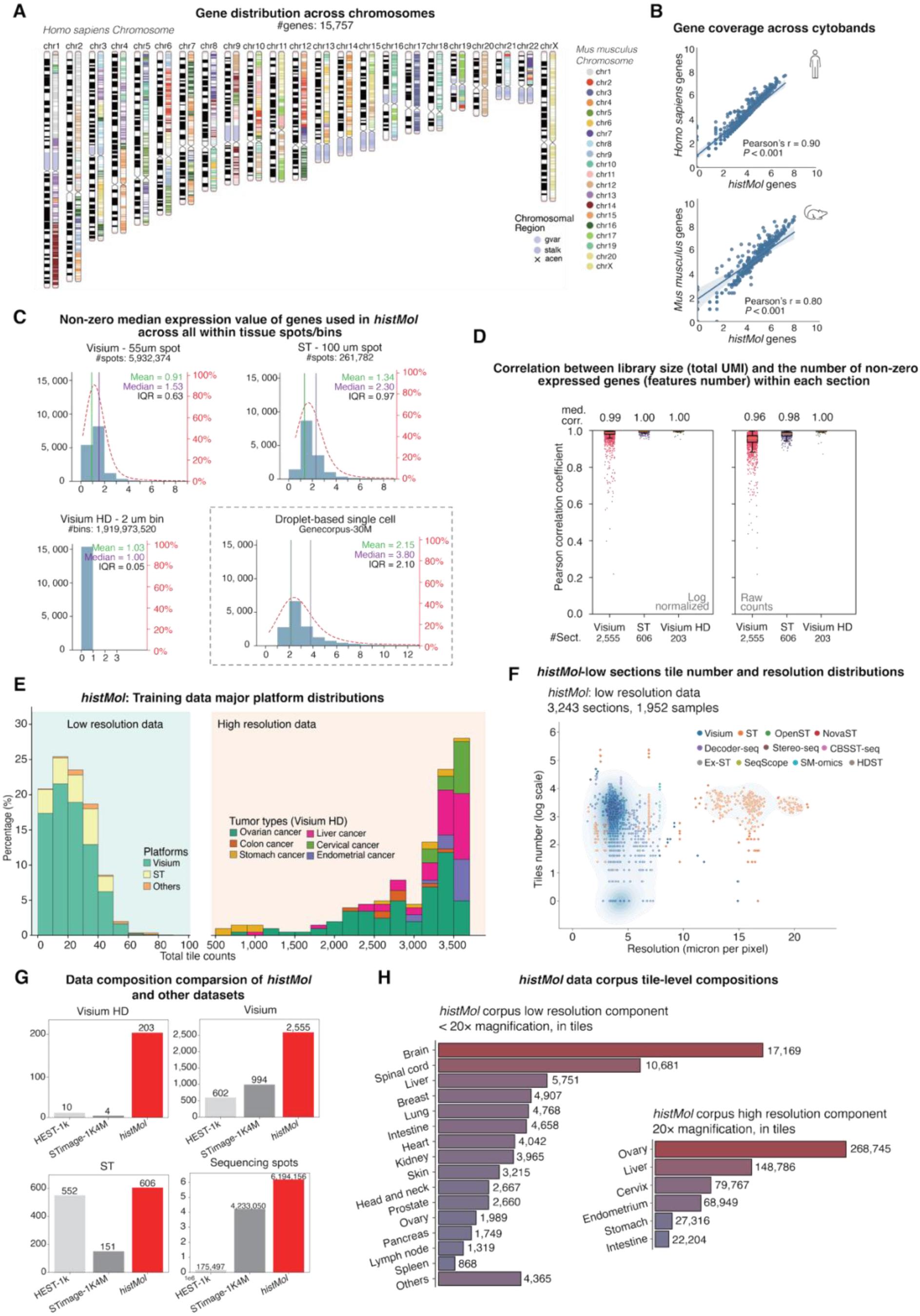
Overview of the *histMol* pretraining corpus. Related to Figure 1. **(A)** Ideogram showing the chromosomal distribution of the 15,757 one-to-one human-mouse orthologous genes retained in the *histMol* corpus. Genes are annotated according to their corresponding mouse chromosomes and genomic regions. **(B)** Scatter plots comparing the cytoband-level coverage of *histMol* genes with that of all annotated genes in the human genome (top) and mouse genome (bottom). Pearson’s correlation coefficients and corresponding *P* values are shown. **(C)** Distributions of the median genes with nonzero expression included in *histMol* across tissue spots or bins from the Visium, ST, and Visium HD subsets, compared with droplet-based single-cell RNA-sequencing data from Genecorpus-30M. Means, medians, and interquartile ranges (IQRs) are indicated; red dashed lines show the cumulative percentages of genes. **(D)** Distributions of section-level Pearson correlation coefficients between library size (total UMI count) and the number of genes with nonzero expression across major spatial transcriptomics (ST) platforms in *histMol*, calculated using log-normalized expression values (left) or raw counts (right). Median correlation coefficients are shown above each distribution. Box plots show the median (center line), upper and lower quartiles (box limits), and values within 1.5 times the IQR (whiskers). **(E)** Distributions of the number of training tiles generated per tissue section in the *histMol*-low (left) and *histMol*-high (right) subsets. The *histMol*-low distribution is stratified by ST platform, whereas the *histMol*-high distribution is stratified by tumor type. **(F)** Relationship between histology image resolution and the number of tiles generated per section in the *histMol*-low subset, comprising 3,243 tissue sections from 1,952 samples. Points are colored by ST platform, and tile numbers are shown on a logarithmic scale. **(G)** Comparison of the data volume in *histMol*, HEST-1K, and STimage-1K4M. The numbers of Visium HD sections, Visium sections, ST sections, and spatially resolved sequencing spots are shown. **(H)** Tissue composition of the *histMol*-low (left; imaging magnification below 20×) and *histMol*-high (right; 20× imaging magnification) subsets, reported as the number of training tiles derived from each tissue type.

**Figure S2.**
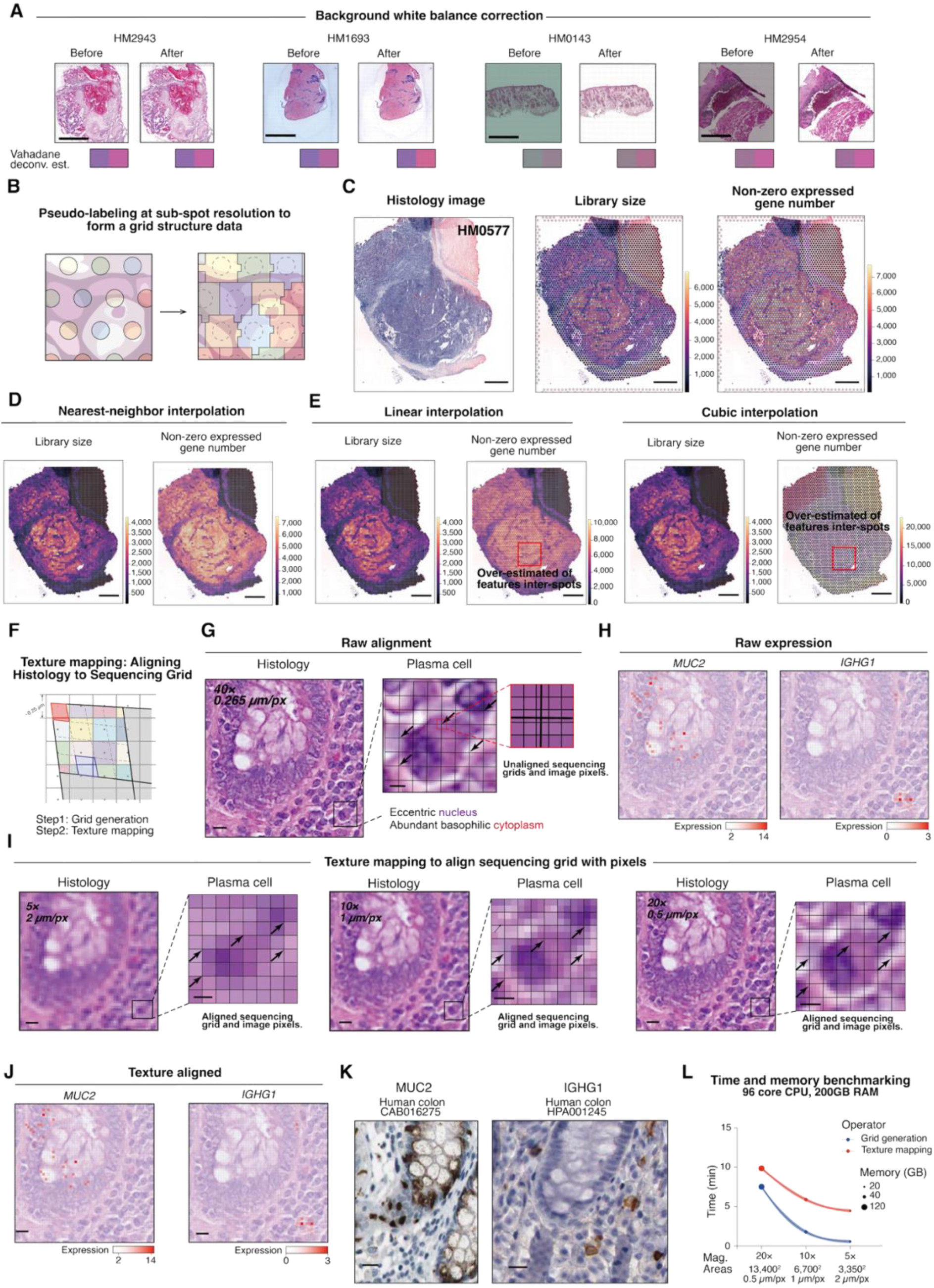
Data preprocessing pipeline for the *histMol* corpus. Related to Figure 1. **(A)–(E)** Preprocessing of *histMol-*low component. **(A)** Background white-balance correction of histology images. Representative images before and after correction are shown, together with stain vectors estimated using the Vahadane method for stain separation. Scale bar, 2 mm. **(B)** Schematic of the pseudo-labeling procedure used to construct a regular, grid-like data structure. Original sequencing spots were subdivided into sub-spots, and gaps between adjacent spots were filled by nearest-neighbor interpolation. **(C)** Representative *histMol*-low section showing the white-balanced H&E image (left), library size measured as the total UMI count (middle), and the number of genes with nonzero expression (right). **(D)** Sub-spot-level library size (left) and number of genes with nonzero expression (right) after pseudo-labeling using nearest-neighbor interpolation of the data shown in **(C)**. **(E)** Sub-spot-level library size and number of genes with nonzero expression generated from the data in **(C)** using linear interpolation (left) or cubic interpolation (right). Both smoothing-based methods overestimated expression between the original sequencing spots, producing checkerboard-like artifacts in the highlighted regions. Scale bars in **(C)–(E)**, 1 mm. **(F) –(L)** Preprocessing of *histMol*-high component **(F)** Schematic of the texture-mapping procedure used to align the ST sequencing grid with histology image pixels. The procedure comprises generation of an image-aligned grid at the target resolution (step 1), followed by texture mapping of the original image pixels onto the grid (step 2). **(G)** Example of misalignment between a raw histology image and its corresponding sequencing grid. A 224 × 224-pixel histology tile from *histMol*-high is shown on the left. Enlarged views of a plasma cell illustrate the displacement between the histological structure and sequencing bins (middle), followed by a further magnified view of the mismatch between the original image pixels and bin boundaries (right). Bold grid lines indicate unaligned sequencing-bin boundaries. Scale bars, 10 µm for the image tile and 2 µm for the enlarged views. **(H)** Raw, unaligned spatial expression maps of *MUC2* and *IGHG1* in the region shown in **(G)**. *MUC2* marks normal colonic goblet cells, whereas *IGHG1* marks plasma cells. **(I)** Texture mapping aligns the sequencing grid with histology image pixels at 5× (left), 10× (middle), and 20× (right) magnification. Arrows indicate corresponding bin borders and corners after alignment. Scale bars, 10 µm for the image tiles and 2 µm for the enlarged views. **(J)** Spatial expression maps of *MUC2* and *IGHG1* after texture mapping, shown for the same tissue region as in **(H)**. **(K)** Immunohistochemical staining of MUC2 and IGHG1 in human colon tissue, obtained from the Human Protein Atlas, used to validate the correspondence between spatial gene expression and histological cell morphology after texture mapping. Scale bars, 20 µm for the immunohistochemistry images and 10 µm for the ST tiles. **(L)** Computational scalability of grid generation and texture mapping across image magnifications and pixel counts. Processing time increased with image pixel count, while point size represents peak memory usage. Lines show the mean and shaded regions indicate the 95% confidence interval across three independent runs.

### SQUALL architecture, multimodal pretraining, and design evaluation

Using *histMol*, we developed SQUALL, a multimodal foundation model that integrates histological morphology with spatial gene-expression programs (**Figures 1C and S3A-S3D**). Based on masked data modeling (**Figure S3A**; **Methods**),^47,48^ SQUALL learned joint representations from paired histology and spatial transcriptomics by masking complementary regions across modalities and reconstructing the missing signals (**Figure S3B**). SQUALL adopts a mixture-of-modality-experts (MoME) Transformer that combines modality-specific feature extraction with shared multimodal fusion layers.^49,50^ This modularized design enables joint multimodal learning while supporting both multimodal and histology-only inference for downstream analyses (**Figure S3C**; **Table S9**). To accommodate the diverse spatial resolutions across technologies, we explicitly encoded spatial resolution and coordinates (**Figure S3D**; **Methods**).^51,52^ We further adopted a stage-wise continual-pretraining strategy (**Figure 1D**), in which SQUALL was first pretrained on low-resolution tiles to learn global morphology-molecular alignment and subsequently on high-resolution tiles to capture fine-scale microanatomical correspondence. Together with its modular architecture, this strategy enables SQUALL to support flexible downstream applications using either joint multimodal representations or histology alone, across spatial scales ranging from sequencing spots to whole-slide histopathology images (**Figure 1E**; **Table S9**).

Image classification evaluations identified several factors contributing to model performance (**Figures S3E-S3I**; **Tables S10-S25**; **Methods**). Combining absolute and relative positional encodings with an explicit resolution parameter yielded the highest accuracy (**Figure S3F**), and increasing model capacity, training duration further improved performance (**Figure S3F**). Stain normalization modestly reduced performance, whereas increasing data diversity by incorporating mouse tissues and multiple platforms during pretraining improved classification performance compared with human-only or Visium-only training (**Figure S3G**). For high-resolution image classification (**Figure S3H**; **Table S22**), stage-wise pretraining outperformed training on either corpus alone (**Figure S3I**). Additional evaluations on downstream tasks confirmed favorable overall performance of the final SQUALL design across choices involving data composition, foundation-model initialization, normalization, and interpolation methods used for pseudo-labeling. (**Figure S3J**; **Table S26**).

Using the optimized training protocol (**Tables S27-S28**), SQUALL converged stably during pretraining (**Figure S3K**). Masked token reconstruction results further demonstrated that SQUALL recovered salient histological features and gene-expression patterns (**Figure S3L**), suggesting that the learned representations captured informative cross-modal features.

**Figure S3.**
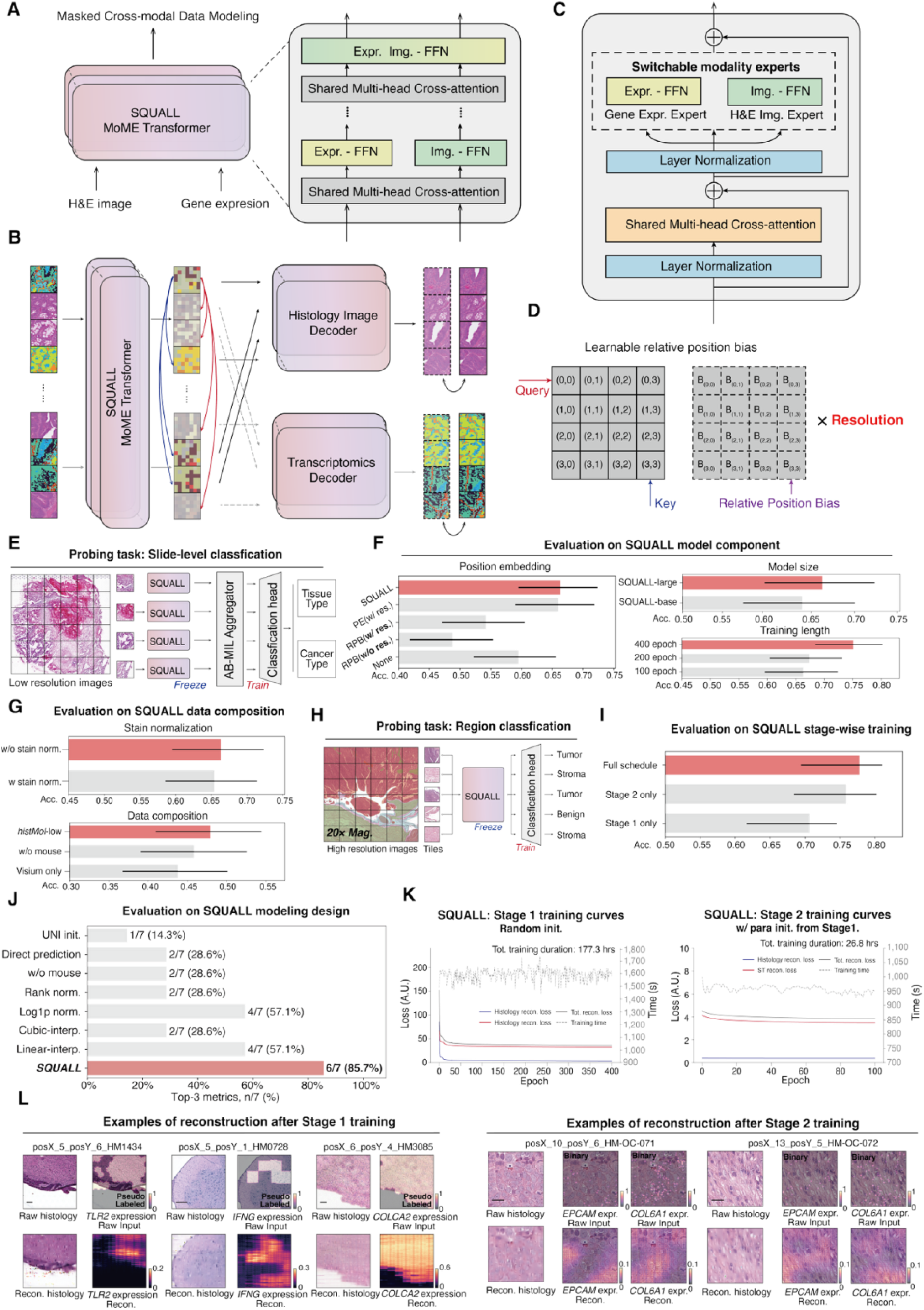
SQUALL model architecture, design evaluations, and training details. Related to Figure 1. **(A)–(D)** Architecture and pretraining framework of SQUALL. **(A)** Overview of masked cross-modal data modeling using the SQUALL mixture-of-modality experts (MoME) Transformer. Paired H&E image and gene expression tokens are processed using modality-specific feed-forward networks and integrated through shared multi-head cross-attention layers. **(B)** Cross-modal reconstruction framework used during pretraining. Masked histology and gene expression tokens are jointly encoded by the SQUALL MoME Transformer and reconstructed using modality-specific histology-image and transcriptomics decoders. Cross-modal attention enables information exchange between the two modalities during reconstruction. **(C)** Architecture of a SQUALL MoME Transformer block. Each block comprises shared multi-head cross-attention layers and switchable modality-specific feed-forward experts for histology images and gene expression, with residual connections and layer normalization. **(D)** Learnable relative positional bias used in the attention layers. Relative spatial offsets between query and key tokens are encoded and scaled according to the spatial resolution of the input data. **(E)** Slide-level classification probing task used to evaluate representation quality and perform model ablations. Features were extracted from low-resolution image tiles using a frozen SQUALL encoder and aggregated using a trainable attention-based multiple-instance learning (ABMIL) module and classification head to predict tissue or cancer type. **(F)** Evaluation of SQUALL model components using the slide-level tissue-classification probing task. Left, comparison of positional-information encoding strategies, including position embedding (PE), relative position bias (RPB) with or without spatial resolution constraints, and no positional encoding. Right, effects of model size (top) and pretraining duration (bottom). Bars show classification accuracy, and error bars indicate 95% confidence intervals. PE, position embedding; RPB, relative position bias; w/ res., with spatial resolution constraints; w/o res., without spatial resolution constraints; Acc., accuracy. **(G)** Evaluation of preprocessing and pretraining data composition. Top, effect of Macenko stain normalization using the median stain-color vector estimated from *histMol*-low. Bottom, effects of removing mouse tissue data or restricting pretraining to Visium data. Including mouse tissues and multiple low-resolution ST platforms improved downstream classification accuracy. Bars show classification accuracy, and error bars indicate 95% confidence intervals. **(H)** Tissue region classification probing task used to evaluate stage 2 representations. Features were extracted from high-resolution image tiles using a frozen SQUALL encoder, and a trainable classification head was used to predict tissue-region annotations. **(I)** Evaluation of the SQUALL stage-wise training strategy. Tissue-region classification accuracy was compared among the complete training schedule, stage 2 training alone, and stage-1 training alone. Bars show classification accuracy, and error bars indicate 95% confidence intervals. **(J)** Evaluation of SQUALL design choices across downstream tasks using seven metrics: PCC, AUC, SSIM, ARI, AMI, C-index, and F1. Bars indicate the number and proportion of metrics for which each configuration ranked among the top three. UNI init., image feature derived from pretrained UNI encoder; Direct pred., direct gene expression prediction from histology; w/o mouse, mouse data excluded from pretraining; Rank norm. and Log1p norm., rank-based and log1p normalization of ST data; Cubic interp. and Linear interp., cubic and linear interpolation of low-resolution ST data; init., initialization; norm., normalization; interp., interpolation; w/o, without. **(K)** Training curves for stage 1 (left) and stage 2 (right). Histology reconstruction loss, gene expression reconstruction loss, total reconstruction loss, and per-epoch training time are shown. Stage 1 was trained from random initialization, whereas stage 2 was initialized using parameters learned in stage 1. The total training duration for each stage is indicated above the corresponding plot. ST., spatial transcriptomics; Tot., total; recon., reconstruction; expr., expression; init., initialization. **(L)** Representative reconstruction results after stage-1 training (left) and stage-2 training (right). Raw histology images and gene expression inputs are shown together with their corresponding SQUALL reconstructions. Stage-1 examples illustrate reconstruction of pseudo-labeled expression profiles at low spatial resolution, whereas stage-2 examples illustrate reconstruction of histology and marker-gene expression at high spatial resolution. Recon., reconstruction; expr., expression.

### SQUALL enables virtual spatial gene expression profiling from histology

Given the widespread availability of histology imaging compared with molecular profiling assays, virtual inference of clinically relevant molecular biomarkers from H&E images could provide a scalable approach for extending molecular characterization to routine pathology workflows.^9,53^ We therefore tested whether SQUALL, after large-scale multimodal pretraining on paired histology and spatial transcriptomics data, could virtually profile spatial gene expression directly from histology. We froze the histology encoder and transcriptomics decoder and generated spatial gene expression profiles from H&E images in a single forward pass using SQUALL (**Figure 2A**; **Methods**).

**Figure 2.**
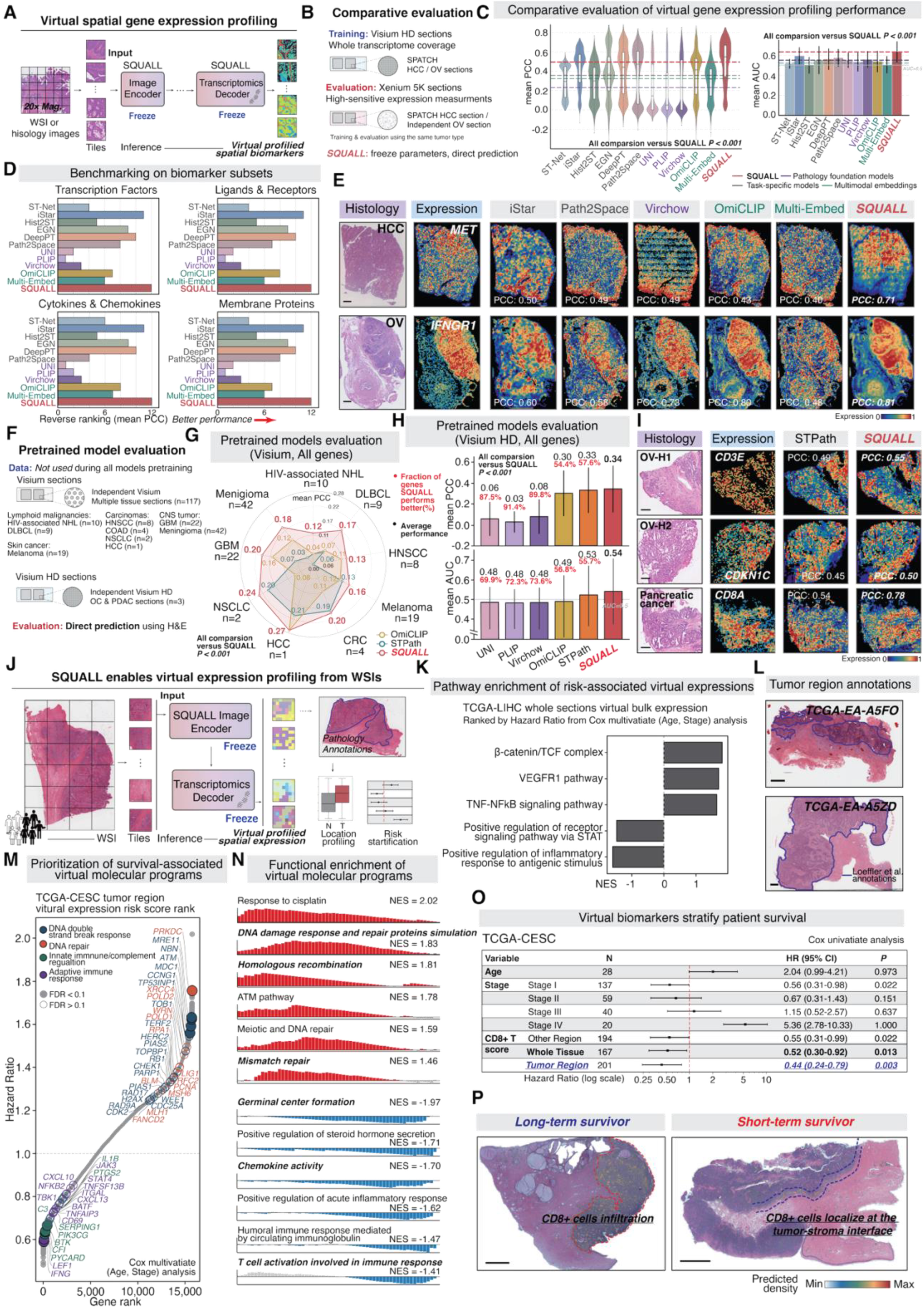
SQUALL enables virtual spatial gene expression profiling from histology and reveals spatial gene expression programs associated with survival. **(A)** Workflow for virtual spatial gene expression profiling directly from histology. Histological tiles are processed by the frozen SQUALL image encoder, and the resulting representations are converted into spatial gene expression profiles by the frozen transcriptomics decoder. **(B)** Data and experimental design for comparative evaluation of virtual gene expression profiling. Compared methods were trained on Visium HD sections to learn associations between tissue morphology and gene expression and were subsequently evaluated on Xenium 5K sections from the same tumor types. OmiCLIP predictions were generated using its pretrained image–ST similarity matrix. HCC, hepatocellular carcinoma; OV, ovarian carcinoma. **(C)** Comparison of virtual gene expression profiling in the comparative evaluation setting. Performance was quantified at the gene-level across 4,421 shared genes using the mean Pearson correlation coefficient (PCC; left) and mean area under the receiver operating characteristic curve (AUC; right). Error bars indicate standard deviations. Comparisons with SQUALL were performed using two-sided Wilcoxon signed-rank tests. See also Figures S4A–S4C. **(D)** Performance across functional biomarker subsets. Genes were grouped into transcription factors, ligands and receptors, cytokines and chemokines, and membrane proteins. Methods were reverse-ranked according to the mean gene-wise PCC within each subset, such that a higher rank indicates better performance. **(E)** Representative virtual biomarker expression profiles. From left to right, panels show the original H&E-stained histology image at 20× magnification, measured expression, and expression profiles predicted by the indicated methods. Rows show *MET* expression in HCC and *IFNGR1* expression in OV. Scale bars, 1 mm. See also Figure S4C. **(F)** Data and experimental design for pretrained model evaluation. Newly collected Visium sections (n = 117) and Visium HD sections (n = 3), spanning multiple cancer types, were collected after pretraining of SQUALL and all comparator histology-ST models was completed. Virtual expression profiles were generated by direct inference from the corresponding H&E-stained images without additional model fine-tuning for pretrained histology-ST models. **(G)** Radar plot summarizing virtual gene expression profiling performance on Visium sections across cancer types. Values represent mean gene-wise PCCs across 14,766 shared genes. Comparisons with SQUALL were performed using two-sided Wilcoxon signed-rank tests. NHL, non-Hodgkin lymphoma; DLBCL, diffuse large B-cell lymphoma; COAD, colon adenocarcinoma; NSCLC, non-small cell lung cancer; GBM, glioblastoma; HNSCC, head and neck squamous cell carcinoma. See also Figures S4D–J. **(H)** Virtual gene expression profiling performance on Visium HD sections. Pathology foundation models were evaluated in fivefold cross-validation using data split into training, validation, and test sets at a ratio of 3:1:1, and results were aggregated across held-out evaluation folds. Bars show the mean gene-wise PCC (top) and AUC (bottom) across 15,757 shared genes; percentages indicate the fraction of genes for which SQUALL outperformed the corresponding method. Error bars indicate standard deviations. Comparisons with SQUALL were performed using two-sided Wilcoxon signed-rank tests. **(I)** Representative virtual biomarker expression profiles from Visium HD sections. From left to right, panels show the original H&E-stained histology image at 20× magnification, measured expression, and expression predicted by STPath and SQUALL. Rows show *CD3E* in ovarian cancer sample OV-H1, *CDKN1C* in ovarian cancer sample OV-H2, and *CD8A* in pancreatic cancer. Scale bars, 1 mm. **(J)** Application of SQUALL to virtual expression profiling from histological whole-slide images (WSIs). Image tiles are processed by the frozen SQUALL image encoder and transcriptomics decoder to generate spatially resolved virtual expression profiles. These profiles can be integrated with pathological annotations to characterize region-specific biomarkers and stratify clinical risk. **(K)** Gene set enrichment analysis (GSEA) of SQUALL-predicted virtual expression profiles across the TCGA-LIHC cohort. Genes were ranked by coefficients estimated using multivariable Cox proportional hazards models adjusted for age and stage. Pathways with normalized enrichment scores (NESs) greater than zero were associated with poorer prognosis, whereas those with NESs less than zero were associated with better prognosis. **(L)** Representative tumor-region annotations in TCGA-CESC WSIs, based on annotations from Loeffler et al. Scale bars, 1 mm. **(M)** Prioritization of survival-associated virtual molecular programs within annotated tumor regions of the TCGA-CESC cohort. Genes are ranked by hazard ratios estimated using multivariable Cox proportional hazards models adjusted for age and stage. Representative genes involved in the DNA double-strand break response (blue), DNA repair (orange), innate immune and complement regulation (green), and adaptive immune responses (violet) are highlighted. Solid circles denote genes with a false discovery rate (FDR) below 0.1. **(N)** GSEA of SQUALL-predicted virtual expression profiles within annotated tumor regions of the TCGA-CESC cohort. Positively enriched pathways were associated with increased risk, whereas negatively enriched pathways were associated with reduced risk. NES, normalized enrichment score. **(O)** Associations between clinical variables, SQUALL-predicted regional CD8⁺ T cell signatures, and survival in the TCGA-CESC cohort. Hazard ratios were estimated using univariate Cox proportional hazards models. Points indicate hazard ratios, and horizontal bars indicate 95% confidence intervals. Statistical significance was assessed using one-sided Wald tests. **(P)** Representative spatial distributions of SQUALL-predicted CD8⁺ T cell signatures in long- and short-term survivors from the TCGA-CESC cohort. In the long-term survivor (left), CD8⁺ T cell infiltration is observed within the tumor region, whereas in the short-term survivor (right), CD8⁺ T cells are predominantly localized at the tumor–stroma interface. Color indicates the predicted density of the CD8⁺ T cell signature. Scale bars, 5 mm. See also Figures S5F–H.

We first evaluated SQUALL’s performance in a comparative setting (**Figure 2B**; **Tables S29-S30**; **Methods**) comprising three model categories: (1) task-specific models, including ST-Net,^54^ iStar,^55^ and Path2Space,^56^ together with three H&E-to-ST models: Hist2ST,^57^ EGN,^58^ and DeepPT,^59^ selected based on their superior overall performance in a benchmark analysis by Wang et al;^60^ (2) pretrained pathology foundation models, including UNI,^23^ PLIP,^61^ and Virchow;^62^ and (3) multimodal H&E-ST joint embedding models, including OmiCLIP and Multi-Embed.^63,64^ To enable a controlled comparison, we trained the task-specific models and pathology foundation models (**Table S31**) on SPATCH Visium HD sections and evaluated all models on independent Xenium 5K sections, with no patient-or section-level overlap with SQUALL pretraining. OmiCLIP predictions were derived from its pretrained image-ST similarity matrix, whereas SQUALL remained fully frozen during inference.

Across the 4,421 genes shared between Xenium 5K and *histMol*, SQUALL achieved greater predictive accuracy and AUC than all comparators (*P* < 0.001 for all comparisons, two-sided Wilcoxon signed-rank test), with a mean gene-wise Pearson correlation coefficient (PCC) of 0.50 and an AUC of 0.63 (**Figures 2C and S4A**; **Table S32**). Across these comparisons, task-specific models achieved mean PCC and AUC values of 0.36 and 0.55, respectively; pathology foundation models achieved 0.23 and 0.55; and multimodal embedding methods achieved 0.33 and 0.52. SQUALL retained this advantage among the top highly variable genes (HVGs; **Figure S4B**), supporting the value of aligned H&E-ST pretraining for virtual gene expression profiling from histology alone. Performance gains were consistent across curated biomarker classes (**Tables S33 and S34**), including transcription factors,^65,66^ ligands and receptors,^67^ cytokines and chemokines,^68^ and membrane proteins (**Figure 2D**).^69^ Representative predictions further illustrated the spatial fidelity of virtually profiled biomarkers (**Figures 2E and S4C**). As an example, SQUALL improved spatial concordance for *MET* (**Figure 2E)**, a growth factor receptor implicated in invasion and metastasis, in hepatocellular carcinoma and recovered the immune-signaling receptor *IFNGR1* in ovarian carcinoma.

We next systematically evaluated virtual expression profiling across cancer types using an independent cohort comprising 117 Visium sections (**Figure 2F**; **Table S35**). We compared SQUALL with the pretrained H&E-ST foundation models OmiCLIP and STPath (**Table S36**; **Methods**).^63,70^ Specifically, all sections were collected after SQUALL pretraining and after release of the comparator pretrained weights. This cohort spanned nine cancer cohorts across eight tissues: HIV-associated non-Hodgkin lymphoma (HIV-associated NHL),^71^ diffuse large B cell lymphoma (DLBCL),^72^ melanoma,^73,74^ head and neck squamous cell carcinoma (HNSCC),^75^ colorectal cancer (CRC), non-small cell lung cancer (NSCLC), hepatocellular carcinoma (HCC),^76^ glioblastoma (GBM),^77^ and meningioma.^78^ Across the 14,766 genes shared by all models, SQUALL achieved significantly higher mean gene-wise PCC and AUC than both comparators (*P* < 0.001 for all comparisons, two-sided Wilcoxon signed-rank test; **Figures 2G and S4D**; **Table S37**), with consistent gains across different numbers of top HVGs (**Figure S4E**). Although performance declined for rare cancer types underrepresented in the pretraining data, including lymphoid malignancies (HIV-associated NHL and DLBCL) and meningioma, SQUALL still outperformed OmiCLIP and STPath at both gene and section levels (**Figure S4F**; **Table S37**). Exploratory analyses associated lower performance with poorer staining quality (**Figures S4G-S4H**), characterized by stronger eosin staining (Pearson’s r = -0.41, *P* < 0.0001) and greater variability in staining intensity (r = -0.35, *P* < 0.0001). Representative predictions included *CD8A*, a cytotoxic T cell marker, in HNSCC; *CD79A*, a B cell lineage marker, in HIV-associated NHL; and *SSTR2*, a tumor-associated biomarker, in meningioma (**Figure S4I**). On newly collected independent Visium HD sections (**Table S38**), SQUALL also outperformed Virchow, PLIP, UNI, OmiCLIP, and STPath across 15,757 shared genes (**Figure 2H**; **Table S39**), including the T cell-associated markers *CD3E* and *CD8A* and the cell-cycle-related gene *CDKN1C* (**Figures 2I and S4J**).

In parallel, H&E-based spatial protein models have been developed for virtual biomarker profiling. We therefore compared SQUALL with the pretrained spatial protein models ROSIE and GigaTIME (**Table S40**),^79,80^ to assess its ability to recover spatial biomarker patterns relative to models developed specifically for protein prediction (**Figure S5A**; **Methods**). Datasets that jointly profile histology, spatial proteomics, and spatial transcriptomics in the same tumor tissue sections remain limited and typically contain small protein panels generated using heterogeneous protocols, complicating direct histology-based comparisons.^81–83^ Given previous reports of concordance in spatial marker organization and cell-type patterns between spatial transcriptomic and proteomic measurements,^26,84^ we therefore used whole-transcriptome-scale Visium HD data as a transcript-level reference to maximize the number of matched gene-protein targets available for evaluation. We restricted the analysis to non-secreted biomarkers with unambiguous one-to-one gene-protein mappings (n = 10 for GigaTIME; n = 34 for ROSIE; **Table S41**) and evaluated their spatial concordance across multiple spatial scales to determine whether spatial correspondence was maintained across resolutions. The independently collected Visium HD sections had not been used to pretrain any of the evaluated models.

Without fine-tuning, SQUALL generally achieved higher spatial concordance with transcript-level reference biomarker expression than the spatial proteomics models at both the 224 × 224- and 448 × 448-pixel scales (**Figures S5B and S5C**). Across biomarkers shared between SQUALL and ROSIE, their biomarker-level concordance with the transcriptomic reference was positively correlated (Pearson’s r = 0.123 at the 224 × 224-pixel scale and r = 0.162 at the 448 × 448-pixel scale), consistent with the modest concordance previously reported between matched transcript and protein measurements.^85,86^ Correspondence between SQUALL and GigaTIME was lower (r = −0.122 and −0.112, respectively), and ROSIE and GigaTIME likewise showed limited correlation across their shared biomarkers (**Figure S5B**). Notably, pairwise comparisons involving GigaTIME were based on a smaller set of shared biomarkers compared with ROSIE. The limited model-to-model concordance may therefore partly reflect restricted biomarker overlap, as well as differences in the scale and composition of their respective pretraining datasets (**Table S40**). SQUALL showed consistent performance across most biomarker categories (**Figures S5D-S5E**; **Tables S42-S43**). Together, these results indicate that, for matched gene-protein targets evaluated against a transcript-level reference, SQUALL provides spatial biomarker predictions that are at least comparable to, and generally show higher concordance than, those of pretrained spatial proteomics models, while extending virtual profiling to a substantially broader set of genes beyond the limited biomarker panels supported by these models.

We next asked whether SQUALL could profile molecular programs at cohort scale directly from whole-slide image (WSI) histology and whether these virtual profiles could support prognostic evaluation. We predicted the spatial expression of 15,757 genes from WSIs and associated these virtual biomarkers with clinical outcomes (**Figure 2J**). In the TCGA-LIHC cohort, multivariable Cox regression adjusted for age and stage showed that immune-activation programs were associated with prolonged survival, whereas invasion-related β-catenin/TCF signaling, a key axis of canonical Wnt signaling, and angiogenesis programs were associated with increased risk (**Figure 2K**). We also tested whether the spatial organization of virtually profiled biomarkers carried prognostic information. Using publicly available tumor-region annotations (**Figure 2L**),^87^ SQUALL recovered the spatial patterns of cervical cancer-associated differentially expressed genes and transcription factor programs reported in a previous study^88^ in the TCGA-CESC WSIs (**Figure S5F**). Region-resolved expression predictions associated DNA double-strand break repair programs with elevated risk (**Figures 2M and 2N**) and immune response pathways, including germinal center formation and chemokine activity, with prolonged survival (**Figure 2N**).

Among these protective immune programs, T cell activation also emerged as a prominent signal in this cohort (**Figure 2N**). Although T cell infiltration is broadly associated with favorable outcomes across cancers, the spatial features linking CD8⁺ cytotoxic activity to prognosis in cervical cancer remain unclear.^89^ SQUALL localized both CD4⁺ and CD8⁺ T cell programs preferentially to tumor regions (**Figure S5G**, left), where both effector and exhausted CD8⁺ T cell states were enriched (**Figures S5G** right and **S5H**). Intratumoral CD8⁺ T cell signatures were significantly associated with improved survival (Hazard ratio = 0.44, 95% CI: 0.24-0.79; one-sided Wald *P* = 0.003, testing for a protective association [HR < 1]) and showed the strongest protective association compared with signatures computed across the whole tissue or non-tumor regions, as well as clinical variables including stage and age (**Figure 2O**). Long-term survivors showed prominent intratumoral CD8⁺ infiltration, whereas short-term survivors showed immune accumulation primarily at the tumor-stroma interface (**Figure 2P**). These results identify intratumoral CD8⁺ immune activity as a spatial determinant of prognosis in cervical cancer and illustrate how SQUALL can extend spatially resolved molecular insights from specialized spatial profiling assays to clinically acquired histopathology WSIs.

**Figure S4.**
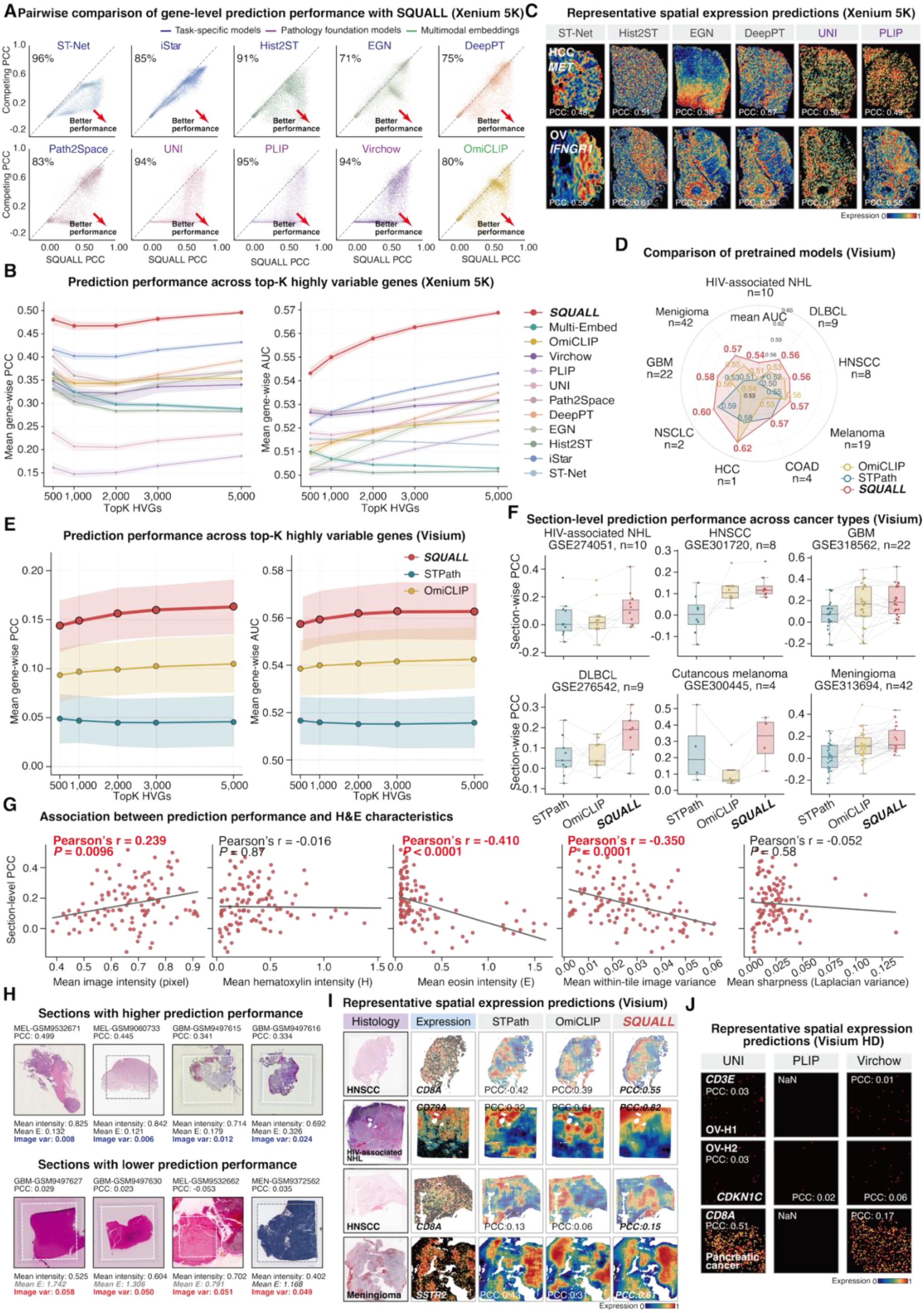
Comparison of SQUALL for virtual spatial gene expression profiling with ST prediction models. Related to Figure 2. **(A) –(C)** Comparative evaluation of virtual gene expression profiling. **(A)** Pairwise comparison of gene-level Pearson correlation coefficients (PCCs) between SQUALL (x-axis) and ST-Net, iStar, Hist2ST, EGN, DeepPT, Path2Space, UNI, PLIP, Virchow, or OmiCLIP (y-axes) across 4,421 shared genes. Percentages indicate the fraction of genes for which SQUALL achieved a higher PCC than the corresponding comparator. Related to Figure 2C. **(B)** Virtual gene expression profiling performance across the top highly variable genes in the comparative Xenium 5K evaluation. Curves show the mean gene-wise PCC (left) and AUC (right), and shaded areas indicate 95% confidence intervals. **(C)** Additional representative virtual biomarker predictions for the samples and genes shown in Figure 2E. Predictions generated by ST-Net, Hist2ST, EGN, DeepPT, UNI, and PLIP are shown; gene-level PCCs are indicated in each panel. HCC, hepatocellular carcinoma; OV, ovarian carcinoma. **(D) –(J)** Evaluation of pretrained models. **(D)** Radar plot summarizing virtual gene expression profiling performance on Visium sections across cancer types. Values represent the mean gene-wise AUC across 14,766 genes shared among the evaluated models. NHL, non-Hodgkin lymphoma; DLBCL, diffuse large B-cell lymphoma; HNSCC, head and neck squamous cell carcinoma; COAD, colon adenocarcinoma; HCC, hepatocellular carcinoma; NSCLC, non-small cell lung cancer; GBM, glioblastoma. Related to Figure 2G. **(E)** Performance of SQUALL, STPath, and OmiCLIP across the top highly variable genes in Visium sections. Curves show the mean gene-wise PCC (left) and AUC (right), and shaded areas indicate 95% confidence intervals. **(F)** Section-level comparison of virtual gene expression profiling performance across cancer types. Section-level PCCs were calculated by averaging gene-wise PCCs within each section. Points represent individual sections, and lines connect results from the same section across models. Center lines indicate medians, boxes indicate interquartile ranges, and whiskers extend to 1.5× the interquartile range. **(G)** Associations between section-level prediction performance and H&E image characteristics. Image intensity, hematoxylin intensity, eosin intensity, image within-tile pixel variance, and sharpness were summarized across tiles within each section. Hematoxylin and eosin intensities were estimated using the Vahadane method for stain separation. Pearson’s correlation coefficients and corresponding *P* values are shown. **(H)** Representative Visium sections with higher (top) or lower (bottom) SQUALL prediction performance. The corresponding mean image intensity, eosin intensity, and image within-tile pixel variance are shown below each section. Dashed outlines indicate the ST sequencing regions. **(I)** Representative virtual biomarker predictions on Visium sections. From left to right, panels show the H&E-stained histology image, measured expression, and expression predicted by STPath, OmiCLIP, and SQUALL. Rows show *CD8A* expression in HNSCC, *CD79A* expression in HIV-associated NHL, and *SSTR2* expression in meningioma. PCCs are indicated for each predicted expression profile. **(J)** Additional representative virtual biomarker predictions on Visium HD sections for the samples and genes shown in Figure 2I. Predictions generated by UNI, PLIP, and Virchow are shown for *CD3E* in ovarian cancer sample OV-H1, *CDKN1C* in ovarian cancer sample OV-H2, and *CD8A* in pancreatic cancer. PCCs are indicated for each prediction; NaN denotes cases in which PCC could not be calculated.

**Figure S5.**
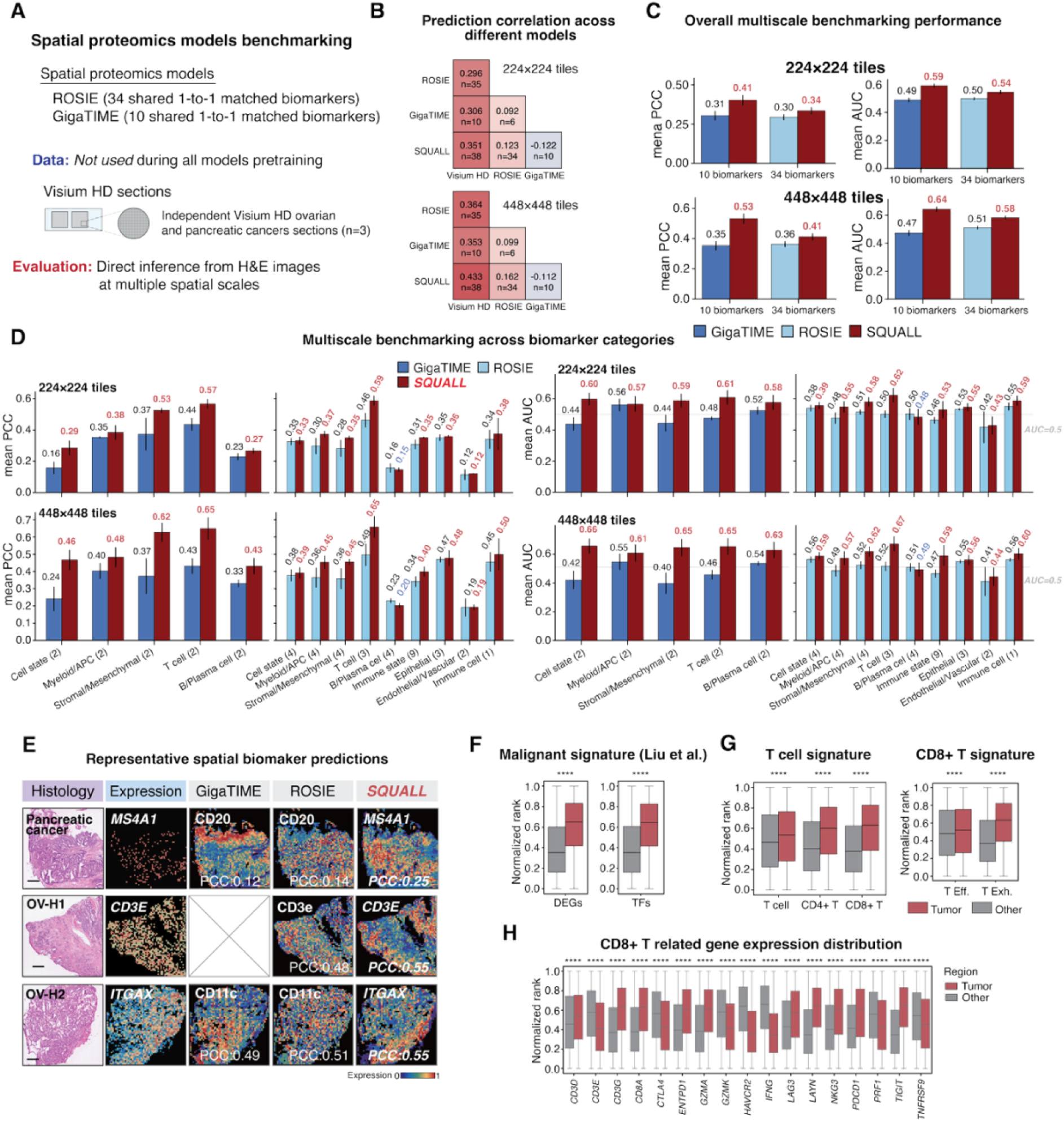
Cross-modality evaluation of SQUALL against pretrained spatial proteomics models and additional virtual biomarker profiling analyses. Related to Figure 2. **(A) –(E)** Evaluation of pretrained spatial proteomics models without additional fine-tuning. **(A)** Data and experimental design. ROSIE and GigaTIME were evaluated using 34 and 10 biomarkers with one-to-one matched gene and protein targets, respectively, that were also predicted by SQUALL. Given that datasets jointly profiling histology, spatial proteomics, and ST remain limited by heterogeneous experimental protocols and limited marker-panel sizes, Visium HD expression measurements were used as the reference for evaluating overlapping biomarkers under controlled conditions. Three independently collected Visium HD sections from ovarian and pancreatic cancer were not used during the pretraining of any evaluated model. All models generated virtual biomarker profiles directly from H&E images without additional fine-tuning. Performance was evaluated using 224 × 224- and 448 × 448-pixel tiles to assess spatial concordance across image scales. **(B)** Pairwise correlations between measured expression or virtual biomarker profiles generated by ROSIE, GigaTIME, and SQUALL at 224 × 224-pixel (top) and 448 × 448-pixel (bottom) resolutions. Values denote mean correlations across the corresponding shared biomarkers, and n indicates the number of biomarkers included in each comparison. **(C)** Overall multiscale evaluation. SQUALL was compared separately with GigaTIME across 10 shared biomarkers and with ROSIE across 34 shared biomarkers at 224 × 224- and 448 × 448-pixel resolutions. Bars show the mean biomarker-wise PCC and AUC; error bars indicate standard deviations. **(D)** Multiscale evaluation across shared biomarker categories. Performance was summarized within each biomarker category at 224 × 224-pixel (top) and 448 × 448-pixel (bottom) resolutions. Bars show the mean biomarker-wise PCC (left) and AUC (right), and error bars indicate standard deviations. Numbers in parentheses indicate the number of shared biomarkers in each category. APC, antigen-presenting cell. **(E)** Representative virtual biomarker predictions on Visium HD sections. From left to right, panels show the H&E-stained histology image, measured expression, and biomarker profiles predicted by GigaTIME, ROSIE, and SQUALL. Rows show the B cell marker *MS4A1*/CD20 in pancreatic cancer, the T cell marker *CD3E*/CD3e in ovarian cancer sample OV-H1, and the myeloid/antigen presenting cell marker *ITGAX*/CD11c in ovarian cancer sample OV-H2. PCCs are indicated for each prediction. The crossed panel indicates that the corresponding biomarker was unavailable for that model. **(F) –(H)** Additional virtual biomarker profiling analyses in the TCGA-CESC cohort. Predicted expression was normalized and ranked across tiles within each tissue section; higher ranks indicate higher predicted expression. Related to Figures 2O **and 2P**. **(F)** Comparison of SQUALL-predicted malignant program signatures between tumor and other tissue regions. Program signatures comprise cervical cancer-associated differentially expressed genes and transcription factors reported by Liu et al. DEGs, differentially expressed genes, TFs, transcription factors. **(G)** Comparison of predicted immune cell program signatures between tumor and other tissue regions. Left, pan T cell, CD4⁺ T cell, and CD8⁺ T cell program signatures. Right, CD8⁺ T cell effector and exhaustion program signatures. T Eff., T cell effector signature; T Exh., T cell exhaustion program signature. **(H)** Comparison of SQUALL-predicted expression ranks for individual CD8⁺ T cell-related genes between tumor and other tissue regions. For box plots in **(F)–(H)**, center lines indicate medians, boxes indicate the upper and lower quartiles, and whiskers extend to 1.5 times the interquartile range. Comparisons were performed using two-sided Mann– Whitney U tests. * *P* < 0.05, ** *P* < 0.01, *** *P* < 0.001, and **** *P* < 0.0001; n.s., not significant.

### SQUALL resolves spatial tumor microenvironment clusters and immune programs across cancers

Identifying spatially aware clusters is essential for interpreting spatial transcriptomics data and generating clinically relevant hypotheses related to tumor microenvironments.^90,91^ Existing approaches, however, often require dataset-specific optimization or prior knowledge of tissue context, limiting their applicability across diverse datasets.^92,93^ We therefore asked whether SQUALL’s frozen multimodal encoder could generate representations that capture spatially organized tumor microenvironment (TME) clusters with biological and clinical relevance (**Figures 3A and S6A**; **Methods**).

**Figure 3.**
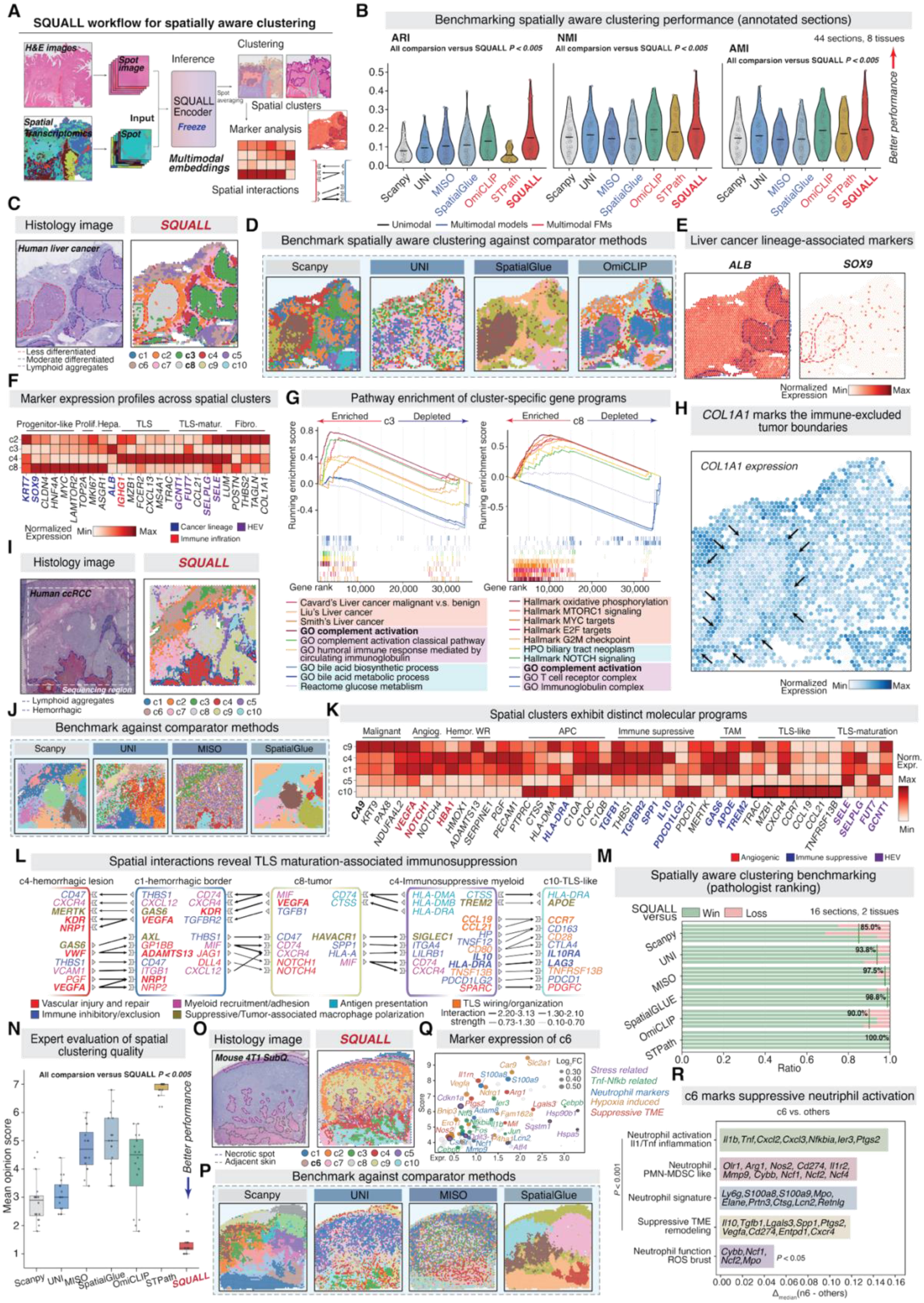
SQUALL characterizes spatial tumor microenvironment clusters and programs across cancer types. **(A)** Workflow for spatially aware clustering using SQUALL. Paired H&E images and ST profiles are processed by the frozen SQUALL encoder to generate multimodal spot-level embeddings. Clusters identified from these embeddings were used for downstream analysis. See also Figure S6A. **(B)** Evaluation of spatially aware clustering across 44 annotated sections representing eight tissues. Violin plots show the adjusted Rand index (ARI), normalized mutual information (NMI), and adjusted mutual information (AMI) for SQUALL and comparator methods. Higher values indicate greater agreement with reference annotations. Comparisons with SQUALL were performed using two-sided Wilcoxon signed-rank tests. See also Figure S6B. **(C)** Spatially aware clustering in a human liver cancer section. Left, H&E image with reference annotations of less-differentiated tumor, moderately differentiated tumor, and lymphoid aggregates. Right, spatial clusters identified by SQUALL. **(D)** Spatially aware clustering in the same liver cancer section using the unimodal and multimodal comparator methods Scanpy, UNI, SpatialGlue, and OmiCLIP. See also Figure S6G. **(E)** Spatial expression of liver cancer cluster-associated markers. Left, *ALB* expression marks the hepatocellular carcinoma cluster, with SQUALL-defined cluster c3 indicated by a blue dashed line. Right, *SOX9* expression marks a progenitor-like tumor cluster, with c8 indicated by a red dashed line. **(F)** Expression profiles of representative marker genes across major SQUALL-defined spatial clusters in the liver cancer section. Color intensity indicates normalized gene expression. Prolif., proliferative; Hepa., hepatocellular; Fibro., fibrotic; TLS, tertiary lymphoid structure; matur., maturation; HEV, high endothelial venule. **(G)** GSEA of cluster-specific molecular programs in the liver cancer section. Genes in cluster c3 (left) and cluster c8 (right) were compared with those in all other clusters. Positive running enrichment scores indicate programs enriched in the indicated cluster, whereas negative scores indicate depleted programs. **(H)** Spatial expression of *COL1A1* in the liver cancer section. Elevated *COL1A1* expression delineates the immune-excluded region surrounding cluster c8, as indicated by arrows. Color intensity represents normalized expression. **(I)** Spatially aware clustering in a human clear cell renal cell carcinoma (ccRCC) section. Left, H&E image with reference annotations highlighting lymphoid aggregates and hemorrhagic regions. Right, spatial clusters identified by SQUALL. **(J)** Spatially aware clustering in the same ccRCC section using Scanpy, UNI, SpatialGlue, and MISO. See also Figure S6J. **(K)** Expression profiles of representative marker genes across major SQUALL-defined spatial clusters in the ccRCC section. The clusters exhibit distinct malignant, angiogenic, hemorrhagic, antigen-presenting, immunosuppressive, tumor-associated macrophage, and tertiary lymphoid structure programs. Color intensity indicates normalized gene expression. Angiog., angiogenic; Hemor., hemorrhagic; WR, wound repair; APC, antigen-presenting cell; TAM, tumor-associated macrophage; TLS, tertiary lymphoid structure; HEV, high endothelial venule. **(L)** Inferred spatial ligand-receptor interactions among SQUALL-defined spatial clusters in the ccRCC section. Key significant interactions illustrate intercluster signaling associated with vascular injury and repair, immune inhibition and exclusion, myeloid recruitment and adhesion, suppressive tumor-associated macrophage polarization, antigen presentation, and TLS organization and maturation. Edge labels indicate the inferred interaction strength. **(M)** Pathologist evaluation of spatially aware clustering on an external dataset comprising 16 sections from two cohorts. Stacked bars show the proportions of cases in which SQUALL was judged to win or lose against comparator methods. The green vertical line indicates SQUALL’s mean win rate across comparisons. **(N)** Mean opinion scores for spatially aware clustering of SQUALL and comparator methods, based on rankings from five independent pathologists. Lower scores indicate better performance. Center lines indicate medians, boxes indicate interquartile ranges, and whiskers extend to 1.5× the interquartile range. Statistical significance was assessed using two-sided Wilcoxon signed-rank tests. See also Figure S6O. **(O)** Spatially aware clustering in a mouse 4T1 orthotopic tumor section. Left, H&E image with reference annotations highlighting necrotic regions and adjacent skin tissue. Right, spatial clusters derived by SQUALL. **(P)** Spatially aware clustering in the same mouse 4T1 tumor section using Scanpy, UNI, MISO, and SpatialGlue. See also Figure S6P. **(Q)** Differential expression analysis comparing SQUALL-defined cluster c6 with all other clusters in the mouse 4T1 tumor. The score represents the standardized Wilcoxon rank-sum statistic used to rank differentially expressed genes, with higher values indicating stronger evidence of differential expression. Point size represents the absolute log2 fold change. **(R)** Comparison of program scores between cluster c6 and all other clusters. Cluster c6 is characterized by neutrophil activation, IL-1- and TNF-associated inflammation, a polymorphonuclear myeloid-derived suppressor cell (MDSC)-like program, hypoxia, and suppressive tumor-microenvironment (TME) remodeling. Bars show the difference in mean signature scores between cluster c6 and the remaining clusters. Statistical significance was assessed using two-sided Mann–Whitney U tests.

We first performed a reference-based evaluation of spatially aware clustering using 44 sections from eight human tissues spanning tumor and normal contexts drawn from *histMol* (**Table S44**), including lung,^94^ soft tissue,^95^ breast,^96^ pancreas,^97^ liver,^97,98^ cervix,^99^ lymph node,^97^ and head and neck.^100^ Following previous evaluation settings,^101,102^ expert pathologists independently annotated the sections to provide reference labels for evaluation. We compared SQUALL with Leiden clustering of spatial transcriptomics spots (Scanpy),^103^ clustering of UNI-derived histology features,^23^ the multimodal methods SpatialGlue and MISO,^92,93^ and the pretrained H&E-ST foundation models OmiCLIP and STPath (**Table S45**).

Across datasets, SQUALL showed greater agreement with pathologist-annotated reference clusters than the comparator methods, as measured by the adjusted Rand index (ARI), normalized mutual information (NMI), and adjusted mutual information (AMI) (*P* < 0.005 for all comparisons, two-sided Wilcoxon signed-rank test; **Figure 3B**), with similarly robust performance under the unsupervised CHAOS metric (**Table S46**). Consistent performance trends were observed across individual cohorts (**Figure S6B**) and different cluster resolutions (**Figure S6C**), supporting robust spatially aware clustering performance across tumor contexts. We further investigated whether image characteristics and transcriptomic properties influenced clustering. Interestingly, compared with image features, transcriptomic properties such as increased housekeeping (Pearson’s r = -0.368, *P* = 0.01) and ribosomal gene abundance (Pearson’s r = -0.294, *P* = 0.05) showed negative correlations with ARI (**Figures S6D-S6E**), suggesting that poorly performing datasets may contain increased transcriptional background and reduced relative contributions of cluster-informative signals (**Figure S6F**). Even in these more challenging tissue sections, including the lung and head and neck cancer sections, SQUALL maintained robust spatially aware clustering performance compared with other approaches (**Figure S6B**).

Beyond these quantitative evaluations, SQUALL also resolved biologically meaningful TME clusters associated with distinct molecular programs. In a hepatocellular carcinoma section (**Figure 3C**),^97^ SQUALL uniquely separated tumor clusters and a lymphoid aggregate cluster within the same tissue compared with other methods (**Figures 3D and S6G**). The *ALB*-expressing hepatocellular carcinoma cluster c3 displayed a more differentiated phenotype, whereas cluster c8 expressed *SOX9* and *KRT7*, consistent with a less differentiated, *SOX9*⁺ progenitor-like tumor state (**Figure 3E**).^104^ Surrounding c8, cluster c2 expressed *POSTN* and *TAGLN* and exhibited a fibrotic stromal program, consistent with a cancer-associated fibroblast (CAF)-rich compartment (**Figures 3C and 3F**). SQUALL also identified a tertiary lymphoid structure (TLS) with features of maturation,^105^ including plasma cell markers (*MZB1* and *IGHG1*), B cell marker (*MS4A1*), T cell marker (*TRAC*), high endothelial venule programs (*GCNT1*, *FUT1*, *SELPLG*, and *SELE*), and the lymphoid-recruiting cytokine *CXCL13* (**Figures 3C and 3F**).

These identified tumor clusters also differed in immune infiltration. The hepatocellular tumor cluster c3 showed greater immune infiltration, whereas the *SOX9*⁺ progenitor-like cluster c8 showed stronger tumor proliferation programs (**Figures 3F and 3G**). Notably, c8 was surrounded by the c2 fibrotic stromal cluster, which was marked by higher *COL1A1* expression (**Figure 3H**), consistent with a stromal barrier to immune infiltration into the tumor parenchyma observed in multiple solid tumors.^106,107^ In TCGA-LIHC, patients with higher c3-derived gene program scores showed a significant positive correlation with CD8⁺ T cell infiltration (Spearman’s ρ = 0.42, *P* = 1.1 × 10⁻¹⁷), whereas patients with higher c8 program scores showed no significant association with CD8⁺ T cell infiltration (ρ = 0.02, *P* = 0.699), suggesting distinct immune associations between these tumor programs (**Figure S6H**). The c3:c8 program score ratio further stratified survival, with relative enrichment of the c8 program associated with worse outcomes (**Figure S6I**). Together, these findings suggest that distinct tumor lineage states are associated with different patterns of immune infiltration and clinical outcome, with TLS-related immune responses occurring within these lineage-dependent tumor contexts.

SQUALL also uncovered microenvironmental heterogeneity in a clear cell renal cell carcinoma (ccRCC) section.^108^ Within this section, SQUALL identified spatial clusters that aligned with annotated lymphoid aggregates and hemorrhagic regions (**Figure 3I**). Compared with existing approaches, SQUALL generated spatial cluster partitions with improved correspondence to the annotations (**Figures 3J and S6J**), recapitulating the annotated TME organization. These clusters represented distinct TMEs characterized by different spatial program expression and tumor states. The hemorrhagic cluster c4 showed elevated vascular-injury and angiogenesis programs, including *HBA1*, *HMOX1*, *PGF*, and *NOTCH4*, whereas the lymphoid aggregate cluster c10 expressed T cell, B cell, and plasma cell markers (**Figure 3K**). Unlike the mature TLS identified in liver cancer (**Figure 3C**), however, the ccRCC lymphoid aggregate cluster lacked the high endothelial venule programs *SELE*, *SELPLG*, *FUT1*, and *GCNT1* (**Figure 3K**).^105,109^ Gene set enrichment analysis showed the liver TLS was enriched for mature TLS-associated programs but not for the ccRCC lymphoid aggregate (**Figure S6K**), suggesting an immature TLS-like state.

Spatial cluster interaction analysis identified a signaling network linking tumor, hemorrhagic, and myeloid clusters that could impair TLS functions and maturation within this section (**Figure 3L**). Lymphoid-organizing *CCL19*/*CCL21*-*CCR7* interactions between myeloid (c5) and TLS-like (c10) clusters coexisted with inhibitory *IL10*-*IL10RA*, *TREM2*-*APOE*, and *HLA-DRA*-*LAG3* signaling, consistent with a suppressive tumor-associated macrophage (TAM)-like state and immunosuppressive microenvironment.^110,111^ This pattern is consistent with recent observations of immunosuppressive macrophages at tumor margins in TLS-containing ccRCC, where they colocalized with exhausted putative tumor-reactive T cells.^112^ Relative to the previously identified mature liver cancer TLS, the ccRCC lymphoid aggregate was enriched for angiogenesis, myeloid leukocyte activation, and macrophage-associated chemokine secretion and migration programs (**Figure S6K**), and also showed stronger exhausted T cell and suppressive myeloid programs together with weaker plasma cell-associated programs (*P* < 0.001 for all comparisons, two-sided Mann-Whitney U test; **Figure S6L**), further supporting a more suppressive and less mature immune state.

Of note, the surrounding tumor-hemorrhage clusters (c8-c1-c4) were enriched for vascular-injury and angiogenesis pathways, including VWF-ADAMTS13 and VEGFA-KDR/NRP1 interactions, suggesting that hemorrhagic clusters represent a vascular-remodeling region (**Figure 3L**). These clusters also exhibited signaling associated with suppressive TAM activation, including GAS6-MERTK/AXL and HAVCR1-SIGLEC1 interactions, suggesting that tumor-associated hemorrhage may establish a myeloid-enriched immunosuppressive microenvironment surrounding TLS-like structures.^112–114^ In the TCGA-KIRC cohort, the hemorrhage gene program signature derived from cluster c1 and c4 differentially expressed genes correlated positively with myeloid infiltration (**Figure S6M**), and higher hemorrhage program scores were associated with a shorter progression-free interval (**Figure S6N**), consistent with a more adverse clinical course. Together, these findings associate tumor-associated vascular injury and hemorrhage with myeloid organization, limited TLS maturation, and adverse outcomes in ccRCC.

We next performed a separate blinded reader evaluation by pathologists on previously unseen datasets of SQUALL pretraining to assess whether SQUALL-generated spatial clusters could facilitate clinically relevant hypothesis generation, as proposed in prior studies.^115^ We applied SQUALL to 16 newly collected sections, including mouse breast tumors and human thyroid cancers,^116,117^ derived from studies distinct from those used during SQUALL’s pretraining (**Table S44**). All pretrained models were evaluated under frozen inference settings, whereas other methods were applied separately to each dataset following the procedures described in their original studies. We assessed whether the spatial clusters identified by each method corresponded to interpretable tissue structures that could support downstream biological interpretation through blinded assessment by five board-certified pathologists. Head-to-head comparisons ranked SQUALL as the top-performing method among all evaluated approaches (**Figure 3M**). SQUALL also achieved the lowest (best) mean opinion score (**Figures 3N and S6O**), indicating that its spatially aware clustering performance was consistently preferred by expert evaluators (**Tables S47 and S48**). Notably, SQUALL outperformed STPath in both annotated section comparison and blinded reader evaluation, suggesting that multimodal representation learning can generate more interpretable spatial representations for biological discovery than pretraining objectives optimized toward a single generative prediction task.

In an external 4T1 mouse breast tumor section of the evaluation cohort, SQUALL delineated distinct spatial clusters (**Figures 3O, 3P, and S6P**) and identified cluster c6 as a necrosis-associated cluster enriched for hypoxia, stress-response, and TNF-NFκB programs (**Figure 3Q**). This cluster showed prominent neutrophil infiltration and polymorphonuclear myeloid-derived suppressor cell (PMN-MDSC)-like signatures (**Figure 3R**), linking tumor necrosis to a neutrophil-rich immunosuppressive microenvironment that may impair antitumor immunity.

**Figure S6.**
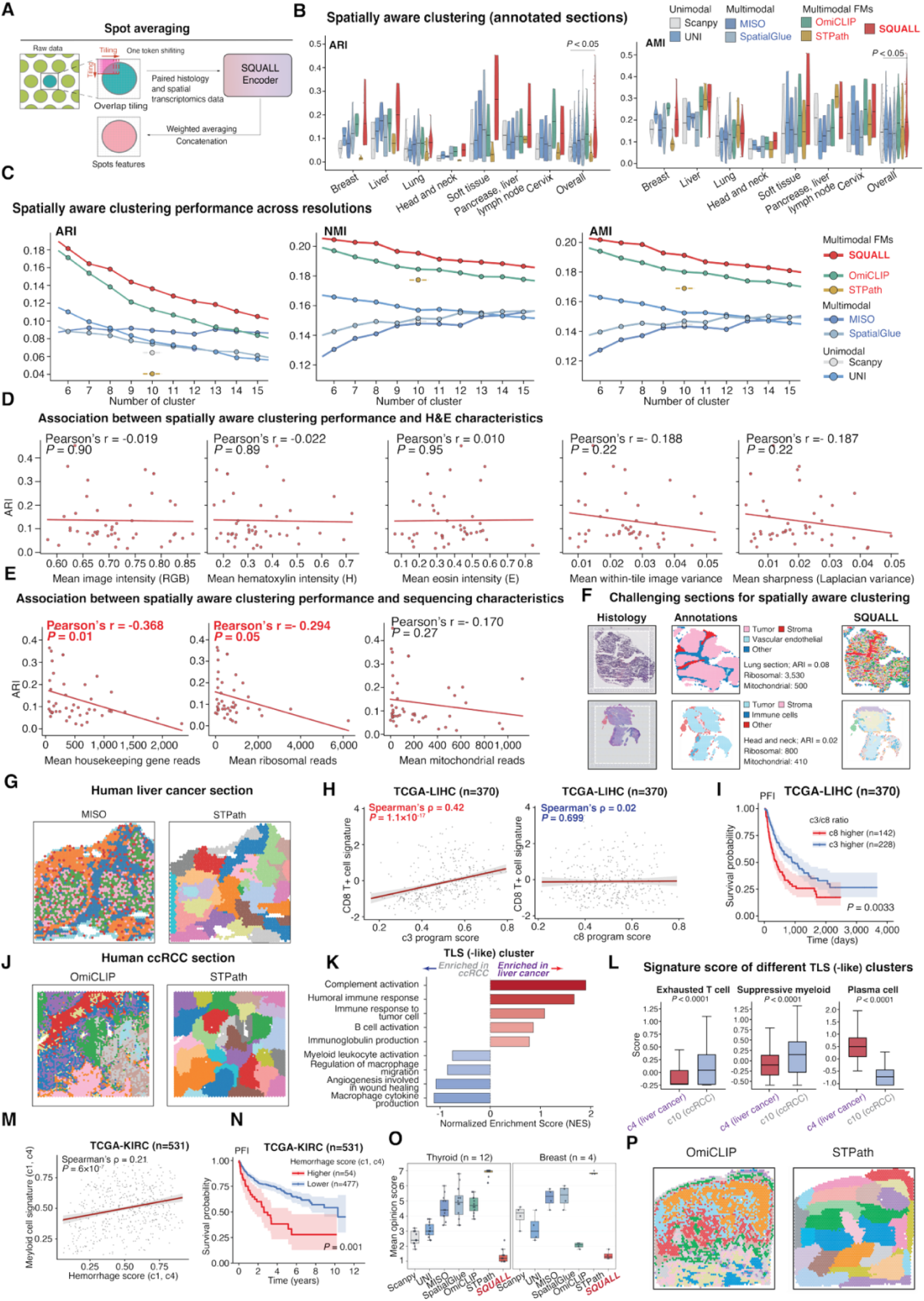
Evaluation SQUALL for spatially aware clustering and additional characterization of tumor-associated spatial clusters. Related to Figure 3. **(A)** Schematic of the spot-averaging procedure used to generate spot-level representations. Overlapping image tiles were processed by the SQUALL encoder, and tile-level representations overlapping each sequencing spot were aggregated by a weighted average of overlapping tile-token representations. **(B)** Evaluation of spatially aware clustering across 44 annotated sections representing eight tissues and seven cohorts. Violin plots show the adjusted Rand index (ARI), normalized mutual information (NMI), and adjusted mutual information (AMI) for SQUALL and comparator methods. Higher values indicate greater agreement with the reference annotations. Comparisons with SQUALL were performed using two-sided Wilcoxon signed-rank tests. Related to Figure 3B. **(C)** Evaluation of spatially aware clustering across resolutions on the annotated sections. For methods that do not explicitly use the number of clusters as a parameter, we report the corresponding metrics obtained using their default settings. **(D)** Associations between section-level spatially aware clustering performance and H&E image characteristics. Mean image intensity, hematoxylin intensity, eosin intensity, within-tile pixel variance, and image sharpness measured by the variance of the Laplacian were summarized across tiles within each section. Hematoxylin and eosin intensities were estimated using the Vahadane method for stain separation. Pearson’s correlation coefficients and corresponding *P* values are shown. **(E)** Associations between section-level spatially aware clustering performance and ST sequencing characteristics. Housekeeping-gene, ribosomal-gene, and mitochondrial-gene read counts were summarized across spots within each section. Pearson’s correlation coefficients and corresponding *P* values are shown. **(F)** Representative sections with low spatially aware clustering performance. From left to right, panels show the H&E-stained histology image, reference pathological annotations, and spatial clusters identified by SQUALL. ARI values and section-level ribosomal- and mitochondrial-gene read counts are indicated. **(G)** Spatial clusters identified by MISO and STPath in the human liver cancer section shown in Figure 3C. **(H)** Associations of the SQUALL-defined cluster c3 (left) and c8 (right) gene expression program signatures with the CD8⁺ T cell signature in the TCGA-LIHC cohort (n = 370). The c3 signature was positively associated with the CD8⁺ T cell signature, whereas no significant association was observed for the c8 signature. Spearman’s correlation coefficients and corresponding *P* values are shown. **(I)** Survival analysis in the TCGA-LIHC cohort stratified according to the relative dominance of the c3 or c8 cluster program signature. Patients were grouped using the c3-to-c8 program signature score ratio. Shaded areas indicate 95% confidence intervals, and the log-rank *P* value is shown. PFI, progression-free interval. **(J)** Spatially aware clustering identified by OmiCLIP and STPath in the human clear cell renal cell carcinoma (ccRCC) section shown in Figure 3I. **(K)** Gene set enrichment analysis comparing the tertiary lymphoid structure-like clusters c4 in liver cancer and c10 in ccRCC. Bars show normalized enrichment scores for immune and stromal functional programs. Positive values indicate enrichment in the liver cancer cluster, whereas negative values indicate enrichment in the ccRCC cluster. NES, normalized enrichment score; TLS, tertiary lymphoid structure. **(L)** Comparison of immune-related functional state program signature scores between liver cancer cluster c4 and ccRCC cluster c10. Exhausted T cell, suppressive myeloid cell, and plasma cell signature scores are shown. Center lines indicate medians, boxes indicate the upper and lower quartiles, and whiskers extend to 1.5 times the interquartile range. Comparisons were performed using two-sided Mann–Whitney U tests. **(M)** Association between the hemorrhage program signature and myeloid cell signature in the TCGA-KIRC cohort (n = 531). Program signatures were derived from the hemorrhage-associated ccRCC clusters c1 and c4. Spearman’s correlation coefficient and corresponding *P* value are shown. **(N)** Survival analysis in the TCGA-KIRC cohort stratified by the hemorrhage program signature score. Shaded areas indicate 95% confidence intervals, and the log-rank *P* value is shown. **(O)** Pathologist evaluation of spatially aware clustering in external breast and thyroid cancer cohorts. Box plots show mean opinion scores based on rankings from five independent pathologists; lower scores indicate better performance. Center lines indicate medians, boxes indicate the upper and lower quartiles, and whiskers extend to 1.5 times the interquartile range. Comparisons were performed using two-sided Wilcoxon signed-rank tests. MOS, mean opinion score. **(P)** Spatial clusters identified by OmiCLIP and STPath in the mouse 4T1 breast tumor section shown in Figure 3O.

### SQUALL reconstructs a progression-associated spatial molecular trajectory in breast cancer

Having shown that SQUALL can resolve tumor heterogeneity, immune programs, and spatial tumor microenvironment clusters, we next investigated whether it could capture progressive changes in tumor states associated with disease progression. Breast cancer accounts for ∼32% of newly diagnosed malignancies in women and remains the second leading cause of cancer-related mortality among females.^118^ Histopathologically, breast tumors exhibit substantial morphological and molecular heterogeneity, ranging from pre-invasive lesions such as ductal carcinoma in situ (DCIS) to invasive subtypes including invasive ductal carcinoma (IDC), invasive lobular carcinoma (ILC), and aggressive variants such as metaplastic breast cancer (MpBC).^119,120^ Despite accumulating evidence suggesting that tumor progression occurs along a continuum rather than through discrete states,^3,13^ the molecular programs accompanying these transitions remain incompletely characterized. We therefore used SQUALL joint histology-spatial transcriptomics embeddings to reconstruct a progression-associated molecular trajectory and characterize the accompanying transcriptional programs (**Figure 4A**).

**Figure 4.**
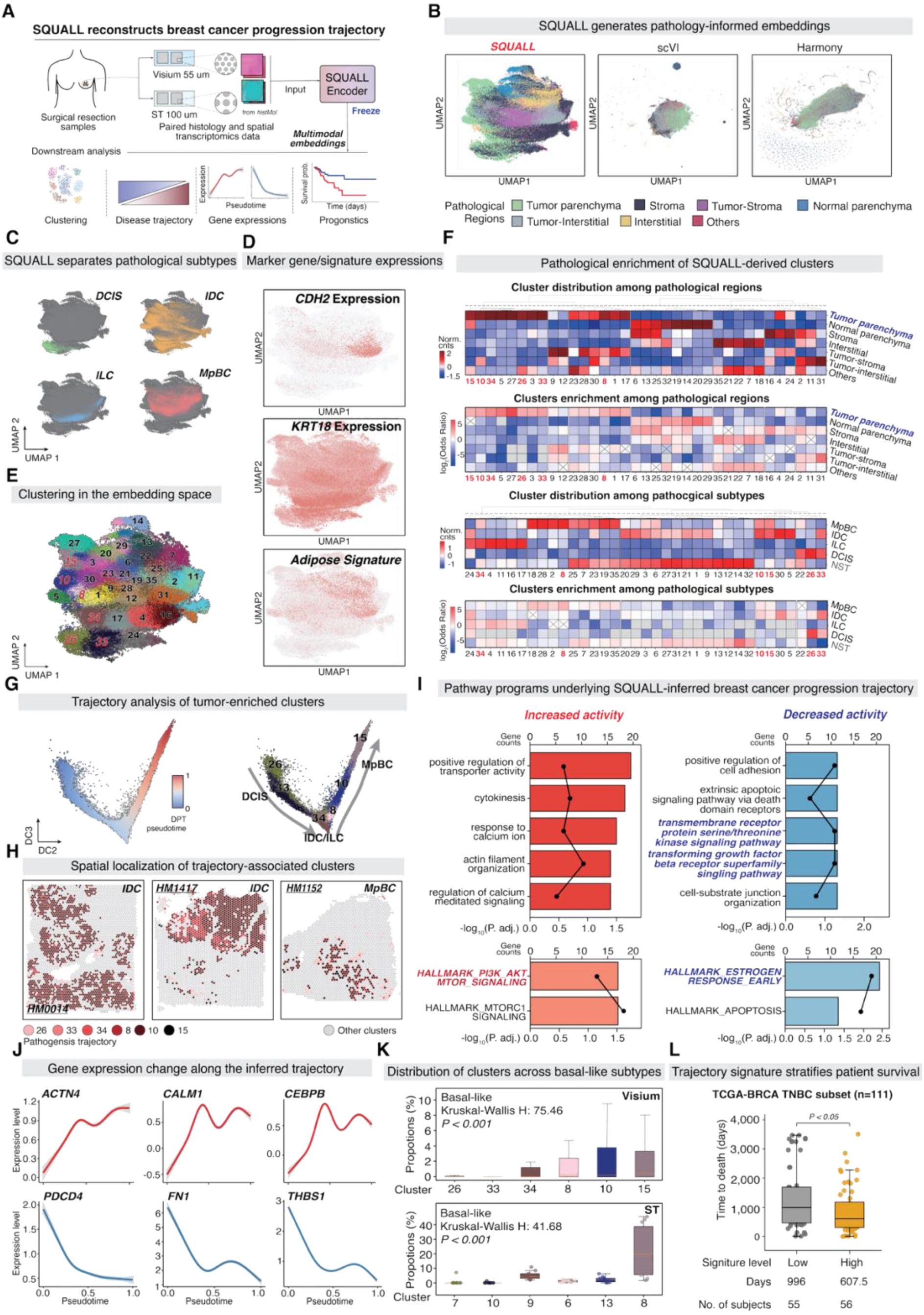
SQUALL infers spatial molecular trajectory associated with breast cancer progression. **(A)** Workflow for modeling breast cancer progression using SQUALL. Paired histology and ST data generated using Visium (55 μm spots) and spatial transcriptomics platforms (ST; 100 μm spots) are processed by the frozen SQUALL encoder to generate multimodal spot-level embeddings. These embeddings support downstream clustering, trajectory inference, gene expression analysis, and prognostic assessment. **(B)** UMAP visualization of spot-level embeddings from breast cancer sections generated by SQUALL, scVI, and Harmony. Spots are colored according to pathologist-annotated tissue regions. **(C)** UMAP visualization of SQUALL-derived spot embeddings highlighting pathological breast cancer subtypes. DCIS, ductal carcinoma in situ; IDC, invasive ductal carcinoma; ILC, invasive lobular carcinoma; MpBC, metaplastic breast cancer. See also Figure S7E. **(D)** Expression of representative marker genes and signatures projected onto the SQUALL embedding space. *CDH2* marks tumor-interstitial regions, *KRT18* marks invasive lobular carcinoma, and the adipose signature marks interstitial adipose tissue. **(E)** Unsupervised clustering of SQUALL-derived spot embeddings, with spots colored by cluster identity. **(F)** Distribution and enrichment of SQUALL-defined clusters across pathologist-annotated tissue regions (top) and breast cancer subtypes (bottom). The upper heatmap in each pair shows the normalized abundance of spots, whereas the lower heatmap shows enrichment quantified as the log2 odds ratio. Clusters enriched in tumor parenchyma and across pathological subtypes are highlighted. Enrichment was assessed using one-versus-all two-sided Fisher’s exact tests. Crosses indicate associations with a false discovery rate (FDR) greater than 0.1. **(G)** Diffusion map analysis of tumor-enriched clusters. Diffusion pseudotime (left) and cluster identity (right) are shown along the trajectory, respectively. The trajectory delineates a continuous progression from DCIS through invasive carcinoma (IC; including IDC and ILC) to MpBC. Arrows indicate the inferred direction of progression, and each point represents one ST spot. DPT, diffusion pseudotime. **(H)** Spatial localization of trajectory-associated tumor clusters in representative breast cancer sections. Colored spots indicate clusters positioned along the inferred progression trajectory, whereas gray spots represent all other clusters. **(I)** Functional programs associated with breast cancer progression. Over-representation analysis was performed separately for genes positively correlated with pseudotime, representing increased activity (left), and genes negatively correlated with pseudotime, representing decreased activity (right). Bars indicate −log10-adjusted *P* values, and points indicate the number of genes enriched in each program. **(J)** Expression dynamics of representative genes along the inferred progression trajectory. *ACTN4*, *CALM1*, and *CEBPB* are positively correlated with pseudotime, whereas *PDCD4*, *FN1*, and *THBS1* are negatively correlated. Lines indicate expression trends, and shaded regions represent 2.5th–97.5th percentile interval of corresponding gene expression. **(K)** Distribution of trajectory-associated tumor clusters across basal-like breast cancers in the Visium (top) and ST (bottom) datasets. Cluster proportions exhibit progressive enrichment patterns consistent with their ordering along the inferred trajectory. Statistical significance was assessed using two-sided Kruskal-Wallis tests. See also Figures S7F–S7J. **(L)** Distribution of observed time to death (days) between low- and high-signature groups in the triple-negative breast cancer subset of TCGA-BRCA (n = 111). Patients were stratified according to their trajectory-derived gene expression signature scores. Statistical significance was assessed using a two-sided Mann–Whitney U test. For the box plots in (**K)** and **(L)**, center lines denote medians, boxes indicate the interquartile range, and whiskers extend to 1.5 times the interquartile range.

To this end, we assembled a spatial cohort comprising 230,184 spots from 111 Visium sections across 70 patients spanning five studies in *histMol* (of 55 μm resolution per spot; **Figure S7A**; **Table S49**). Spots were encoded using SQUALL to generate multimodal embeddings integrating histological morphology and spatial gene expression. Matched H&E sections were also independently annotated by experienced pathologists using a consensus hierarchical framework for evaluation (**Figures S7B and S7C**; **Tables S50 and S51**).

To evaluate whether multimodal integration improved spatial representation, we compared SQUALL embeddings with alternative integration frameworks based on spatial transcriptomics alone, including Harmony and scVI.^121,122^ These methods were applied to the same cohorts using their standard dataset-specific workflows. SQUALL embeddings showed greater correspondence with histopathological structure and spatial context than the alternative representations, resulting in improved separation of annotated tissue compartments. In UMAP space, spot embeddings segregated into major histological compartments, including tumor parenchyma, stromal regions, tumor-stroma interfaces, normal parenchyma, interstitial regions, and tumor-interstitial interfaces (**Figure 4B**). The embeddings also separated breast cancer subtypes, including DCIS, IDC, ILC, and MpBC (**Figure 4C**).

Corresponding gene expression patterns supported the separations. *CDH2*, which encodes N-cadherin, is a cell adhesion molecule associated with invasive tumor migration through the interstitial extracellular matrix,^123^ corresponding to the tumor-interstitial region (**Figure 4D**, top). *KRT18* (encoding cytokeratin 18) is a luminal epithelial marker and is broadly expressed in ILC,^124^ consistent with their luminal epithelial phenotype (**Figure 4D**, middle). An adipose signature was preferentially expressed in interstitial regions (**Figure 4D**, bottom). These results demonstrate that SQUALL-derived embeddings capture both tissue-microenvironment organization and patient-level pathological subtype heterogeneity in breast cancer tissues.

Unsupervised clustering of the embeddings identified 35 recurrent molecular clusters spanning different pathological subtypes (**Figure 4E**). Enrichment analysis revealed strong associations between clusters and specific histological compartments or tumor subtypes (**Figure 4F**). These clusters corresponded closely to histology image features (**Figures S7D** and **S7E**), with tumor-enriched clusters displaying hyperchromatic nuclei and high tumor cell density, stromal clusters exhibiting intermediate morphology, and interstitial clusters enriched for adipocyte-associated programs and displaying lighter staining intensity. These findings indicate that SQUALL-derived molecular program states align with pathological tissue architecture.

To test whether these clusters reflected disease progression, we performed diffusion pseudotime analysis on tumor-enriched clusters shared across pathological subtypes (**Figures 4F and 4G**).^125^ This analysis revealed a continuous transcriptional state space spanning DCIS, invasive carcinomas (IDC and ILC), and more aggressive MpBC-like states (**Figure 4G**). Individual sections assigned to a single pathological subtype often contained multiple SQUALL-derived clusters (**Figure 4H**), suggesting substantial spatial heterogeneity within conventionally annotated lesions.

Genes that increased with pseudotime were enriched for actin cytoskeleton organization and calcium signaling, consistent with cytoskeletal remodeling and increased cell motility during progressive invasion (**Figure 4I**, left).^126,127^ These genes included *ACTN4*, an actin-binding protein involved in cytoskeletal reorganization and migration; *CALM1*, a mediator of calcium-dependent signaling; and *CEBPB*, a transcription factor implicated in tumor progression and inflammatory signaling (**Figure 4J**, top).

Conversely, genes that decreased with pseudotime were enriched for cell adhesion and apoptosis pathways, consistent with progressive loss of epithelial adhesion and attenuation of tumor-intrinsic apoptotic programs from DCIS through invasive carcinomas to MpBC (**Figure 4I**, right). These genes included *PDCD4*, a tumor suppressor involved in apoptosis; *FN1*, an extracellular-matrix component involved in adhesion; and *THBS1*, a matricellular protein linked to TGF-β signaling and microenvironmental remodeling (**Figure 4J**, bottom). TGF-β receptor signaling also declined, indicating progressive attenuation of tumor-intrinsic TGF-β transcriptional programs along the inferred invasive progression trajectory.

Of note, estrogen-response pathways likewise declined along invasion pseudotime (**Figure 4I**, right), consistent with a shift from hormone-dependent programs toward estrogen-independent growth. This shift was accompanied by increased PI3K/AKT/mTOR signaling, an oncogenic program frequently associated with basal-like and triple-negative breast cancers.^128^ The convergence of reduced estrogen signaling and increased PI3K/AKT/mTOR activity subsequently prompted us to examine the relationship between intrinsic molecular subtype and invasive progression. We assigned molecular subtypes using a PAM50 classifier on section-level pseudobulk profiles (**Methods**).^129^ The resulting assignments agreed strongly with curated patient metadata (Cohen’s κ = 0.68, *P* = 10^-6^, two-sided permutation test; **Figure S7F**). Within basal-like tumors, trajectory clusters were progressively enriched along pseudotime (*P* < 0.001, Kruskal-Wallis test; **Figure 4K**, top), linking basal-like transcriptional programs to the inferred invasive progression trajectory. We assessed the reproducibility of this trajectory in a separate, lower-resolution cohort drawn from *histMol*, comprising 37,761 spots from 87 sections across 33 patients (100 μm; **Figure S7G**; **Table S49**). SQUALL again resolved the major pathological compartments (**Figures S7H-S7J**) and recapitulated the progressive enrichment of trajectory clusters along pseudotime (Kruskal-Wallis test, *P* < 0.001; **Figure 4K**, bottom).

Finally, gene program signatures derived from pseudotime-associated programs stratified patients in the TCGA-BRCA cohort into prognostically distinct groups (**Figure S7K**). Within triple-negative breast cancer (TNBC), stronger expression of the SQUALL-derived invasion program was associated with shorter survival (**Figure 4L**), supporting the clinical relevance of the inferred progression trajectory-associated gene expression programs.

**Figure S7.**
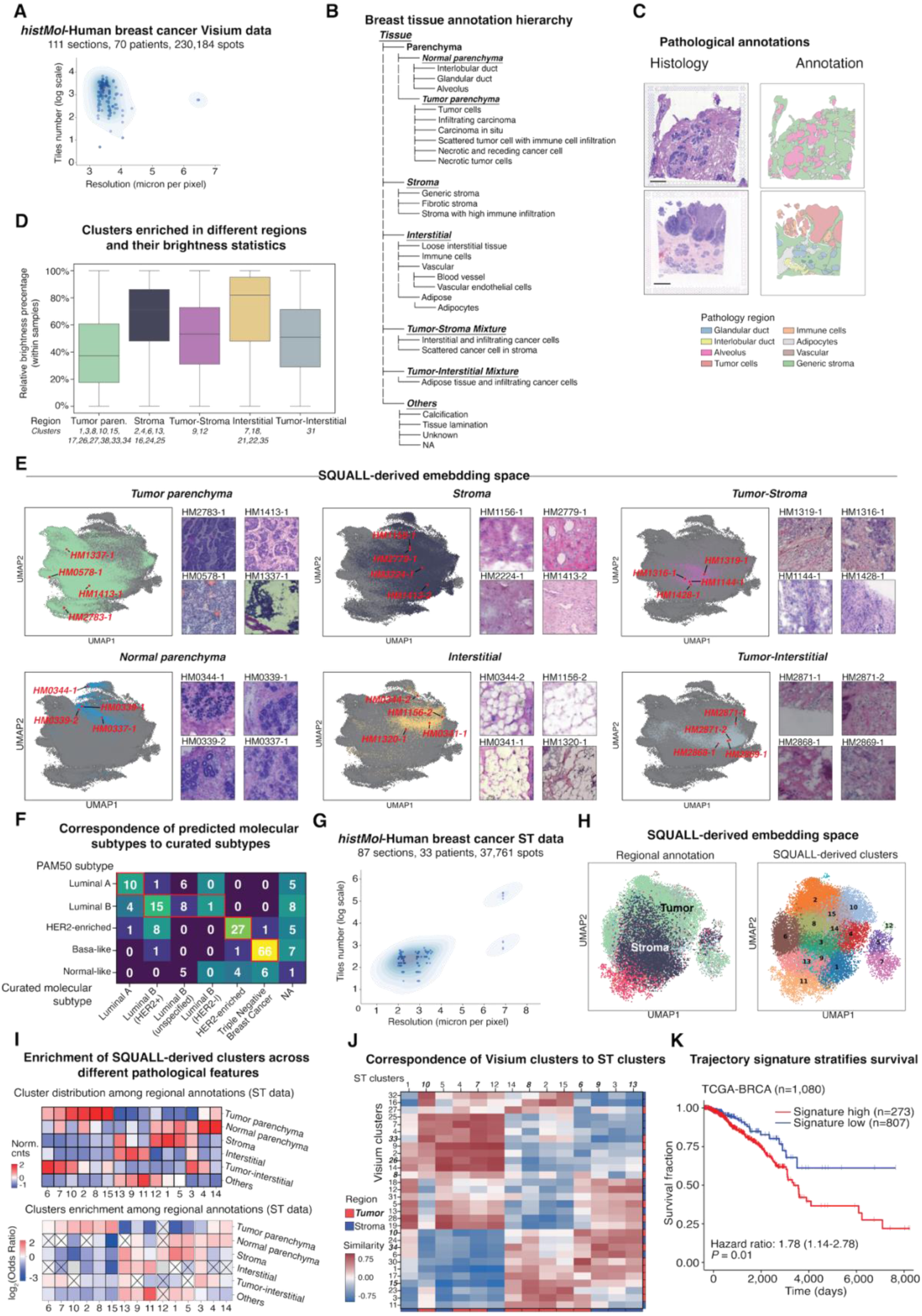
Additional characterization of SQUALL-inferred progression-associated trajectory in breast cancer from multimodal embeddings. Related to Figure 4. **(A)** Distribution of histology image resolution and the number of tiles generated per section in the *histMol* breast cancer Visium dataset, comprising 111 sections totaling 230,184 sequencing spots. Tile numbers are shown on a logarithmic scale. **(B)** Hierarchy of pathological annotations for breast tissues, established by consensus among pathologists and organized according to breast anatomical and pathological structures. Unknown denotes regions that could not be assigned to a specific class; NA denotes regions with missing annotations. **(C)** Representative breast cancer sections with pathological annotations. H&E-stained histology images are shown on the left, and corresponding non-overlapping annotation polygons are shown on the right. Scale bar, 1 mm. **(D)** Relative histology image brightness across SQUALL-derived clusters enriched for different pathological regions. Brightness was normalized within each tissue section. Center lines indicate medians, boxes indicate the upper and lower quartiles, and whiskers extend to 1.5 times the interquartile range. **(E)** UMAP visualization of the SQUALL-derived spot-level embedding space. Spots associated with tumor parenchyma, stroma, tumor-stroma, normal parenchyma, interstitial, and tumor-interstitial regions are highlighted separately. Representative histology image patches corresponding to selected positions in the embedding space are shown alongside each UMAP. **(F)** Correspondence between curated molecular subtypes and PAM50 subtypes inferred from ST data. Curated subtypes were obtained from the original publications and reviewed by pathologists, whereas PAM50 subtypes were inferred from pseudobulk expression profiles generated for each tissue section. Sections with valid subtype assignment were shown. Values indicate the number of sections assigned to each subtype combination. **(G)** Distribution of histology image resolution and the number of tiles generated per section in the *histMol* breast cancer ST dataset, comprising 87 sections totaling 37,761 sequencing spots. Tile numbers are shown on a logarithmic scale. **(H)** UMAP visualization of the SQUALL-derived embedding space for ST tissue sections, colored by pathological regional annotations (left) or K-means-derived clusters (right). **(I)** Distribution and enrichment of SQUALL-derived clusters across pathological regions in the ST dataset. Top, normalized cluster counts within each pathological annotation. Bottom, cluster enrichment assessed by one-versus-all odds ratios using two-sided Fisher’s exact tests. Crosses indicate associations with a false discovery rate greater than 0.1. **(J)** Correspondence between SQUALL-derived clusters in the Visium and ST datasets. Similarity was defined as the median cosine similarity between each ST cluster and Visium cluster, with the highest-scoring Visium cluster assigned as the corresponding counterpart. Annotation bars indicate clusters associated with tumor or stromal regions. **(K)** Kaplan–Meier survival analysis in the TCGA-BRCA (n = 1,080) cohort using a prognostic program signature constructed from genes most strongly associated with the SQUALL-derived tumor trajectory. Patients were stratified into signature-high and signature-low groups. The hazard ratio and its 95% confidence interval are shown, and statistical significance was assessed using a log-rank test.

### SQUALL identifies a recurrence-associated tumor-adaptive, immune-excluded spatial region in ovarian tumor

We next asked whether SQUALL could identify spatial regions associated with clinical outcomes. In advanced ovarian cancer, recurrence remains frequent after initial therapy, with more than 70% of patients experiencing relapse, yet the spatial microenvironments associated with recurrence remain poorly defined.^130^ We applied SQUALL to a Visium HD ovarian tumor cohort from *histMol* (**Table S52**),^27^ representing each 112 μm × 112 μm tissue tile as a multimodal embedding of its local microenvironment (**Figure 5A**). The embeddings organized tissue regions into a pathology-aware space that closely matched expert histopathological annotations (**Figures S8A-S8D**; **Table S22**).

**Figure 5.**
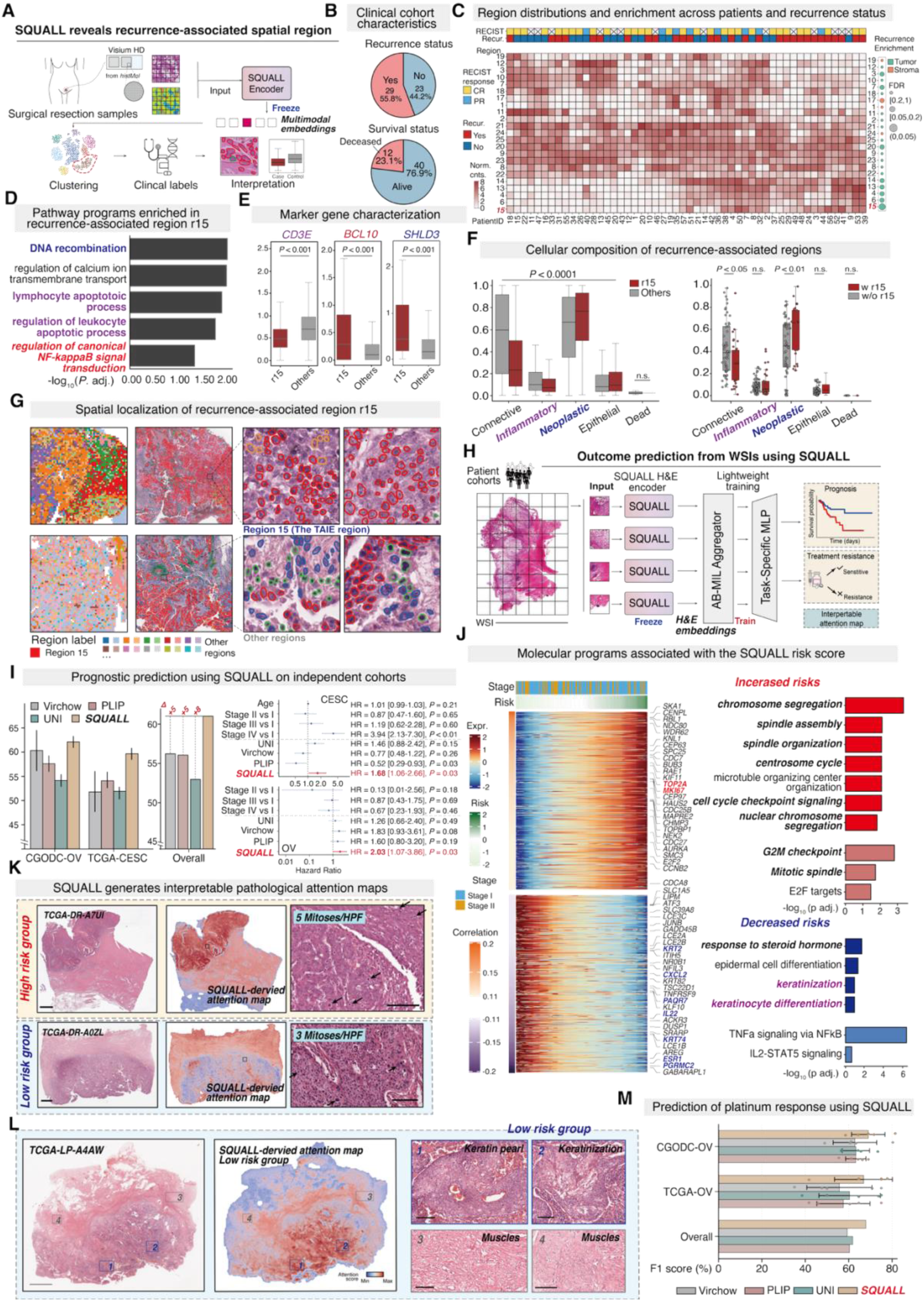
SQUALL facilitates clinically relevant feature discovery and outcome prediction. **(A)** Workflow for identifying recurrence-associated spatial regions in human ovarian tumor using SQUALL. Visium HD sections are processed by the frozen SQUALL encoder to generate multimodal tile-level embeddings. Clustering of these embeddings, followed by integration with clinical outcomes, enables the identification and pathological interpretation of recurrence-associated spatial regions. **(B)** Clinical characteristics of the Visium HD ovarian tumor cohort. Pie charts show the distributions of recurrence status (top) and survival status (bottom). **(C)** Identification of recurrence-associated spatial region 15 (r15). Patient-level RECIST response and recurrence status are indicated above the heatmap. The heatmap summarizes the distribution of spatial regions defined by unsupervised clustering across patients, based on SQUALL-derived tile-level embeddings. Annotations on the right indicate enrichment of individual regions by recurrence status and pathological category. Enrichment was assessed using two-sided Fisher’s exact tests with FDR correction. CR, complete response; PR, partial response; Recur., recurrence; crosses, missing clinical data. **(D)** Gene Ontology (GO) term enrichment analysis of genes upregulated in r15 relative to all other spatial regions. Bars indicate −log10-adjusted *P* values. **(E)** Expression of representative marker genes in r15 and all other spatial regions. Box plots show the expression levels of *CD3E*, *BCL10*, and *SHLD3*. Statistical significance was assessed using a two-sided Mann– Whitney U test. **(F)** Cellular composition of the recurrence-associated region. Left, comparison of connective, inflammatory, neoplastic, epithelial, and dead cell fractions between r15 and all other regions. Right, patient-level comparison of cellular composition between sections containing r15 and those without r15. Region r15 is characterized by an increased neoplastic-cell fraction and reduced inflammatory-cell infiltration. Whole-section cell segmentation and classification were performed using CellViT++. Statistical significance was assessed using a two-sided Mann–Whitney U test. n.s., not significant. **(G)** Spatial localization and cellular characterization of r15 in representative ovarian tumor sections. Left, SQUALL-defined spatial regions, with r15 highlighted in red. Middle, corresponding whole-section cell segmentation and classification generated using CellViT++. Right, representative high-magnification fields from r15 and other regions, with cells colored by predicted class. **(H)** Workflow for predicting survival and treatment resistance from cohort-level H&E-stained WSIs. Image tiles are processed by the frozen SQUALL H&E encoder, and the resulting embeddings are aggregated using attention-based multiple-instance learning (ABMIL). Lightweight task-specific networks are trained to predict prognosis or treatment resistance, while attention scores provide spatially interpretable prediction maps. **(I)** Survival prediction using SQUALL in independent cohorts. Left, concordance index (C-index) from fivefold cross-validation in the CGODC ovarian cancer (CGODC-OV) and TCGA cervical cancer (TCGA-CESC) cohorts, with overall performance shown across cohorts. SQUALL is compared with Virchow, PLIP, and UNI; values above the overall bars indicate SQUALL’s performance gain in percentage relative to each comparator. Error bars indicate ±1 standard deviation. Right, associations between clinical variables, model-derived risk scores, and survival. Hazard ratios were estimated using multivariable Cox proportional hazards models for TCGA-CESC and CGODC-OV. Points indicate hazard ratios, and horizontal bars indicate 95% confidence intervals. Statistical significance was assessed using two-sided Wald tests. **(J)** Molecular programs associated with SQUALL-derived risk scores in the TCGA-CESC cohort. Left, heatmaps showing top genes positively or negatively correlated with predicted risk as evaluated using corresponding bulk RNA-seq profiles, with patients ordered by increasing risk score and annotated by clinical stage. Right, GO term enrichment analysis of genes associated with increased or decreased predicted risk. Bars indicate −log10-adjusted *P* values. **(K)** Histopathological features associated with SQUALL-predicted survival risk in representative TCGA-CESC cases. Rows show examples from the high- and low-risk groups. From left to right, panels show H&E-stained WSIs, SQUALL-derived attention maps, and representative high-attention regions used to quantify mitotic figures. Arrows indicate mitotic figures. Scale bars, 2.5 mm for WSIs and 100 μm for magnified regions. See also Figure S9C–S9F. **(L)** Pathological interpretation of a representative low-risk TCGA-CESC case. From left to right, panels show the H&E-stained WSI, the corresponding SQUALL-derived attention map, and regions with distinct attention scores. High-attention regions contain differentiated squamous features, including keratin pearls and keratinization, whereas low-attention regions contain muscle tissue. Scale bars, 2.5 mm for WSIs and 100 μm for magnified regions. **(M)** Prediction of platinum-based chemotherapy resistance in two independent ovarian cancer cohorts. F1 scores from fivefold cross-validation are shown. SQUALL is compared with Virchow, PLIP, and UNI. Bars and error bars represent the mean ±1 standard deviation, respectively. For the box plots in **(E)** and **(F)**, center lines denote medians, boxes indicate the interquartile range, and whiskers extend to 1.5 times the interquartile range.

Hierarchical clustering identified recurrent spatial regions across tumors (**Figures 5B and 5C**), among which region r15 was enriched in tumors from patients who experienced recurrence (FDR < 0.05). Genes upregulated in r15 were enriched for NF-κB signaling, DNA recombination, calcium transport, and lymphocyte-apoptosis program (**Figure 5D**). Given the context-dependent roles of NF-κB signaling in promoting tumor survival while impairing anti-tumor immune responses,^131,132^ these observations linked NF-κB-associated programs to recurrence and were consistent with combined tumor-promoting and immunosuppressive effects. Consistent with this hypothesis, this region showed reduced *CD3E* expression, indicating lower T cell abundance, together with elevated expression of *BCL10*, an activator of NF-κB signaling,^133^ and *SHLD3*, a component of the Shieldin complex involved in DNA double-strand break repair (**Figure 5E**). These features defined a microenvironment of tumor persistence and local immune suppression associated with ovarian tumor recurrence.

We termed r15 the tumor-adaptive, immune-excluded (TAIE) region. Joint cell segmentation and cell-type classification with CellViT++^134^ confirmed that TAIE regions contained fewer inflammatory cells and more neoplastic cells than other tissue regions (**Figure 5F**). At the patient level, tumors harboring TAIE regions had larger tumor-cell fractions and tumor-dominant compartments with minimal lymphocyte infiltration (**Figures 5F and 5G**). Thus, the TAIE region defines a recurrence-associated ovarian tumor microenvironment marked by tumor predominance, immune exclusion, and NF-κB-linked adaptive signaling.

**Figure S8.**
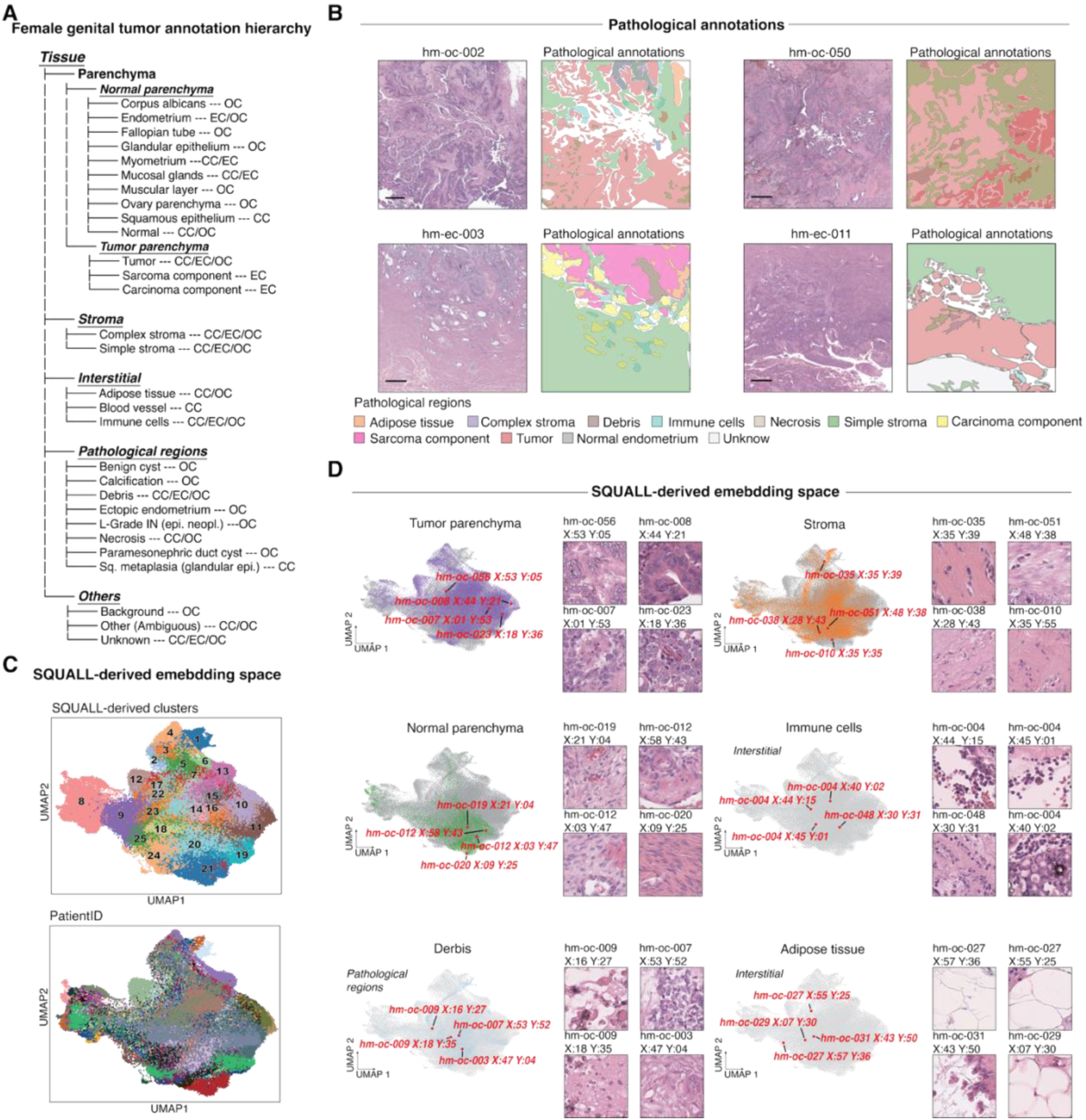
Additional characterization of SQUALL-identified recurrence-associated spatial region in ovarian cancer using multimodal embeddings. Related to Figures 5A–5G. **(A)** Hierarchical framework for pathological annotation of female genital tract tumors, established by consensus among three experienced pathologists. Annotations encompass normal and malignant parenchyma, stroma, interstitial components, disease-associated pathological regions, and other tissue features. OC, ovarian cancer; EC, endometrial carcinoma; CCEC, clear cell endometrial carcinoma; CC, cervical carcinoma; Sq, squamous. **(B)** Representative H&E-stained tissue regions and corresponding non-overlapping pathological annotation polygons. Colors indicate adipose tissue, complex stroma, debris, immune cells, necrosis, simple stroma, carcinoma, sarcoma, tumor, and normal endometrium. Scale bar, 500 µm. Only OC-related annotations and sections were used for recurrence analysis. **(C)** UMAP visualization of tile-level SQUALL embeddings, colored by SQUALL-derived cluster identity (top) or patient identity (bottom). **(D)** Distribution of pathological regions within the SQUALL-derived embedding space. Tiles associated with tumor parenchyma, stroma, normal parenchyma, immune cells, debris, and adipose tissue are highlighted separately. Representative H&E image tiles corresponding to selected positions in the embedding space are shown alongside each UMAP.

### SQUALL improves outcome prediction from whole-slide histology and reveals interpretable risk programs

Prognostic prediction remains challenging because of tumor heterogeneity, yet H&E-stained whole-slide images are routinely available and integral to patient care.^53^ Having shown that SQUALL virtually profiled spatial gene expression could be linked to patient survival (**Figures 2J-2P**), we next asked whether molecular information encoded by the SQUALL image encoder during multimodal pretraining could support end-to-end outcome prediction directly from histology, without explicitly reconstructing gene expression profiles.

To this end, we extracted tile-level image representations from WSIs, aggregated them using attention-based multiple-instance learning, and trained Cox proportional hazards models for survival prediction without fine-tuning the pretrained histology encoder (**Figure 5H**; **Table S53**). None of the patient data from the evaluated cohorts were used to pretrain SQUALL or, as reported, the comparator foundation models, thereby ensuring an independent evaluation. Although SQUALL was pretrained on a comparatively modest corpus of approximately 0.7 million paired histology-spatial transcriptomics tiles, compared with hundreds of millions to billions of images used to train UNI,^23^ Virchow,^62^ and PLIP (**Tables S54 and S55**),^61^ it achieved higher survival prediction performance across three cohorts comprising 770 patients (**Tables S56-S58**). In five-fold cross-validation at 20× magnification, SQUALL achieved a C-index of 0.61 across the gynecologic cancer cohorts (TCGA-CESC and our newly collected CGODC-OV cohort), compared with 0.53 for UNI, 0.56 for both PLIP and Virchow (**Figure 5I**, left), and also showed improved performance in the stomach adenocarcinoma (TCGA-STAD) cohort (**Table S60**). Subsequent multivariable Cox regression analysis showed that SQUALL-derived risk scores remained significantly associated with prognosis after adjustment for clinical variables in both cohorts, whereas the other evaluated pathology foundation models did not demonstrate consistent significant prognostic associations across cohorts. These findings suggest that SQUALL captures prognostically relevant information beyond these clinical variables and retains prognostic value under multivariable adjustment more consistently than the other evaluated pathology foundation models (**Figure 5I**, right).

To assess whether WSI characteristics influenced model performance, we additionally evaluated SQUALL on the TCGA-OV cohort. In contrast to the primary gynecologic cohorts, TCGA-OV was characterized by substantially larger tissue areas and greater staining variation (**Table S61**), and SQUALL showed reduced predictive performance in this setting, whereas Virchow performed comparatively well (C-index = 0.63). This advantage may reflect the greater robustness conferred by Virchow’s large-scale pathology pretraining (**Table S55**). Consistent with this observation, lower SQUALL performance across cohorts was associated with larger tissue areas and greater color variance (**Table S61**), suggesting that whole-slide scale and staining heterogeneity may contribute to cohort-dependent variation in performance.

In the CGODC-OV cohort, SQUALL attention maps preferentially localized to stromal regions in the predicted high-risk group, in contrast to those of competing models (**Figure S9A**). Notably, SQUALL recovered these prognostically relevant stromal features directly from histology images, consistent with stromal and extracellular matrix programs previously identified through molecular profiling studies in ovarian cancer.^135^ In the TCGA-CESC cohort, using model-specific optimal cutoffs determined from each model’s predicted risk scores by the same maximally selected rank statistics procedure,^136^ SQUALL was the only model that significantly stratified survival risk among patients with early-stage disease (stage I/II; hazard ratio = 2.42; two-sided Wald test for hazard ratio, P < 0.05; log-rank test for survival, P = 0.00877; **Figure S9B**), a clinically relevant setting in which prognostic discrimination remains limited.

To probe the underlying biology, we performed an exploratory correlation analysis between patient-level SQUALL-derived risk scores and bulk RNA-seq profiles in TCGA-CESC. Higher risk scores were associated with proliferation and mitotic programs, including *MKI67* and *TOP2A* (**Figure 5J**), consistent with increased cell-cycle activity in HPV-driven tumorigenesis. Conversely, genes negatively correlated with risk were enriched for epidermal differentiation and immune pathways, including TNF-α signaling through NF-κB and IL2-STAT5 signaling (**Figure 5J**), suggesting that poor outcomes accompany loss of differentiation and diminished immune surveillance. Using histology alone, SQUALL also recapitulated the favorable association between steroid-hormone response and prognosis previously identified by TCGA through RPPA-based stratification.^137^ Two board-certified pathologists confirmed the corresponding morphology: high-risk regions contained more mitotic figures (*P* < 0.05, two-sided Mann-Whitney U test; **Figure 5K**; **Figure S9C-F)**, whereas low-risk attention localized to keratin pearls and other keratinization features recognizable on histology images (**Figure 5L**).

Finally, we tested whether SQUALL captured treatment-relevant phenotypes by predicting resistance to platinum-based chemotherapy, a standard treatment for ovarian cancer,^130,138^ in two independent cohorts not used for pretraining of SQUALL and all comparators (**Tables S56 and S59**). Notably, despite its lower performance in survival prediction in the TCGA-OV cohort, SQUALL remained effective for predicting platinum resistance. Although overall accuracy was comparable across models, SQUALL consistently achieved higher F1 scores in both cohorts (**Figure 5M**; **Table S62**), indicating improved detection of treatment resistance under class imbalance, a common challenge of clinical datasets.^139^

**Figure S9.**
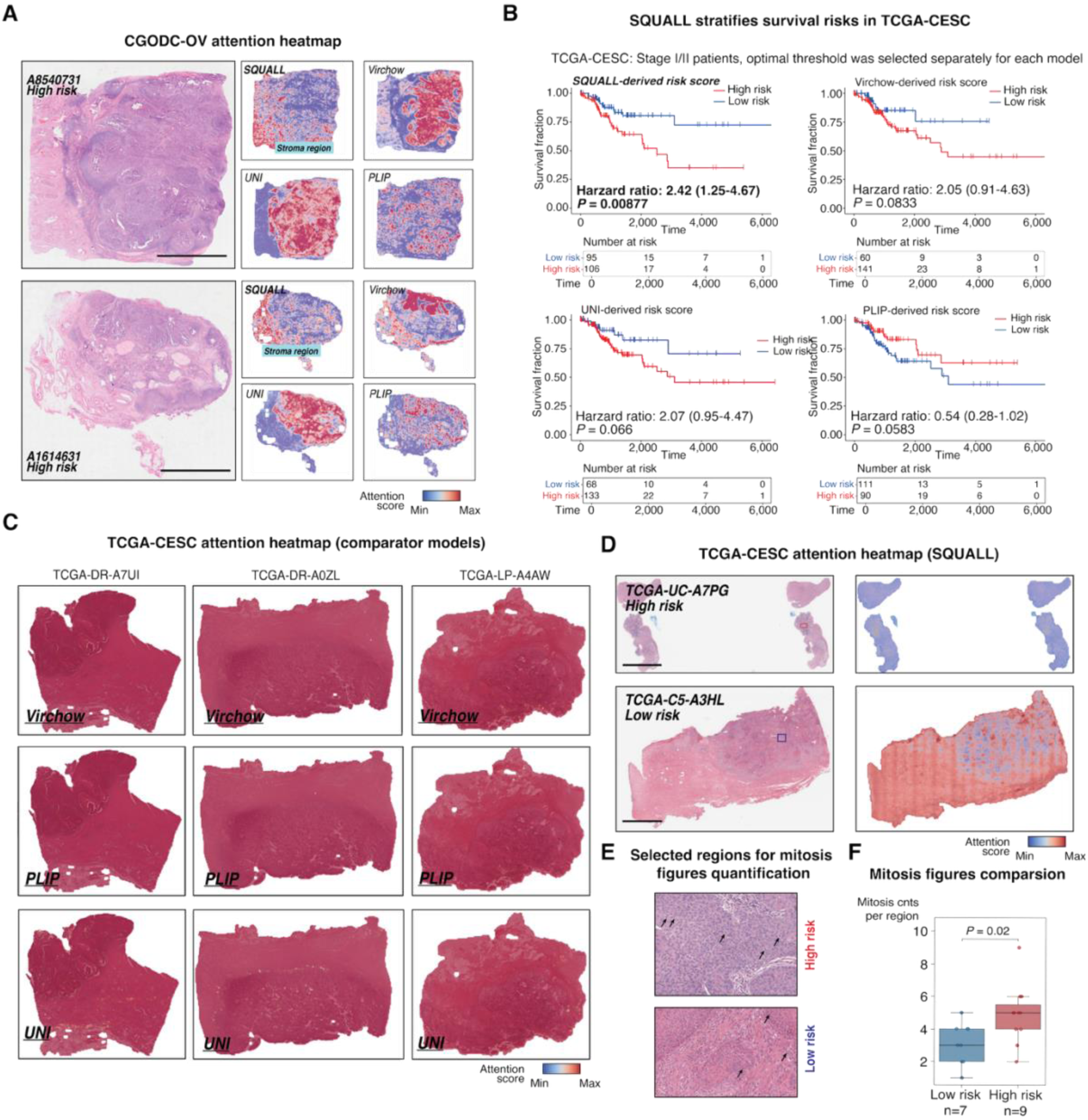
Additional evaluation of prognostic prediction from whole-slide histology images using SQUALL. Related to Figures 5H–5M. **(A)** Representative high- and low-risk ovarian cancer cases from the CGODC-OV cohort. H&E-stained whole-slide images are shown on the left, and attention heatmaps generated by SQUALL, Virchow, UNI, and PLIP are shown on the right. Color intensity represents the model-derived attention score. Scale bar, 5 mm. Related to Figure 5I. **(B)** Kaplan–Meier survival analysis of patients with stage I/II cervical cancer in the TCGA-CESC cohort, stratified using risk scores derived by SQUALL, Virchow, UNI, or PLIP. The optimal threshold for defining high- and low-risk groups was selected based on maximum separation setting separately for each model. Hazard ratios and their 95% confidence intervals are shown, and statistical significance was assessed using log-rank tests. Tables show the numbers of patients at risk. Related to Figure 5I. **(C)** Representative attention heatmaps generated by the comparator models Virchow, PLIP, and UNI for three TCGA-CESC whole-slide images. Color intensity represents the model-derived attention score. Related to Figure 5K. **(D)** Additional representative high- and low-risk TCGA-CESC cases. H&E-stained whole-slide images are shown on the left, and SQUALL-derived attention heatmaps are shown on the right. Color intensity represents the model-derived attention score. Scale bar, 5 mm. **(E)** Representative regions selected from SQUALL-derived attention maps for quantification of mitotic figures in the predicted high- and low-risk groups. Arrows indicate representative mitotic figures. Related to Figure 5K. **(F)** Comparison of mitotic-figure counts per selected region between the SQUALL-predicted high- and low-risk groups. Center lines indicate medians, boxes indicate the upper and lower quartiles, and whiskers extend to 1.5 times the interquartile range. Statistical significance was assessed using a two-sided Mann–Whitney U test. Related to Figure 5K.

## Discussion

A central challenge in cancer biology and pathology is that routine histopathology captures the architectural context through which tumors are diagnosed and interpreted, yet provides only indirect access to the molecular programs associated with tumor progression, immune organization, and therapeutic outcome. Our results suggest that tissue morphology contains informative correlates of these molecular states that can be made accessible through integrative training with spatial molecular measurements. To enable such learning at scale, we assembled *histMol*, comprising 1.93 billion paired histology-spatial transcriptomics spots or bins across diverse tissues, platforms, and spatial resolutions, and developed SQUALL as a multimodal foundation model linking tissue morphology with spatial molecular programs. Compared with approaches based on bulk molecular profiles or limited spatial datasets, SQUALL provides a systematic framework for learning morphology-molecular correspondence across a large and heterogeneous spatial transcriptomics corpus, complementing existing image-based,^23,62^ image-text,^61^ and protein-oriented pretraining strategies.^79,80^

SQUALL revealed three complementary forms of transfer from spatial molecular pretraining. First, its frozen histology encoder and transcriptomics decoder enabled direct virtual spatial gene expression profiling from histology across independent cohorts and spatial profiling platforms, demonstrating that molecular information acquired during pretraining could be recovered without explicit molecular measurements. Second, joint histology-transcriptomics representations organized tissue regions into biologically meaningful spatial tumor states associated with clinically relevant tumor microenvironment clusters, immune infiltration, TLS maturation, and disease progression. Third, when only the pretrained histology encoder was applied to whole-slide images, SQUALL-derived risk scores retained prognostic value after adjustment for clinical variables, whereas risk scores derived from the other evaluated pathology foundation models did not retain consistent prognostic associations under the same multivariable adjustment. Together, these findings suggest that spatial molecular pretraining contributes not only to direct molecular decoding but also to transferable representations that capture complementary biological and clinical information from histology.

The virtual profiling results illustrate one practical consequence of this transfer. From routine H&E images alone, SQUALL enables virtual spatial gene expression profiling on independently collected cohorts and across multiple spatial profiling platforms, consistently outperforming task-specific methods and pretrained pathology models. This improvement highlights the value of incorporating large-scale paired H&E-spatial transcriptomics data into multimodal pretraining. When evaluated on matched gene-protein targets using transcript-level references, SQUALL achieved spatial concordance comparable to pretrained spatial proteomics models across shared biomarker categories, while extending virtual profiling from limited biomarker panels to transcriptome-scale prediction of more than 15,000 genes. This broader coverage enables more systematic interrogation of spatial gene expression programs directly from histology. Importantly, these predictions retained spatial information relevant to clinical interpretation. In cervical cancer, region-resolved virtual biomarkers distinguished intratumoral CD8⁺ T cell activity from immune accumulation at the tumor-stroma interface, with intratumoral CD8⁺ programs showing stronger prognostic associations. Thus, the value of virtual molecular profiling extends beyond gene-level prediction to the spatial localization of molecular programs within histological tissue architecture. Performance analyses further identified underrepresented tumor types and greater staining variability as factors associated with reduced prediction performance, helping define the conditions under which morphology-to-molecular transfer is most reliable and emphasizing the importance of diversity and quality in paired H&E-spatial transcriptomics data used for pretraining.^140^

Beyond direct molecular decoding, SQUALL joint representations resolved spatially organized tumor microenvironment states that were more clearly delineated than by comparator morphology-based, transcriptomics-based, and multimodal methods. In liver cancer, SQUALL distinguished tumor states with markedly different relationships to immune infiltration despite the presence of a mature tertiary lymphoid structure (TLS) within the same tissue. In ccRCC, lymphoid aggregates lacking features of mature TLSs coexisted with hemorrhagic, angiogenic, and suppressive myeloid programs. Building on previous observations that myeloid boundaries can constrain TLS maturation and function, our findings further associate tumor-associated hemorrhage and vascular remodeling with increased myeloid infiltration and limited TLS maturation. Together, these observations suggest that the presence of lymphoid aggregates or TLS-like structures alone may not indicate effective local antitumor immunity, and that their functional state depends on maturation and the surrounding spatial microenvironment. Performance analyses further showed that spatially aware clustering performance varied with transcriptomic background, providing insight into the data characteristics that influence multimodal representation quality. Evaluation on previously unseen datasets, together with blinded pathologist assessment, further showed that SQUALL could recover biologically interpretable tissue organization outside the pretraining corpus. In an external 4T1 tumor section, SQUALL identified necrosis-associated regions enriched for neutrophil and PMN-MDSC-like programs. These findings provide a potential spatial explanation for the established association between tumor necrosis and poor clinical outcome,^141^ suggesting that necrotic tumor regions may define a microenvironment associated with immunosuppressive myeloid states.

The same joint representation also enabled analysis of disease-associated molecular continua across large spatial cohorts. Whereas many methods integrating histology with spatial omics focus on dataset-level tasks such as segmentation,^142,143^ deconvolution,^144^ resolution enhancement,^145^ or imputation,^146^ SQUALL provides pretrained representations in which recurrent molecular states can be compared across sections, patients, and studies. In breast cancer, these representations revealed a progression-associated continuum spanning DCIS, invasive carcinomas, and more aggressive MpBC states. Progression pseudotime was accompanied by coordinated changes in cytoskeletal, calcium-signaling, estrogen-response, PI3K/AKT/mTOR, and TGF-β-associated programs, and the trajectory was also reproducible in lower-resolution spatial cohorts. The spatial coupling between TGF-β dynamics and invasive transition in human tumors also complements prior mechanistic studies in mouse models by extending these findings to direct observation in human tumors.^147^ In ovarian tumor, SQUALL similarly identified a recurrence-associated tumor-adaptive, immune-excluded region characterized by tumor predominance, reduced inflammatory cell infiltration, and NF-κB-associated programs. Together, these findings illustrate how joint histology-transcriptomics representations can connect tissue spatial organization with transcriptional programs associated with progression and recurrence.

Spatially aligned molecular pretraining also enhanced the utility of histology-only representations for clinically oriented prediction. Despite being pretrained on substantially fewer histology images than large pathology foundation models trained on images alone or image-text pairs, SQUALL achieved higher survival prediction performance across the primary independent cohorts. More importantly, SQUALL-derived risk scores remained independently prognostic after adjustment for clinical variables, indicating that the learned representation contained complementary prognostic information not fully captured by clinical covariates. In cervical cancer, high-risk attention localized to mitotically active tumor regions, whereas low-risk regions were enriched for keratinization-associated morphology, linking predicted risk to recognizable tissue phenotypes and corresponding transcriptional programs. For platinum resistance in ovarian cancer, SQUALL achieved consistently higher F1 scores across two independent cohorts, suggesting improved identification of resistant cases under class imbalance. These findings support the potential value of spatial molecular supervision for learning histology-image-only representations that remain useful even when molecular measurements are absent at inference.

Several limitations remain. First, current spatial transcriptomics technologies generally profile selected regions rather than whole-slide sections, limiting the spatial context available for multimodal pretraining. To enable scalable training across platforms and resolutions, SQUALL uses tile-level modeling and gene expression binarization, improving computational efficiency and cross-platform robustness but not explicitly capturing long-range tissue interactions or quantitative expression variation. Future large-area spatial profiling and scalable architectures may enable broader spatial and continuous molecular modeling. Second, although *histMol* spans diverse tissues and platforms, coverage remains uneven, and reduced performance in underrepresented cancers and poorly stained or sequenced samples highlights the value of broader, higher-quality pretraining data. Third, several biological analyses used sections seen during masked pretraining, although downstream labels were not used. With SQUALL kept frozen, these analyses demonstrate the utility of its pretrained representations for resolving spatial tissue states and disease-associated biological patterns without task-specific supervision or fine-tuning, rather than providing a test of out-of-distribution generalization. Transferability was evaluated separately using unseen cohorts for virtual profiling, blinded pathologist assessment for spatially aware clustering, and independent clinical datasets for outcome prediction. Future work should examine task-specific fine-tuning and broader validation before clinical deployment.

Biological systems are inherently spatial, and the continuing expansion of spatial transcriptomics provides increasing opportunities to learn representations that jointly capture morphology, molecular state, and tissue organization. Higher-throughput dense-array platforms and improved histology co-registration should further increase the scale and diversity of multimodal pretraining corpora.^26,30,148^ Future extensions could incorporate three-dimensional spatial context,^149,150^ multiplexed imaging,^81^ spatial proteomics,^80^ and pathology reports, while improving robustness across staining conditions, tissue types, and clinical populations. Rather than replacing molecular profiling, models such as SQUALL provide a route for transferring information learned from spatially resolved molecular measurements to the much larger collections of routinely acquired histological images,^70,151,152^ advancing the emerging fields of virtual tissue modeling and virtual biomarker development with potential applications in patient stratification and management.^20,56,80,153^ Together, our results support spatially aligned multimodal pretraining as a scalable strategy for connecting tissue morphology with molecular programs and extending spatial molecular insight to histopathology at scale.

## Resources availability

All processed *histMol*-low data used for stage 1 pretraining, together with a subset of processed *histMol*-high data and a version data manifest, are available at https://zenodo.org/records/23180078. Due to privacy regulations, the full *histMol*-high dataset is restricted; the corresponding raw (unprocessed) data of *histMol*-high sourced from SPATCH can be accessed via https://spatch.pku-genomics.org/#/homepage. Because the datasets comprising *histMol* are governed by heterogeneous licenses and data-use terms, we cannot redistribute all original quantitative data through a single repository. We provide a versioned *histMol* manifest documenting dataset provenance and public accession information where available for further reuse of the data.

SQUALL has been released as a Python package via GitHub. Source code and Jupyter notebooks for reproducing figures are available at https://github.com/OswaldZhang/SQUALL-release, and tutorials for SQUALL usage are accessible at https://squall.readthedocs.io/en/latest/#/. The pretrained model weights can be accessed through https://huggingface.co/zongxu/SQUALL.

Any additional information required to reanalyze the data reported in this paper is available from the lead contact upon reasonable request.

## Acknowledgements

This work was supported by the National Natural Science Foundation of China (92574301, 92374116, T2321001), Innovative Drug Research and Development-National Science and Technology Major Project (2025ZD1800400), Fundamental and Interdisciplinary Disciplines Breakthrough Plan of the Ministry of Education of China (JYB2025XDXM502), Beijing Advanced Center of Cellular Homeostasis and Aging-Related Diseases, and Peking-Tsinghua Center for Life Sciences. We thank China Bester Group Telecom Co., Ltd. for providing part of the computational resources for SQUALL’s pretraining. We also thank Dr. Stephen Williams and Dr. Michael Schnall-Levin from 10x Genomics, Inc., for assistance with formatting Visium HD data, and Mr. Zhuo Xu for help with figure preparation. We also thank the authors of the original studies for sharing their data, which enabled the curation of the *histMol* corpus. Part of the analysis was conducted on the High-Performance Computing Platform of the Center for Life Sciences at Peking University.

## Author contributions

Z.Z. (Zongxu Zhang), Z.Z. (Zexian Zeng), Y.M. and Z.L. conceived the study and designed experiments and analyses. Z.Z. (Zongxu Zhang) derived, developed, and implemented SQUALL with assistance from B.Q., Z.Q., H.X., Y.W., J.D., and P.Z.; Z.Z. (Zongxu Zhang), B.Q., Z.Q., H.X., Y.W., and J.D. performed computational analyses and visualization. Z.Z. (Zongxu Zhang), B.Q., Y.Z. (Yahui Zhao), Y.W., J.D., A.C., N.W., L.N., T.X., S.L. assembled the *histMol* corpus and curated sample metadata. Z.Z. (Zhe Zhang), Y.M. provided CGODC ovarian cancer specimens, with Z.Z. (Zongxu Zhang) and W.Z. performed histological imaging experiments of CGODC specimens with technical support from D.P.; Z.Z. (Zongxu Zhang), Y.Z. (Yahui Zhao), Z.L. designed and supervised the pathological annotation process with the help of L.X., X.X., Z.Y., J.X., C.W.; H.Z., Y.Z. (Yanping Zhao), P.R., and J.M., provided biomarker-gene categories for spatial biomarker expression predictions. Z.Z (Zexian Zeng), Y.M., and Z.L. supervised the work. Z.Z. (Zongxu Zhang), Z.Z. (Zexian Zeng), Y.M., and Z.L. wrote and revised the manuscript with the help of other authors.

## Declaration of interests

D.P. received sponsored research funding from Bayer AG and Boehringer Ingelheim. These grants were not related to the research reported in this study.

## Methods

### PATIENTS AND SAMPLES

No human tissue specimens were newly profiled by spatial transcriptomic sequencing specifically for the development or evaluation of SQUALL. The human specimens and associated clinical information represented in the in-house datasets incorporated into *histMol* were obtained through independently conducted source studies under protocols approved by the Research and Biomedical Ethical Committee of Peking University (IRB00001052-24061) and the ethics committee of Chinese PLA General Hospital #S2022-403-01 with written informed consent obtained when appropriate. These previously generated datasets were subsequently incorporated into the SQUALL pretraining corpus.

Given the importance of data provenance for foundation model evaluation, we provide a versioned *histMol* manifest documenting dataset provenance, available public accession identifiers, and non-identifying metadata at the cohort, patient, and section levels. The manifest enables investigators to assess potential pretraining exposure when designing or interpreting downstream benchmark studies.

Histology images from the Chinese Gynecologic Oncology Multi-Omics Data Center (CGODC) were obtained from specimens collected at the Chinese PLA General Hospital under approval from the Ethics Committee of the Chinese PLA General Hospital (#S2022-403-01), with written informed consent obtained when appropriate. These specimens were used only for histology imaging analyses and were not used in the construction, training, or development of *histMol*.

### COLLECTION AND ASSEMBLY THE HISTMOL CORPUS

We assembled *histMol*, a multimodal pretraining corpus of paired histology images and spatial transcriptomics profiles spanning 33 tissues, 12 platforms, and 3,446 tissue sections. The corpus contained approximately 1.93 billion sequencing spots or bins collected from 15 public repositories representing 308 studies, together with in-house datasets generated by us (**Figures 1A and 1B**; **Tables S1 and S2**).

Based on imaging and sequencing resolution, *histMol* was divided into low-resolution (*histMol*-low) and high-resolution (*histMol*-high) components. *histMol* -low comprised 6,427,526 spots from 3,243 sections representing 1,952 samples and was used for stage 1 pretraining. *histMol*-high comprised 1,919,973,520 bins from 203 sections representing 181 samples and was used for stage 2 pretraining. For *histMol*-high, histology images were standardized to 0.5 μm per pixel and aligned with 2-μm sequencing bins. After preprocessing, *histMol* contained 74,773 low-resolution and 615,767 high-resolution tiles (**Figures 1B**, **1D, S1E, S1F, and S1H**; **Tables S1, S2, S4, and S8**).

### Dataset curation

Publicly available spatial transcriptomics datasets were collected from the Gene Expression Omnibus, ArrayExpress, Zenodo, Mendeley Data, SOAR, CELLxGENE, the National Genomics Data Center, Figshare, the Single Cell Portal, Dryad, Spatial Research, the Human Cell Atlas, KU Leuven Research Data Repository, STOmics Data, and 10x Genomics released prior to November 1, 2024. For each dataset, we collected the raw or processed spot-by-gene or bin-by-gene count matrix, spatial coordinates, paired histology images, and available sample metadata.

Gene expression matrices were obtained in formats including HDF5, H5AD, RDS, MTX, GEM, and delimited text files. Histology images were collected in PNG, JPEG, TIFF, or whole-slide image formats. Dataset inventories from existing spatial transcriptomics resources were cross-referenced with HEST-1k and STimage-1K4M during curation to improve coverage and identify potentially duplicated records. Duplicate sections were manually identified and removed. Sections without matched histology and spatial transcriptomics data, sections containing non-H&E images, and sections with insufficient tissue area or inadequate image quality were excluded.

For Visium datasets lacking complete spatial coordinate information, spot barcodes were matched to the corresponding barcode whitelist, and the spatial coordinates were reconstructed using Loupe Browser where applicable. For datasets in which the sequencing array and histology image were not spatially registered, anchor-based image alignment was performed using SimpleITK. Tissue foreground masks were generated using Otsu thresholding, and sequencing spots or bins outside the tissue area were excluded.

All curated low-resolution datasets were converted into a unified AnnData representation and stored as H5AD files. Data conversion and preprocessing were implemented using Scanpy, Seurat, SciPy, NumPy, pandas, and scikit-image. Platform-specific metadata, including species, tissue type, disease status, preservation method, imaging resolution, sequencing resolution, and array geometry, were retained whenever available.

High-resolution paired histology and spatial transcriptomics data were obtained from STAGE, SPATCH, publicly released 10x Genomics datasets, and other in-house cohorts. Raw bin-by-gene matrices were stored in HDF5 files, spatial coordinates were stored in Parquet or platform-specific feature files, and the paired histology images were available at approximately 40× magnification. When available, a section-specific 3 × 3 perspective-transformation matrix mapping sequencing-bin coordinates to histology-image coordinates were retained for subsequent texture mapping.

### Gene harmonization and expression preprocessing

To construct a unified cross-species gene space, genes with non-zero expression were first collected separately from human and mouse sections. One-to-one human–mouse orthologs were identified using the Ensembl API, and the resulting gene set was intersected with the genes included in the Visium HD probe set. This procedure produced a shared gene vocabulary of 15,757 genes (**Figures S1A and S1B; Table S3**).

For each tissue section, genes were reordered according to this shared vocabulary. Genes not measured by a given platform were represented by zero padding, ensuring that all sections had an identical gene dimension. This harmonization enabled joint pretraining across species, platforms, and spatial resolutions.

For quality assessment, library size and the number of detected genes were calculated for each spot or bin using *scanpy.pp.calculate_qc_metrics*. Gene-level expression distributions were summarized using t-digest. Spatial transcriptomics profiles were substantially sparser than large-scale single-cell RNA-sequencing profiles, and library size was strongly correlated with the number of detected genes across the major spatial profiling platforms (**Figures S1C and S1D**). Based on these observations, gene-expression values were binarized before model input. Previous studies have shown that binarized transcriptomic representations retain biologically meaningful information and can serve as effective reconstruction targets for representation learning. For each spot or bin, genes with non-zero counts were assigned a value of 1, whereas genes with zero counts were assigned a value of 0. This representation preserved gene-detection patterns while reducing sensitivity to platform-specific count scales, sequencing depth, and expression-magnitude differences.

Histology-image resolution was estimated from the available scanner metadata, fiducial frames, spatial-array geometry, or inter-spot distances. Resolution metadata were retained and explicitly supplied to SQUALL during pretraining.

### Background white-balance correction

Histology images acquired under suboptimal imaging or illumination conditions were corrected using an approach based on the perfect-white-reflection assumption. Tissue foreground was first identified using the tissue mask, and fiducial frames or other non-tissue structures were excluded where applicable. The remaining background region was treated as the white reference.

The median red, green, and blue intensities of the background pixels were calculated to reduce sensitivity to artifacts and uneven illumination. Channel-specific scaling factors were then estimated relative to pure white, defined as RGB = [255, 255, 255], and applied linearly to the entire image. Pixel values were clipped to the valid image-intensity range after transformation. This procedure reduced background color variation while preserving the relative hematoxylin and eosin staining patterns within tissue regions (**Figure S2A**).

### Conversion of low-resolution data into a grid-like representation by pseudo-labeling

Spatial transcriptomics platforms represented in *histMol*-low used heterogeneous array geometries, including square, hexagonal, and centered rectangular lattices, with platform-dependent spot diameters and center-to-center distances. To provide a unified image-like representation for multimodal pretraining, the original spot-level profiles were converted into a regular grid through a procedure termed pseudo-labeling (**Figure S2B**).

For each section, the image-space boundary of every sequencing spot or bin was determined from its spatial coordinates and platform geometry. The spatial coordinates were then mapped to a resolution-aware raster grid. At a histology resolution of 0.5 μm per pixel, a 2-μm sequencing unit corresponded to a 4 × 4-pixel image region. The binary gene-expression vector associated with each original spot was assigned to the grid cells covered by that spot.

Because many low-resolution platforms contain physical gaps between adjacent spots, unobserved grid cells within the tissue area were filled using nearest-neighbor interpolation. This procedure was selected for its computational efficiency and its limited effect on library-size distributions relative to the alternative interpolation methods evaluated (**Figures S2C-S2E**).

### Alignment of high-resolution data by texture mapping

In *histMol*-high, sequencing data were represented as contiguous 2-μm square bins arranged on a regular lattice. The corresponding histology images were acquired at approximately 40× magnification and were mapped to the sequencing coordinate system using a section-specific 3 × 3 perspective-transformation matrix.

To establish pixel-level correspondence between the histology image and sequencing grid, we developed a texture-mapping operator inspired by real-time graphics rendering (**Figures S2F-S2J**). A target raster grid was first generated at 0.5 μm per pixel. The corner coordinates of each target raster element were transformed from sequencing space into histology-image space using the perspective-transformation matrix.

Each transformed target element formed a quadrilateral in the original histology image. The intersection between this quadrilateral and the original image pixels was determined using a ray-casting procedure. The areas of the resulting overlap polygons were calculated using the shoelace formula and were used as weights for area-based downsampling. This procedure produced a resolution-standardized histology image that was spatially aligned with the 2-μm sequencing-bin grid.

For transformed coordinates extending beyond the original image boundary, the corresponding output pixels were imputed using the median RGB intensity of the image background. The texture-mapping procedure was implemented in Python and accelerated using Numba just-in-time compilation and parallel-range multithreading. In benchmark testing, mapping an approximately 180.6-million-pixel 40× histology image to the 20× target resolution required approximately 10 min on a 32-core CPU server. The resulting correspondence between sequencing bins and histology pixels was stored in Parquet format to support efficient input/output operations.

### Pathological taxonomy harmonization

Tissue and disease annotations were curated from the original publications, pathology reports, and accompanying metadata. Three experienced pathologists reviewed the collected annotations and harmonized the tissue hierarchy by consensus.

For metastatic specimens, tissue labels were assigned according to the anatomical site of the profiled tissue rather than the presumed primary tumor site. Hippocampal, olfactory bulb, and cortical specimens were grouped under brain. Embryonic specimens from different developmental stages were grouped under embryo, whereas fetal tissues were assigned to their corresponding anatomical categories. Salivary gland, periodontal, nasopharyngeal, and oral tissues were grouped under head and neck. Colon, small intestine, rectosigmoid, duodenum, and ileum were grouped under intestine.

When a physical section contained spatially distinct tissue types or multiple tissue-microarray cores, it was divided into single-label analytical sections for section-level classification. These derived units were used only for classification analyses and did not alter the physical-section count of 3,243 for *histMol*-low or 3,446 for the complete *histMol* corpus (**Tables S1 and S6**).

Cancer labels were harmonized only for human tumor sections. Each section was assigned a single cancer label based on the National Cancer Institute classification by anatomical location or organ system, with minor modifications to merge redundant terms (https://www.cancer.gov/types/by-body-location). For *histMol*-high, pathological subtype labels were obtained from the original pathology reports and were manually reviewed during metadata curation (**Tables S7 and S8**).

### Pathological annotations

For the *histMol*-low breast cancer cohort, three to five board-certified pathologists reviewed the available H&E images and established a consensus annotation hierarchy. Regions were annotated at the finest reliable resolution supported by the corresponding histology image. The resulting annotations covered major pathological compartments, breast cancer subtypes, stromal and interstitial regions, and other relevant tissue structures (**Figures S7B and S7C**; **Table S51**).

For the high-resolution gynecological tumor subset used in stage 2 evaluation and downstream spatial analyses, pathological regions were annotated in QuPath. Annotation classes and granularity were determined by consensus among the participating pathologists. Annotation objects were exported from the corresponding QPData files and converted into GeoJSON polygon representations. Each image tile was assigned the label of its corresponding polygon. Tiles overlapping multiple incompatible polygon classes were excluded from supervised evaluation (**Figures S8A and S8B**; **Table S22**).

### Data tiling and storage

Tissue foreground masks were generated from the histology images using Otsu thresholding. For high-resolution images, segmentation was performed on an image downsampled by a factor of 20 to reduce computational cost. Morphological erosion and dilation were applied to remove small artifacts and fill minor gaps. The resulting tissue mask was then resized to the original image dimensions.

Spatially aligned histology images and gene-expression grids were exhaustively partitioned into non-overlapping tiles. Each histology tile had a shape of 224 × 224 × 3 pixels, and the corresponding molecular tile was represented as a 56 × 56 × 15,757 tensor. Tiles containing more than 50% background area were excluded.

Histology tiles were stored as TIFF files to preserve pixel values, whereas molecular tiles were stored as sparse PyTorch tensors in PT format. The position of each tile relative to the complete tissue section was encoded in the file metadata or filename. Histology-image intensities were scaled to the range [0, 1]. ImageNet mean and standard-deviation normalization and random resized cropping were not applied during pretraining.

A customized collation function was implemented for efficient batching of paired image and molecular tensors. Data were loaded using a streaming iteration strategy to reduce memory and storage-access overhead during distributed model training. After all preprocessing steps, the final corpus comprised 74,773 *histMol*-low tiles and 615,767 *histMol*-high tiles.

### DEVELOPMENT OF SQUALL

#### Overview of SQUALL

SQUALL is a multimodal self-supervised learning framework designed to integrate histological morphology with spatial gene expression programs. The model was pretrained on the *histMol* corpus using a two-stage continual-pretraining strategy that accommodated the different imaging and sequencing resolutions represented in the corpus (**Figures 1C and 1D**).

After pretraining, SQUALL can be used in several configurations. The complete encoder supported joint histology–transcriptomics representation learning, whereas the histology branch could be applied independently to spatial histology tiles or whole-slide images. The transcriptomics decoder could additionally be coupled with the frozen histology encoder for virtual spatial gene-expression profiling (**Figure 1E; Table S9**). The model architecture, pretrained parameters, and implementation were released to support reproducibility and further method development.

#### Tissue section representation

Spatial transcriptomics protocols generally acquire molecular profiles and H&E-stained histology images from the same physical tissue section or from directly corresponding tissue material. H&E staining provides pixel-level information about nuclear morphology, cytoplasmic composition, extracellular matrix organization, and tissue architecture. Histology images are commonly acquired at approximately 20× magnification, corresponding to approximately 0.5 μm per pixel, or at approximately 40× magnification, corresponding to approximately 0.25 μm per pixel.

The spatial resolution of molecular measurements depends on the profiling platform and is generally lower than that of the paired histology image. To establish a common spatial reference system, both modalities were represented on aligned two-dimensional grids.

For a tissue tile, let ℐ ∈ ℝ*^r^*^ℎ×*rw*×*C*^ denote the histology image and X ∈ ℝ^ℎ×*w*×*g*^ denote the corresponding molecular-expression grid. Here, *C* = 3 denotes the RGB image channels, *g* denotes the number of genes in the harmonized gene vocabulary, ℎ and *w* denote the height and width of the molecular grid, and *r* denotes the ratio of the molecular-grid spacing to the histology pixel size.

Each molecular patch *X_ij_* ∈ ℝ^ℎ^ ^×*w*^ ^×*g*^ corresponded spatially to an *r* × *r* histology patch,

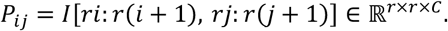

For example, when 2-μm sequencing bins were paired with histology images represented at 0.5 μm per pixel, *r* = 4, such that each sequencing unit corresponded to a 4 × 4*-*pixel histology patch. Both modalities were partitioned into spatially aligned, non-overlapping tiles before model input. This representation preserved local correspondence while limiting the computational cost of attention-based modeling, similar to previous pathology image foundation models’ design.

### Complementary multimodal masking

SQUALL was trained using complementary masking across the histology and spatial transcriptomics modalities (**Figure 1C**; **Figures S3A and S3B**). At each spatial location, either the histology patch or its corresponding gene-expression vector was presented to the encoder, while the paired component from the other modality was masked and used as a reconstruction target.

For a molecular grid of size ℎ × *w*, we sampled a binary masking matrix *M* = [*m_i_*_,*j*_]_ℎ×*w*_ ∈ 0,1^ℎ×*w*^, where

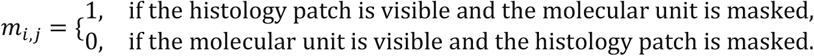

The molecular-grid mask was expanded to the histology-pixel domain as

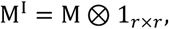

where ⊗ denotes the Kronecker product. Consequently, all pixels within the *r* × *r* histology region corresponding to a molecular unit shared the same masking status.

The visible histology and molecular sets were defined as

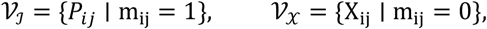

and the corresponding reconstruction targets were

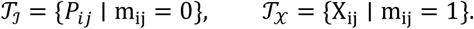

Thus, the number of visible histology tokens and visible molecular tokens was

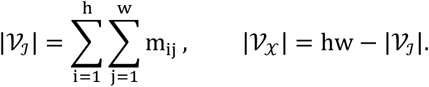

This complementary masking scheme maintained one visible modality at each spatial location while preventing the encoder from directly observing both paired inputs simultaneously. The model was therefore required to use spatial context and cross-modal relationships to reconstruct the missing histological or molecular information.

The approach can be viewed as a modality-aware denoising objective. Reconstruction of a masked molecular vector required the model to infer gene expression programs from the corresponding tissue morphology and surrounding molecular context. Conversely, reconstruction of a masked histology patch required information from spatial molecular states and neighboring histological structures. This reciprocal constraint encouraged the model to learn spatially coherent representations that retained both modality-specific and shared biological information.

### Input projections and positional encoding

Visible histology patches and molecular vectors were then projected into an embedding space of dimension *p*. For a visible molecular vector *X_ij_*, the input token was obtained using a learnable linear projection:

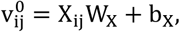

where *W_X_* ∈ ℝ*^g^*^×*p*^ and *b_X_* ∈ ℝ*^p^*.

For a visible histology patch *P_ij_*, a lightweight convolutional projection mapped the *r* × *r* × *C* input region to a *p*-dimensional image token:

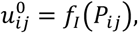

where *f* : ℝ*^r^*^×^ *^r^*^×^ *^C^* → ℝ*^p^*.

The visible image and molecular tokens were combined into a single spatially indexed sequence. Each token retained its modality identity and two-dimensional tissue coordinate.

To accommodate differences in spatial resolution across technologies, physical spatial information was explicitly incorporated into the positional encoding. For a token located at grid coordinate *c_ij_* = [*i*, *j*], its absolute positional embedding was defined as

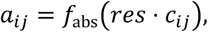

where *res* denotes the physical resolution associated with the tissue section and *f*_abs_ is a shallow learnable projection network.

The initial token representation was

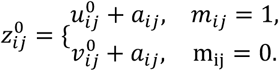

In addition to absolute position, SQUALL used a learnable relative positional bias. For tokens at coordinates *c_ij_* and *c_st_*, the physical relative displacement was

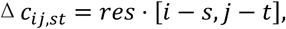

and the corresponding attention bias was

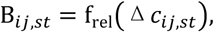

where *f*_rel_ is a shallow neural network. This formulation allowed the model to distinguish identical grid offsets arising at different physical resolutions (**Figure S3D; Table S11**).

### SQUALL encoder

The SQUALL encoder was based on a Transformer architecture using a mixture-of-modality-experts design (MoME). Only visible inputs were passed through the encoder, reducing computational cost in a manner analogous to masked autoencoders (**Figure 1C; Figures S3A and S3C; Table S17**).

For the token sequence *z_l_* at layer *l*, the query, key, and value matrices were calculated as

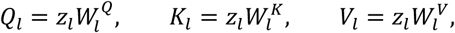

where 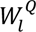, 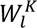, and 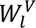 are learnable projection matrices.

For each attention head, attention was calculated as

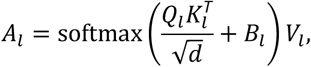

where *d* denotes the head dimension and *B_l_* denotes the resolution-aware relative positional bias.

Since image and molecular tokens were included in the same attention sequence, the attention matrix contained both intra-modality interactions and cross-modality interactions.

The output of the attention block was updated using pre-layer normalization and a residual connection:

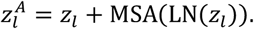

In the modality-specific layers, histology and molecular tokens were routed through separate feed-forward experts:

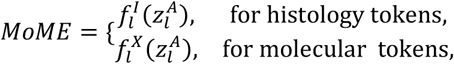

with

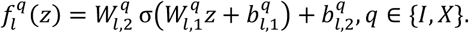

Here, *σ* denotes the GELU activation function. The layer output was then calculated as

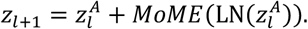

The shallow encoder layers retained modality-specific feed-forward experts to extract specialized histological and molecular features. In the deeper fusion layers, modality-specific routing was replaced by shared feed-forward layers, allowing the two token types to be progressively integrated into a common representation space.

This architecture separated modality-specific feature transformation from cross-modal attention. Consequently, SQUALL could process paired histology and transcriptomics during multimodal inference while retaining the ability to process histology alone when molecular measurements were unavailable.

#### Pretraining objective

SQUALL was pretrained using masked multimodal data reconstruction. For each paired histology image and gene-sequencing profile (ℐ, X), random masks partition each modality into visible components (ℐ*_v_*, X*_v_*) and masked targets (ℐ*_m_*, X*_m_*). The SQUALL encoder, pretrained at scale on the *histMol* corpus, maps the visible inputs to a shared latent representation, while shallow modality-specific transformer decoders reconstruct the masked image pixels and gene-sequencing units (**Figures S3A and S3B**).

Let *M^I^* and *M^X^* denote the complementary masks applied to the image and sequencing modalities, respectively. Under this notation, SQUALL obtains the latent representation as

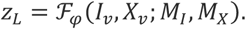

Equivalently, this deterministic mapping can be represented by the Dirac conditional distribution

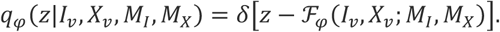

Since SQUALL does not fit a probabilistic latent prior during pretraining, learning is formulated directly in terms of the masked conditional distribution. Conditional on the latent representation, we assume that the masked components of the two modalities are independent:

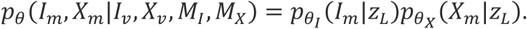

Let *n*_ℐ_ = |*M*_ℐ_| and *n*_X_ = |*M*_X_|. Using the per-modality mean reduction employed during training, the masked negative log-likelihood is

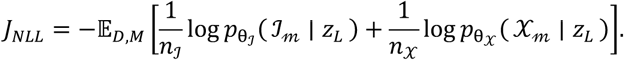

Assuming independent Gaussian reconstruction errors with a fixed variance, this objective reduces, up to constants, to

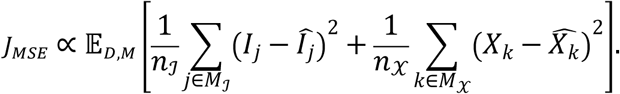

For robustness to large residuals, we replace the squared error with the smooth *L*_1_ loss

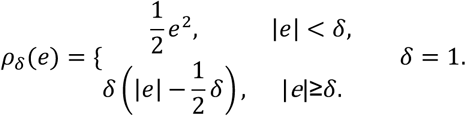

#### Stage-wise continual pretraining

SQUALL was pretrained using a two-stage continual-pretraining strategy to accommodate the complementary spatial scales of *histMol*-low and *histMol*-high (**Figure 1D; Figure S3I; Table S24**).

During stage 1, the model was pretrained on *histMol*-low. This stage exposed the model to a broad range of tissues, species, preservation protocols, spatial transcriptomics platforms, and imaging resolutions. The resulting parameters provided both a transferable feature extractor and an initialization for subsequent high-resolution training.

During stage 2, the SQUALL model was initialized using the stage 1 parameters and continually pretrained on *histMol*-high. This stage used spatially dense sequencing bins and higher-resolution histology images to refine the correspondence between local morphology and molecular state. The model architecture was retained across stages, while the physical-resolution encoding allowed inputs from the two corpus components to be interpreted in their corresponding spatial units.

Training convergence was monitored using the multimodal reconstruction loss and ancillary classification probes. Stage-specific training was continued until the reconstruction loss and probe classification performance reached stable plateaus. Masked image and molecular reconstruction outputs were additionally inspected throughout training (**Figures S3K and S3L**).

The optimized learning rate, batch size, training duration, warm-up schedule, masking ratio, optimizer, and weight-decay configuration used for each stage are summarized in (**Tables S27 and S28**).

### DESIGN EVALUATION OF SQUALL

#### Ancillary probing tasks

We developed section-level and tile-level classification probes to compare candidate model configurations during pretraining. These probes were designed as lightweight evaluations of representation quality and were not used as pretraining objectives.

For section-level evaluation, tile representations were extracted from each *histMol*-low section using the evaluated encoder. A [CLS] token was included in each tile sequence, and its final encoder representation was used as the tile-level feature *x_i_* ∈ *R^p^*.

Tile representations from the same tissue section were aggregated using gated attention-based multiple-instance learning. The section representation was

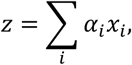

where *α_i_* is the learnable weighted to tile *i* that prioritizes tile level embedding and is calculated as:

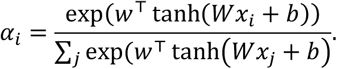

A shallow linear classification head was trained using the aggregated section representation. The prediction target was either the harmonized tissue label or cancer-type label, depending on the evaluation setting. Data were divided at the section level to prevent tiles from the same section from appearing in both training and evaluation partitions.

For tile-level evaluation, the evaluated encoder was frozen, and a linear classifier was trained on the [CLS] representation of each tile. This evaluation was performed on the pathologist-annotated *histMol*-high subset. Each tile was assigned a pathological label based on the polygon annotation covering that tile; tiles overlapping incompatible annotations were excluded.

Unless otherwise specified, classification performance was evaluated using top-1, top-3, and top-5 accuracy, weighted F1 score, and area under the receiver operating characteristic curve. Identical probing architectures, data partitions, and optimization settings were used when comparing model configurations.

#### Architecture and pretraining design evaluations

We performed controlled design evaluations to determine the final SQUALL architecture and pretraining configuration. Each comparison altered a defined model, preprocessing, or data-composition factor while retaining the remaining applicable settings.

The evaluated factors included: (1) model capacity and pretraining duration; (2) positional-encoding strategy; (3) explicit encoding of physical resolution; (4) inclusion of mouse samples; (5) inclusion of multiple spatial transcriptomics platforms; (6) histology stain normalization; (7) pretraining-data scale; and (8) stage-wise initialization for high-resolution training (**Figures S3F, S3G, and S3I; Tables S11-S25**).

Both SQUALL-base and SQUALL-large were evaluated to characterize model-scaling behavior. Models were trained using either a 30% subset or the complete *histMol*-low corpus to assess the contribution of pretraining-data scale. Representation quality was measured at multiple training checkpoints to evaluate the effect of pretraining duration.

For positional encoding, the evaluated configurations included sinusoidal absolute positional embeddings, learned absolute positional embeddings, two-dimensional rotary positional embeddings, learnable relative positional biases, bucketed relative positions, and relative positions encoded using a shallow neural network.

We additionally evaluated combinations of absolute and relative positional encodings, with or without explicit physical-resolution information.

To assess the contribution of cross-species data, a human-only training set was constructed by removing mouse samples from the corresponding *histMol*-low subset. The number of optimization iterations and downstream evaluation protocol were matched to those used for the mixed-species reference setting.

To assess the contribution of platform diversity, a Visium-only training set was compared with the corresponding multi-platform training set. The number of sampled tiles and optimization steps was controlled between settings.

For stain-normalization evaluation, a median stain basis was estimated from *histMol*-low using the Macenko method. The normalized images were used to train the corresponding model setting, while all non-image inputs and optimization settings were retained.

The contribution of stage-wise continual pretraining was evaluated using three high-resolution training conditions: initialization from stage 1 parameters, training the same architecture from random initialization, and direct application of the stage 1 model without additional high-resolution pretraining. All models were evaluated using the same annotated high-resolution tile-classification task.

For learning-rate and batch-size selection, we evaluated learning rates in

- learning rate ∈ {5e−4, 2e−4, 1e−4, 5e−5, 2e−5, 1e−5, 5e−6},
- batch size ∈ {64, 128, 256, 512, 1,024}.

At each training stage, the learning rate was first selected using a fixed batch size, followed by batch-size selection while keeping the selected learning rate fixed. The final configuration was selected using performance on the predefined probing partitions. The complete model and optimization settings are provided in **Tables S16, S27 and S28**.

#### Cross-task evaluation of preprocessing, architecture, and data composition

We performed an additional controlled comparison of nine model settings spanning expression preprocessing, spatial interpolation, model architecture, initialization, and pretraining-data composition. All settings used the same 30% pretraining data tiles to control for training set size and computational cost. Each setting modified one principal component relative to the reference configuration while retaining the remaining applicable settings.

The reference setting used binarized gene expression with nearest-neighbor interpolation for pseudo-labeling. Two interpolation variants for pseudo-labeling were compared with this reference: (1) linear interpolation; (2) Clough–Tocher interpolation. We also included two data normalization methods: (1) rank-transformed expression; (2) log1p-normalized expression. These comparisons separately assessed the effects of gene-expression representation and spatial interpolation.

Three architecture or initialization variants were evaluated. In the UNI-init setting, image features were first extracted using UNI and then projected into the SQUALL histology encoder, while the remaining multimodal architecture was retained unchanged. In the direct-expression-prediction setting, the molecular masking ratio was set to 100%, requiring the model to predict the complete expression profile using histology context without observing molecular tokens. In the image-only masked-modeling setting, the multimodal reconstruction objective was replaced by an image-only masked-autoencoder objective to evaluate the benefit of molecular alignments. A data composition variant was constructed by excluding mouse samples from the pretraining corpus while retaining the reference preprocessing, interpolation, and model settings.

All nine configurations were evaluated using identical downstream protocols across four tasks: virtual spatial gene expression profiling, representative spatially aware clustering, survival prediction, and platinum-resistance prediction. Performance was assessed using the corresponding task-specific metrics reported in (**Figure S3J; Table S26**). To summarize performance consistency across tasks, we counted, for each configuration, the number of metrics on which it ranked among the top three. We additionally recorded the wall-clock time required for preprocessing and the disk space occupied by each resulting dataset.

#### Hardware and software

All model development and pretraining experiments were conducted using Python 3.8.13 and PyTorch 2.4.0 with CUDA 12.1. Large-scale pretraining was performed using 32 NVIDIA H100 GPUs with 80 GB of memory per GPU. GPUs were distributed across multiple compute nodes, with NVLink used for intra-node communication and InfiniBand used for inter-node communication. Distributed training was implemented using PyTorch DistributedDataParallel.

Histology images were loaded using scikit-image. Custom PyTorch datasets, collation functions, and streaming data loaders were implemented to improve input/output efficiency when loading paired histology and sparse molecular tensors.

The total training time was approximately 177.3 h for stage 1 and 26.8 h for stage 2. Training losses, probing metrics, learning-rate schedules, and hardware utilization were monitored using TensorBoard. Detailed stage-specific hardware utilization and training configurations are provided in (**Figure S3K, Tables S27 and S28**).

Unless otherwise specified, downstream evaluation experiments were performed using a single NVIDIA A100 GPU with 80 GB of memory.

### VIRTUAL SPATIAL GENE EXPRESSION PROFILING

#### SQUALL inference

Each H&E image was processed at 20× magnification when available; otherwise, the highest-resolution image available was used. Images were then partitioned into non-overlapping 224 × 224-pixel tiles. For each tile, the frozen SQUALL histology encoder generated spatial histology representations, which were passed to the frozen transcriptomics decoder while all transcriptomic inputs were masked. For each input tile, SQUALL generated a spatial prediction tensor of 56×56×15,757. Predictions from individual tiles were placed at their corresponding image coordinates to reconstruct section-level virtual expression maps. SQUALL parameters were kept frozen throughout the evaluations.

#### Comparison with H&E-to-ST models

We evaluated whether multimodal pretraining enabled SQUALL to generate spatially resolved gene-expression profiles directly from routine H&E histology. During virtual profiling, the pretrained histology encoder and transcriptomics decoder were frozen. An H&E image was provided together with masked transcriptomic tokens, and the corresponding spatial gene-expression profiles were generated in a single forward pass without dataset-specific fine-tuning (**Figure 2A; Table S9**).

The H&E-to-ST comparison comprised three settings:

1. Controlled comparative evaluation on Xenium 5K sections, comparing SQUALL with task-specific H&E-to-expression models, pathology foundation models, and multimodal H&E–ST embedding models trained on Visium HD sections from matched tumor categories.
2. Cross-cohort and cross-cancer generalizability using 117 newly collected Visium sections with pretrained H&E-to-ST foundation models.
3. Evaluation at higher spatial resolution using three independent Visium HD sections.

#### 1. Controlled comparative evaluation using Xenium 5K sections

Two Xenium 5K datasets were used for this evaluation. Neither dataset was included in SQUALL pretraining at the section or patient level. Two SPATCH Visium HD sections—one ovarian cancer section and one hepatocellular carcinoma section—were used to train models requiring task-specific training. Neither Xenium 5K evaluation section was used to fit a prediction model. The comparison was conducted using the 4,421 genes shared among Xenium 5K, the *histMol* gene vocabulary, and all evaluated model outputs (**Figure 2B; Tables S29–S31**).

#### Implementation of comparison models

All comparison models were evaluated using 224×224-pixel H&E tiles. Xenium 5K measurements were aggregated onto the tile level by averaging all measurements. Only spatial locations jointly covered by the prediction map, measured-expression map, and tissue mask were considered valid for quantitative evaluation. Background and non-overlapping locations were excluded. For each gene, predicted and measured expression values were quantile transformed and mapped to the interval [0,1] to ensure robust evaluation. Models requiring task-specific fitting were trained using the Visium HD sections. Released pretrained weights and official preprocessing procedures were used where available.

##### iStar

iStar used a pretrained hierarchical Vision Transformer histology backbone followed by an MLP gene-expression prediction head. Its input pipeline was adapted to support Visium HD input, while the remaining applicable settings followed the released implementation and trained with a 400-epoch schedule.

##### ST-Net

ST-Net use an ImageNet-pretrained DenseNet-121 backbone followed by a fully connected expression-prediction layer. Visium HD measurements were aggregated within each H&E tile to generate tile-level targets, and the model was trained using the published protocols.

##### Path2Space

Path2Space use CTransPath for histology feature extraction followed by an MLP expression-prediction network. The model was fitted using its nested-fold ensemble strategy on the Visium HD sections. Predictions from all ensemble members were averaged to produce the final Xenium 5K predictions. The released preprocessing procedure, including Macenko stain normalization, was retained, with outputs generated at the tile level.

##### EGN

EGN combines exemplar retrieval with an exemplar-guided Transformer prediction network. Morphologically similar reference tiles and their expression profiles were incorporated as auxiliary information during prediction. Visium HD measurements were aggregated into tile-level targets, and the output layers were adapted to the matched gene panel.

##### DeepPT

DeepPT use an ImageNet-pretrained ResNet-50 encoder, an autoencoder for feature compression, and an MLP for expression prediction. H&E tiles were processed using Macenko stain normalization. The original slide-level aggregation step was removed so that predictions were retained at the tile level.

##### Hist2ST

Hist2ST combines a ConvMixer image encoder, a Transformer, and a graph neural network. Each H&E tile was treated as a graph node, and spatial neighborhoods were constructed from tile coordinates. The published expression-modeling and self-distillation objectives were retained.

##### OmiCLIP

OmiCLIP was applied using its released pretrained image–transcriptomics representation and similarity-based expression inference procedure. Image embeddings were compared with transcriptomics-defined reference embeddings, and associated expression profiles were aggregated using similarity-derived weights. Since OmiCLIP was pretrained on the STbank dataset, we used the released model and feature similarity matrix without retraining or fine-tuning, following the original reported protocol to preserve the original feature space. For evaluation on Xenium sections, spatial measurements were aggregated at the tile level to match the spatial scale of legacy ST sequencing resolutions. In this comparsion, OmiCLIP was categorized as a multimodal embedding-based approach to evaluate SQUALL against diverse embedding strategies for H&E- to-ST predictions.

##### Multi-Embed

Multi-Embed used a frozen UNI encoder to generate histology features, which were aligned with matched transcriptomic representations in a shared latent space. The cross-modal adaptation module and downstream expression-prediction head were trained, whereas the UNI encoder remained frozen.

##### Pathology foundation models

The released UNI, PLIP, and Virchow image encoders were frozen and used to extract tile-level histology representations. Each encoder was paired with an MLP regression head. For the Xenium comparsion, the prediction heads were trained using the Visium HD sections. Tile-level targets were defined as the mean log-normalized expression of all spatial observations contained within the corresponding tile. The same data split, optimization procedure, and checkpoint-selection criterion were used for all three models.

For compasrion, ST-Net, Hist2ST, EGN, DeepPT, Path2Space, and MultiEmbed, together with MLP heads trained on frozen pathology foundation models derived image features, were trained using the SPATCH Visium HD dataset. A held-out subset of the Visium HD data was used for model selection, and hyperparameters and checkpoints were determined based on validation performance before evaluation on the Xenium sections. iStar was implemented according to its original training pipeline: the model was trained for 400 epochs with no separate validation set or checkpoint-selection procedure was used, and the final checkpoint was used for Xenium inference. OmiCLIP was applied directly using its released image–ST similarity matrix. Xenium expression measurements were used exclusively for final evaluation and were not used for training, validation, hyperparameter tuning, or checkpoint selection for any method. Additional architectures, optimization parameters, and configurations are provided in **Tables S30 and S31**.

#### 2. External conventional Visium cohorts

We assembled 117 conventional-resolution Visium sections from eight newly collected cohorts. All sections were collected after completion of SQUALL pretraining and after release of the comparator pretrained weights. They were not used for SQUALL pretraining or any dataset-specific fine-tuning, adaptation, or calibration of the evaluated models. This comparison was restricted to models supporting direct inference without target-cohort retraining. SQUALL was therefore compared with OmiCLIP and STPath using the 14,766 genes shared between all three models and the measured Visium expression profiles (**Figures 2F and 2G; Tables S35–S37**).

STPath is a generative foundation model pretrained on large-scale paired whole-slide images and spatial transcriptomic profiles. It integrates histology features extracted from pretrained Prov-GigaPath, spatial geometry, organ identity, sequencing-technology information, and gene representations within a geometry-aware Transformer, and was pretrained to recover masked gene expression directly. All evaluations were performed under a direct-inference setting.

#### 3. Independent Visium HD sections

For transcriptome-wide evaluation on Visium HD sections, SQUALL was compared with UNI, PLIP, Virchow, OmiCLIP, and STPath. SQUALL, OmiCLIP, and STPath were applied directly using their released pretrained parameters without section-specific fitting. Since UNI, PLIP, and Virchow are image encoders rather than direct gene-expression predictors, each frozen encoder was paired with a newly initialized regression head evaluated using a five-fold out-of-fold procedure within each section. Tiles from each section were divided into five non-overlapping folds. In each iteration, three folds were used to fit the regression head, one-fold was used for validation, and the remaining fold was held out for testing. The validation fold was rotated relative to the test fold so that every tile was included exactly once in an out-of-fold test set. Training was performed for a maximum of 50 epochs with validation-based early stopping, and the checkpoint with the lowest validation loss was retained. Predictions from the five mutually exclusive test folds were restored to their original spatial coordinates to construct the complete section-level prediction map. Thus, every reported UNI, PLIP, and Virchow prediction was generated when the corresponding tile had been excluded from both training and validation. These models were evaluated as frozen pathology encoders with locally fitted out-of-fold regression heads, whereas SQUALL, OmiCLIP, and STPath were evaluated by direct frozen inference (**Figures 2F**, **2H, and 2I; Tables S31, S38, and S39**).

#### Evaluation with pretrained spatial proteomics models

SQUALL was separately compared with ROSIE and GigaTIME (**Figure S5A; Tables S40 and S41**), two pretrained models that infer spatial protein biomarker patterns from H&E images. Datasets jointly profiling histology, spatial proteomics, and spatial transcriptomics in the same tumor tissue sections remain limited and typically include small protein panels generated using heterogeneous protocols that may compromise histological analysis. We therefore used the independent whole-transcriptome Visium HD sections as a transcript-level spatial biomarker expression reference to ensure maximum gene–protein correspondence coverage. Evaluation was restricted to non-secreted markers with unambiguous one-to-one protein–gene mappings. In total, 10 markers were retained for comparison with GigaTIME and 34 for comparison with ROSIE. Each protein model was evaluated separately, and SQUALL was restricted to the same genes as the corresponding model.

#### GigaTIME and ROSIE inference

For GigaTIME, each 224×224-pixel H&E tile was resized to the required 256×256-pixel input size and processed using the official normalization procedure. The pretrained model was applied without retraining. For each protein channel, the predicted sigmoid activation map was averaged over tissue pixels to obtain a tile-level relative protein score.

ROSIE was applied using its released pretrained model without retraining, following the official inference and preprocessing pipeline, with raw regression outputs retained for downstream analysis.

#### Multiscale spatial comparison

Spatial comparisons were conducted at 224×224 and 448×448-pixel scales to assess whether spatial correspondence was preserved (**Figures S5C and S5D; Tables S42 and S43**). At the 224×224-pixel scale, one prediction value was derived for each input tile. At the 448×448-pixel scale, spatially adjacent 2×2 groups of tiles were aggregated. Visium HD reference expression were aggregated over the same physical regions and evaluated.

#### Evaluation metrics

##### Pearson correlation coefficient

For each gene, quantile-normalized predicted and measured values at the *N* valid spatial locations were represented by vectors *x* and *y*. Pearson correlation coefficient (PCC) was calculated as

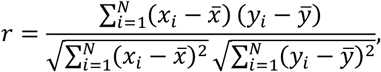

##### Area under the receiver operating characteristic curve

Following Path2Space’s evaluation, AUC was used to quantify whether a model distinguished relatively high-from low-expression regions. For each gene, quantile-normalized measured values were dichotomized at 0.5:

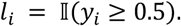

The corresponding predicted values *x_i_* were retained as continuous scores, and gene-level AUC was calculated as

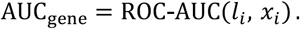

##### Structural similarity index measure

SSIM was used as a complementary measure of local spatial agreement:

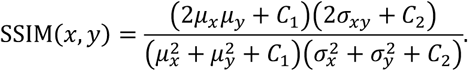

##### Gene- and section-level aggregation

Gene-level comparisons were performed using the same genes shared by all models within each comparison. Section-level performance was calculated by averaging valid gene-level metrics within each section. Cancer-level summaries were derived from the corresponding section-level results. Comparisons between SQUALL and each comparator were performed using two-sided paired Wilcoxon signed-rank tests across matched genes or sections, as specified for each analysis. Performance was additionally evaluated across different numbers of highly variable genes. Highly variable genes were ranked from the measured expression data within the relevant evaluation dataset, and the same ranked subsets were applied to all models.

#### Association between H&E characteristics and prediction performance

We investigated whether section-level variation in H&E staining and image quality was associated with virtual expression profiling performance across the 117 external Visium sections. For each section and model, overall prediction performance was summarized as the mean valid gene-wise PCC. Each H&E image was divided into occupied 224×224-pixel tiles according to the coordinates of in-tissue Visium spots. Pixels with mean normalized RGB intensity below 0.95 were classified as tissue, and tiles containing fewer than 100 tissue pixels were excluded. Five section-level image characteristics were quantified: 1. mean pixel intensity; 2. mean hematoxylin concentration; 3. mean eosin concentration; 4. mean within-tile pixel variance; and 5. mean image sharpness (**Figures S4G and S4H**).

Hematoxylin and eosin concentrations were estimated separately for each section using Vahadane stain decomposition based on sparse non-negative matrix factorization. Image sharpness was quantified as the variance of the grayscale Laplacian over tissue pixels. Pixel intensity and stain concentrations were aggregated across tiles using tissue-pixel counts as weights. RGB variance and sharpness were averaged across retained tiles. Pearson correlation was calculated between each image characteristic and mean prediction PCC. Two-sided *P* values were reported.

### WHOLE-SLIDE VIRTUAL MOLECULAR PROFILING IN TCGA

The pretrained SQUALL was applied to diagnostic H&E whole-slide images (**Figure 2J**). These slides were generated from FFPE tissue. This analysis did not include matched spatial-transcriptomics measurements. Publicly available tumor-region annotations were obtained for TCGA-CESC and converted into polygon representations aligned with the whole-slide image.

#### Whole-slide inference and tile-level expression

Each whole-slide image was processed at 20× magnification and divided into non-overlapping 224×224-pixel tiles. For each tile, the frozen SQUALL model generated a 56×56 spatial prediction map for each of the 15,757 genes. Spatial predictions were averaged within each tile to obtain one tile-level value per gene. Each slide contained approximately 10,000 analyzed tiles.

For gene *g*, tile-level values were ranked within each slide and transformed using an inverse-normal rank transformation:

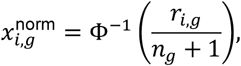

where *r_i_*_,*g*_ denotes the rank of tile *i* among all *n_g_* eligible tiles within the same sample, and Φ^−1^ denotes the inverse cumulative distribution function of the standard normal distribution.

#### Survival analysis

Slide-wide virtual gene and molecular-program scores were evaluated for association with overall survival using multivariable Cox proportional-hazards regression. Continuous molecular features were standardized across patients before model fitting. Models were adjusted for age, tumor stage, and tumor fraction where available. Hazard ratios, confidence intervals, and Wald-test *P* values were presented for each feature. Multiple-testing correction across exploratory gene-and program-level analyses was performed using the Benjamini–Hochberg false-discovery-rate procedure. This analysis was designed to evaluate the clinical relevance of cohort-scale virtual molecular profiling rather than gene-expression reconstruction accuracy.

#### Spatially resolved regional virtual expression analysis

Previously reported cervical cancer-associated genes and transcription-factor programs were first evaluated to determine whether SQUALL recovered their expected spatial distributions in whole-slide images (**Figures 2L-2P and S5F-S5H**).

For each patient, molecular features were summarized in three ways: 1. across all eligible tiles in the slide; 2. across tumor-annotated tiles; and 3. across non-tumor tiles.

The within-tumor rank-expression feature for gene or program *g* was defined as

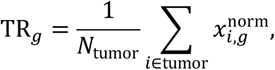

where *N*_tumor_ denotes the number of tumor tiles. Tumor fraction was defined as the number of tumor tiles divided by the total number of eligible tiles.

Region-resolved survival associations were evaluated using univariable or multivariable Cox regression, with adjustment for age and tumor stage in multivariable analyses. Region-specific molecular features were standardized across patients before regression. The prognostic values of tumor-restricted features were compared with the corresponding whole-slide and non-tumor features to determine whether spatial localization provided information beyond overall predicted abundance.

### SPATIALLY AWARE CLUSTERING

We evaluated whether representations generated by the frozen SQUALL multimodal encoder captured spatially organized tissue tumor microenvironment clusters with biological and clinical relevance. The analysis comprised two settings: 1. reference-based comparison against pathologist-defined spatial clusters; 2. evaluation of model generalizability on unseen sections through blinded assessment by board-certified pathologists.

#### Reference-based evaluation datasets

For reference-based evaluation of spatially aware clustering, expert reference annotations were generated for 44 sections spanning eight human tissues and including both tumor and non-tumor normal tissue contexts. Sections with detailed annotations supplied by the original studies were incorporated where appropriate. Histological reference annotations were additionally compared with the spatial expression of established anatomical or pathological marker genes as a quality-control procedure (**Figure 3B; Figure S6B; Tables S44 and S46**). Three experienced pathologists independently reviewed the H&E images and annotated anatomically and pathologically interpretable clusters. When necessary, corresponding marker expression patterns were provided as additional reference to assist cluster annotation. The resulting pathologist-derived reference cluster annotation were aligned with the spatial transcriptomics coordinates, and each in-tissue sequencing spot was assigned the corresponding reference cluster label.

These sections were included in SQUALL pretraining, although the pathologist-derived reference clusters were not used during pretraining, allowing us to assess the quality of representations learned through multimodal pretraining. Notably, these datasets were publicly available before the release of the pretrained OmiCLIP and STPath weights, and potential overlap with the pretraining data of these models cannot be fully excluded. We kept all pretrained models frozen and applied them according to their reported spatially aware clustering protocols, whereas other methods were trained, fine-tuned, or applied separately to each dataset following the procedures described in their original studies.

#### SQUALL inference

The pretrained SQUALL model was used as a frozen feature extractor. For each Visium spot, the corresponding H&E region and spatial gene-expression profile were processed by the histology and transcriptomics branches of the SQUALL encoder, respectively. The modality-specific representations were concatenated to form a unified multimodal representation. For each spot, given that the spatial resolution of a spot is typically larger than that of a single token and considering the limited input window size of Vision Transformers (ViT), we developed a spot-averaging method to resample the feature map and obtain a robust representation for each spot. Specifically, for each tissue section, we first partitioned it into overlapping tiles with a one-token stride and passed them through the frozen SQUALL encoder to generate detailed token-level feature maps that capture local neighborhood interactions. We then performed spot averaging, where each token was weighted according to its interaction area with the corresponding spot. Embedding of histology image token and gene expression token of the same spatial location were concatenated to obtain a 2048-dimension vector. The resulting spot-level representations were clustered using K-means clustering (K=10) for all reference-based evaluations (**Figure 3A; Figure S6A; Table S9**).

#### Implementations of comparator methods

SQUALL was compared with Leiden clustering of measured spatial gene expression implemented in Scanpy, clustering of UNI-derived histology representations, the multimodal spatial-omics methods MISO and SpatialGlue, and the pretrained H&E–spatial transcriptomics models OmiCLIP and STPath. All evaluation datasets were publicly available before the release of the OmiCLIP and STPath pretrained weights; therefore, potential overlap between evaluation data and comparator training data could not be independently excluded. To ensure a consistent and unbiased comparison, all pretrained encoders, including SQUALL, were applied using their released parameters without annotation-guided fitting or target-cohort fine-tuning. MISO, SpatialGlue, and other dataset-specific unsupervised procedures were applied separately to each section according to their published protocols. Pathologist-derived cluster annotations were used only for final evaluation and were not used to train, tune, or select any clustering configuration.

##### Scanpy

Measured Visium gene-expression matrices were library-size normalized, log-transformed, and processed using the standard Scanpy workflow. Highly variable genes were selected, followed by principal-component analysis, construction of a nearest-neighbor graph, and Leiden clustering following default parameters. The resulting Leiden clusters were used as the expression-only baseline.

##### UNI

For each in-tissue spot, a spot-centered H&E patch corresponding to approximately the Visium spot diameter was extracted and processed using the frozen UNI image encoder. No measured gene-expression values were provided to UNI. Within each tissue section, the top 50 principal components of the features were clustered using K-means with the same settings of SQUALL.

##### MISO

MISO was applied using normalized Visium expression profiles and H&E-derived features following its released implementation. The model learned modality-specific representations and pairwise cross-modal interactions to generate an integrated spatial representation. The resulting representations were partitioned using the clustering protocol provided by MISO.

##### SpatialGlue

SpatialGlue was applied to normalized, log-transformed Visium expression profiles and H&E-derived features following its official workflow. Spatial and feature neighborhood graphs were constructed, and the model jointly learned intra-modality and cross-modality relationships using its dual-attention architecture. Spatial clusters were obtained from the integrated representation using the recommended clustering procedure.

##### OmiCLIP

Spot-centered H&E regions were extracted and padded to 224×224 pixels without magnifying the original tissue content. L2-normalized image representations were generated using the frozen OmiCLIP image encoder. Within each section, the top 50 principal components of the features were clustered using K-means with the same settings of SQUALL.

##### STPath

A 224×224-pixel H&E patch was extracted around each in-tissue Visium spot and encoded using the released GigaPath image encoder. The image representations, spatial coordinates, organ identity, and Visium technology identifier were supplied to the frozen STPath model to predict spot-level gene expression. Within each section, the 2,000 predicted genes with the highest spatial variance were standardized and reduced to 50 principal components. A 15-nearest-neighbor graph was constructed, followed by Leiden clustering at a resolution of 1.0 as reported by the original study.

Additionally, for methods in which the number of clusters can be explicitly specified as a parameter, we evaluated a range of cluster numbers to assess model consistency, with the number of clusters varying over the interval (5, 15]. Additional model details and settings are provided in **Table S45**.

#### Evaluation metrics

Agreement between predicted spatial clusters and reference annotations was primarily quantified using adjusted Rand index, normalized mutual information, and adjusted mutual information. For each metric, SQUALL was compared with each alternative method across matched tissue sections using a two-sided paired Wilcoxon signed-rank test.

##### Adjusted Rand index

Adjusted Rand index (ARI) was calculated as:

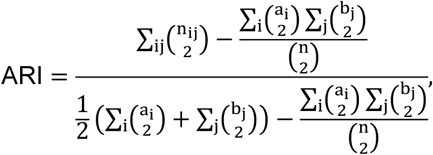

where *n_ij_* is the number of spots shared between reference cluster *i* and predicted cluster *j*, *a_i_* and *b_j_* are the corresponding marginal totals, and *n* is the total number of evaluated spots.

##### Normalized mutual information

For predicted labels *U* and reference labels *V*, normalized mutual information (NMI) was calculated as

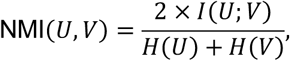

where *I*(*U*; *V*) denotes the mutual information between the proposed region labels by the model *U* and the true labels *V*, and *H*(⋅) represents Shannon entropy. A higher NMI indicates stronger agreement between proposed and reference tissue partitions.

##### Adjusted mutual information

Adjusted mutual information (AMI) corrected the observed mutual information for its expected value under random label assignments:

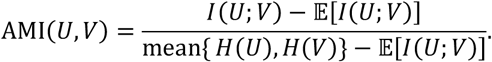

Higher AMI indicates greater agreement after accounting for chance and differences in cluster number or size.

##### CHAOS

As a supplementary metrics, CHAOS was used to measure the spatial continuity of the identified clusters. Clusters in the tumor microenvironment are not necessarily spatially continuous; therefore, CHAOS was treated as a supplementary rather than primary metric. CHAOS was defined as

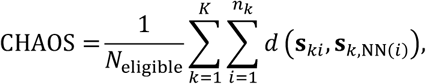

where *K* is the number of predicted clusters, *n_k_* is the number of spots in cluster *k*, **s***_ki_* denotes the standardized spatial coordinates of spot *i* in cluster *k*, and NN(*i*) denotes its nearest neighbor within the same cluster. *N*_eligible_ is the total number of spots in clusters containing more than two spots. Lower CHAOS values indicate shorter within-cluster neighbor distances and therefore greater spatial continuity.

#### Association between data characteristics and spatially aware clustering performance

We investigated whether variation in histological image characteristics or transcriptomic composition was associated with SQUALL spatially aware clustering performance across the 44 reference-annotated sections. Spatially aware clustering performance was represented by section-level ARI. Histological characteristics included section-level stain intensity, staining variability, and image-quality features calculated from occupied H&E regions. Transcriptomic characteristics included the abundance of predefined housekeeping and ribosomal gene sets (**Figures S6D and S6E**). Pearson correlation was calculated between each section-level characteristic and ARI. Two-sided *P* values were reported.

#### Differential expression and biological program analysis

Spatial gene expression matrices were normalized and log transformed. Highly variable genes were identified, and the top 2,000 genes were retained for dimensionality reduction and applicable downstream analyses. Cluster-specific differential expression was evaluated by comparing each spatial cluster with all remaining clusters using the Wilcoxon rank-sum test. Genes with a Benjamini–Hochberg-adjusted *P* < 0.05 and log fold change > 0.05 were considered significantly upregulated.

Differentially expressed genes and predefined biological gene sets were used to characterize tumor differentiation, stromal organization, immune-cell composition, vascular injury, angiogenesis, hypoxia, stress-response, and tertiary lymphoid structure-associated programs. Gene set enrichment was performed using ranked cluster-level differential-expression statistics. For comparisons of program scores between spatial clusters, two-sided Mann–Whitney *U* tests were used. Multiple comparisons were corrected using the Benjamini–Hochberg procedure where applicable.

#### Spatial cluster interaction analysis

To characterize intercellular communication between spatial clusters, ligand-receptor interaction analysis was performed across spatial proximity clusters (**Figure 3L**). Ligand-receptor interactions between spatial proximity clusters were evaluated by assessing ligand expression in the source cluster and receptor expression in the target cluster. Statistical significance of each interaction was estimated using a permutation test. Interactions with detectable ligand expression were retained, and ligand-receptor pairs were ranked according to their statistical significance for each source-target cluster combination. The top-ranked representative interactions were selected to summarize spatial cluster-to-cluster signaling relationships.

#### External validation of spatial cluster-derived gene program signatures

Spatial cluster-derived gene programs were externally evaluated using measured bulk RNA-sequencing and clinical data from TCGA-LIHC and TCGA-KIRC. The hepatocellular carcinoma c3 and c8 signatures were derived from genes characterizing the corresponding spatial tumor clusters. The hemorrhage signature was derived from genes characterizing the ccRCC hemorrhagic clusters c1 and c4. The myeloid signature was defined using the corresponding myeloid-associated marker genes (**Figures S6H, S6I, S6M, and S6N**).

#### Program signature score definition

Within each cohort, the expression value of each gene in the identified programs was converted to a percentile rank across samples. Hemorrhage and myeloid signature scores were converted to percentile ranks across samples. Each signature score was defined as the mean percentile rank of its genes. The CD8 T cell signature score was calculated from *CD3D*, *CD3E*, *CD3G*, *CD8A* and *CD8B* using *scanpy.tl.score_genes* function.

### Correlation analysis

Associations between signature scores were calculated using two-sided Spearman rank correlation tests implemented with the R function *stats::cor.test*. In TCGA-LIHC, the c3/c8 niche signature scores were correlated with the CD8 T cell signature score. In TCGA-KIRC, the hemorrhage score was correlated with myeloid cell signature score.

#### Survival analysis

Survival analyses were performed in R using *survival::Surv*. Survival curves were plotted using survminer::ggsurvplot, with 95% confidence intervals.

#### Blinded pathologist reader evaluation on external datasets

To evaluate whether SQUALL can reliably perform spatially aware clustering to generate clinically and biologically meaningful spatial cluster partitions on unseen data, we conducted a generalizability evaluation using data not included during SQUALL pretraining (**Figures 3M and 3N; Figure S6O; Tables S47 and S48**).

Five board-certified pathologists, each with at least five years of diagnostic experience, independently reviewed the annotation results generated by pathological annotation methods, including SQUALL and other competing approaches. The experts were blinded to model identities and evaluated each sample independently without interacting with the models or with one another. For each sample, the pathologists scored the clustering partitions according to their clinical interpretability and preference. Scores were assigned on a 7-point annotation-quality rank scale (1 = excellent, 7 = poor) and aggregated per model.

For pairwise comparisons between SQUALL and each competing method, we defined ‘Win’ when the expert score for SQUALL was lower and ‘Loss’ when the SQUALL score was higher than the comparator score. The Mean Opinion Score (MOS) for each method was computed as the average rating across all pathologists and samples:

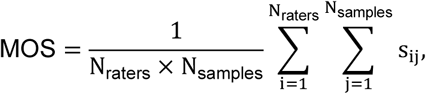

where *s_ij_* denotes the score assigned by rater *i* to sample *j*.

We further calculated the pooled pathologist win rate for each comparator method as the proportion of pathologist–section evaluations in which SQUALL received a lower (better) rank, pooling evaluations across all raters.

### IDENTIFICATION OF BREAST CANCER PROGRESSION-ASSOCIATED TRAJECTORY

All breast sections were selected from the *histMol*-low corpus and underwent rigorous metadata cleaning. Each case’s pathological characteristics were manually rechecked based on histology images and corresponding pathological reports, if available, from the original publications. To mitigate platform-specific artifacts that could hinder biological discovery, we used Visium data for primary discovery analyses, reserving ST data for validation (**Figure 4A; Figures S7A and S7G; Table S49**).

#### PAM50 subtype assignment

Raw expression data of individual spots from the spatial transcriptomics (ST) dataset were aggregated into pseudobulk RNA expression profiles per section. The resulting pseudobulk expression matrix was log₂-transformed with a pseudocount of 1 to stabilize variance.

For each gene g, the expected expression level across all samples was estimated as the arithmetic mean 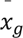 and gene-wise mean normalization was applied by transforming each observation as 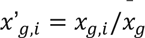. The normalized matrix was then linearly rescaled using *genefu::rescale*, which projects gene-wise expression values onto a common [0,1] range to ensure comparability across genes during centroid correlation. PAM50 molecular subtypes were subsequently inferred using *genefu::molecular.subtyping (sbt.model=“pam50”)* based on the rescaled expression matrix and the pretrained PAM50 classification centroids. For each pseudobulk sample, the procedure returned the predicted intrinsic subtype: Luminal A, Luminal B, HER2-enriched, Basal-like, or Normal-like, along with centroid correlation scores and subtype probabilities, where applicable.

To evaluate classification reliability, PAM50-assigned subtypes were compared with curated clinical molecular subtype annotations, and concordance was assessed using Cohen’s kappa (κ) statistic (**Figure S7F**).

#### Pathological region enrichment analysis

For each spot, we obtained a multimodal representation using the procedure described for spatially aware clustering. The embeddings were then clustered using K-means. The final cluster number was selected based on trade-off to capture enough between-group diversity while not over-fragmenting the biological groups. After clustering the spot embeddings into recurrent groups, we assessed cluster enrichment across pathological regions and subtypes using the corresponding annotations.

To assess whether SQUALL-derived spot clusters were preferentially associated with specific pathological subtypes or spatial regions, we performed one-versus-all enrichment analysis using odds ratios (ORs) and two-sided Fisher’s exact tests at the spot level. For each cluster, spots assigned to that cluster were compared with all remaining spots to evaluate enrichment across each pathological subtype or region label. A 2×2 contingency table was constructed for each comparison, with rows representing the presence or absence of the target subtype/region label and columns representing cluster assignment versus all other clusters.

The OR quantified the strength and direction of association, with OR > 1 indicating enrichment of a cluster within the specified subtype or region. Statistical significance was assessed using two-sided Fisher’s exact tests, followed by Benjamini–Hochberg false discovery rate (FDR) correction for multiple comparisons.

#### Diffusion trajectory analysis

Following the enrichment analysis, clusters that were (i) significantly enriched within tumor regions and (ii) exhibited enrichment patterns across different pathological subtypes were selected for diffusion trajectory analysis. Specifically, clusters significantly enriched in tumor regions and spanning diverse breast pathologies were cluster 8 (OR: MpBC = 5.77, FDR < 10^-300^; IDC = 1.12, FDR = 1.74×10^-4^), cluster 10 (IDC = 2.84, FDR < 10^-300^; MpBC = 1.72, FDR = 2.33×10^-52^), cluster 15 (MpBC = 1.40, FDR = 6.58×10^-52^; IDC = 1.80, FDR = 4. 54×10^-56^), cluster 26 (DCIS = 277.70, FDR < 10^-300^; IDC=2.84, FDR < 2.84×10^-50^), cluster 33 (DCIS = 6.41, FDR < 10^-300^; IDC=40.20, FDR < 10^-300^), and cluster 34 (MpBC = 1.28, FDR = 1.43×10^-16^; ILC = 2.20, FDR = 6.30×10^-40^).

Principal component analysis (PCA) was first applied for dimensionality reduction, and the top 32 principal components were used to construct a diffusion graph capturing the manifold of pathogenesis. Diffusion components were then computed using Scanpy (*sc.tl.diffmap, n_comps=10*), yielding a low-dimensional representation that reflects the continuum of progression states among tumor spots. The continuous pathological progression order of breast cancer subtypes along the diffusion manifold was inferred by diffusion pseudotime (DPT) analysis implemented in Scanpy (*sc.tl.dpt, n_dcs=4*).

#### Correspondence of ST data to Visium data

To map the ST dataset to Visium dataset, human breast cancer ST sections were first encoded using the SQUALL encoder, following the same preprocessing and feature extraction procedure applied to the Visium data. The resulting ST embeddings were subsequently clustered using the same strategy as described above for the Visium dataset, ensuring comparable cluster granularity across platforms. To quantitatively assess the correspondence between ST and Visium-derived clusters, we computed the cosine similarity between the feature embeddings of each ST spot and those of all Visium spots. For each ST cluster, the median cosine similarity was calculated against every Visium cluster. The Visium cluster with the highest median similarity was then assigned as the corresponding counterpart for that ST cluster, defining a robust cross-platform mapping at the cluster level.

#### Survival analysis

Survival data and gene-by-count expression matrix of TCGA-BRCA cohort were downloaded from the UCSC Xena. Genes were ranked according to their Pearson correlation coefficient with Visium pseudotime, and signatures were defined using the top or bottom correlated genes. A weighted risk score was calculated for each sample as the sum of gene expression values multiplied by the corresponding signed correlation coefficients. Patients were stratified into high- and low-risk groups. Survival analysis was performed using the R package survival (v3.8.3).

### IDENTIFICATION OF OVARIAN TUMOR RECURRENCE-ASSOCIATED SPATIAL REGIONS

All ovarian tumor sections were selected from *histMol*-high corpus and underwent rigorous metadata cleaning (**Figures 5A-5C; Figures S8C and S8D; Table S52**). Each case’s pathological characteristics were manually rechecked based on histology images and corresponding pathological reports. To obtain a regional (112 um) summarization of tumor microenvironments, for each transcriptomics-histology image tiles’ token sequence, we prepended a constant [CLS] token and passed the complete token sequence through the frozen SQUALL encoder. Subsequently, the final [CLS] tokens representation for each tile was used for discoveries. After hierarchical clustering, clinical endpoint and pathological region labels were used for enrichment analysis.

#### Cell type segmentation and classification

CellViT++, using SAM-L as its backbone architecture, was employed for simultaneous cell segmentation and classification from H&E histology images. The model was trained to categorize individual cells into five major histopathological classes: connective, inflammatory, neoplastic, epithelial, and dead cells. Following segmentation, the classified cell masks were aggregated within each spatial tile to quantify cell-type composition and to calculate proportional differences across tissue regions. These spatially resolved cellular distributions were subsequently used for downstream spatial correlation and phenotypic association analyses (**Figures 5F and 5G**).

### CLINICAL OUTCOME PREDICTION USING SQUALL IMAGE REPRESENTATIONS

We evaluated the pretrained SQUALL histology encoder in the independent TCGA and CGODC cohorts to assess its ability to predict clinical outcomes directly from histological images, without reconstructing molecular profiles. All whole-slide images were processed at 20× magnification to ensure a consistent spatial scale across cohorts (**Tables S56-S59**).

#### Histological imaging of the CGODC-OV cohort

Formalin-fixed, paraffin-embedded (FFPE) clinical tissue sections were collected from pathology archives following institutional ethical approval and standard biospecimen handling protocols. All samples were sectioned at a thickness of 4–5 µm, mounted on glass slides, and stained with hematoxylin and eosin (H&E) according to routine clinical procedures.

Slides were digitized using a whole-slide scanner (Olympus SLIDEVIEW VS200) at 40× optical magnification, corresponding to a spatial resolution of approximately 0.25 µm per pixel. The resulting whole-slide images (WSIs) were stored in high-resolution digital format (TIFF) for downstream computational analyses.

All FFPE samples were derived from clinically annotated cases, ensuring the availability of associated diagnostic and prognostic metadata such as tumor grade, histological subtype, and clinical outcomes.

#### Survival prediction

For survival prediction, we adopted the ABMIL framework to aggregate tile-level embeddings into a patient-level representation. The aggregated embedding was mapped to a scalar log-risk score, which was optimized under the Cox proportional hazards model. Specifically, for each patient with observed survival time t*_i_*, event indicator *δ_i_*, and predicted log-risk l*_i_*, the objective was to maximize the Cox partial likelihood

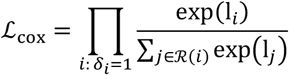

where ℛ(*i*) denotes the risk set of patients still under observation at time t*_i_*. Model training minimized the negative log-partial likelihood. Model development and evaluation were carried out using a 5-fold cross-validation scheme. Predictive performance was quantified using C-index, and the learned attention weights also provided interpretability by highlighting histologic regions most influential in shaping patient-level risk estimates.

#### Platinum-based chemotherapy resistance prediction

Platinum sensitivity was defined according to the interval between completion of platinum-based chemotherapy and documented disease progression. Patients with progression more than 6 months after completion of platinum-based therapy were classified as platinum-sensitive, whereas those with progression within 6 months were classified as platinum-resistant. We curated patient-level platinum response information from both the CGODC-OV and TCGA-OV cohorts. For each cohort, patients were randomly partitioned into five folds for patient-level cross-validation. Models were trained and evaluated using an ABMIL framework with cross-entropy loss, and classification performance was assessed using accuracy, AUROC, and F1 score. For patients with available time-to-recurrence information, recurrence prediction was additionally evaluated. Recurrence time was defined as the interval from initial treatment to the first documented recurrence event, and predictive performance was quantified using C-index.

#### Benchmarking against other foundation models

For comparative analyses with other pathological foundation models, we implemented feature extraction using open-source frameworks: CLAM (https://github.com/mahmoodlab/CLAM) for UNI, and modified implementations of PLIP and Virchow based on the timm library (https://huggingface.com) using corresponding methods’ reported protocols. Pretrained parameters for the benchmark models were obtained from the Hugging Face Model Hub including PLIP (https://huggingface.co/vinid/plip), UNI(https://huggingface.co/MahmoodLab/UNI), and Virchow (https://huggingface.co/paige-ai/Virchow). All models were trained using the same protocol as for SQUALL (**Tables S54 and S55**).

#### Evaluation of SQUALL-derived risk scores with clinical variables

To determine whether the model-derived risk stratification provided prognostic information independent of established clinical factors, multivariable Cox proportional-hazards analyses were performed separately in the TCGA-CESC and CGODC-OV cohorts. Cross-validation results were deduplicated to retain one out-of-fold prediction per patient. Risk scores generated by SQUALL, UNI, PLIP and Virchow were each dichotomized into low- and high-risk groups using the corresponding model-specific optimal cutoff. For each model, a separate multivariable Cox model was fitted by including its predicted risk group together with patient age and clinical stage. Clinical stage was treated as a categorical variable, with the lowest stage represented in the corresponding cohort used as the reference category. An L2 penalization parameter of 0.05 was applied to improve estimation stability. The adjusted hazard ratio for each model-derived high-risk group, together with its 95% confidence interval and Wald test *P* value, was presented. The four model-derived risk groups were not entered simultaneously into the same Cox model; instead, each was evaluated in an independently fitted model using the same clinical covariates. A hazard ratio greater than 1 indicated a higher event hazard in the predicted high-risk group relative to the corresponding low-risk group. This analysis enabled direct comparison of the independent prognostic value of risk stratification derived from SQUALL, UNI, PLIP and Virchow after adjustment for age and clinical stage.

#### Risk stratification evaluation

We integrated the five-fold cross-validation results. Predicted risk scores were robustly normalized across folds and then merged into a single dataset. The optimal cutoff value for the normalized risk scores to stratify patients was determined. Based on this dichotomization, we calculated hazard ratios with corresponding 95% confidence intervals and performed Kaplan-Meier survival analyses to visualize survival differences between groups. Statistical significance of the separation was assessed, and the resulting Kaplan-Meier curves with annotated HRs provided an interpretable summary of the prognostic value of the predicted risk scores.

#### Correlation of SQUALL-derived risk scores with RNA expression

For correlation analysis between model-predicted survival risk and bulk RNA-seq expression profiles, we restricted TCGA bulk RNA-seq data to primary tumor samples (**Figure 5J**). Replicate libraries from the same patient were averaged to obtain patient-level expression profiles. Raw counts were log-transformed, and gene expression values were further normalized within each patient by dividing each gene’s expression by the total expression across all genes in that patient, yielding relative expression abundances. The resulting matrix was merged with normalized survival risk scores. For each gene, association with risk was quantified using the Pearson correlation coefficient across patients. Genes were ranked by correlation, and the top and bottom 5% (positively vs. negatively associated) were selected for visualization and pathway enrichment analysis. To visualize cohort-level gene expression trends along the model-predicted risk, patients were ordered by their risk scores.

#### Pathological inspection of attention heatmap

Tile-level attention scores were derived from the SQUALL-feature-trained ABMIL module, in which each tile’s attention weight quantifies its relative contribution to the section-level prediction. To visualize the spatial distribution of these learned attentions, we constructed section-level heatmaps by combining the spatial coordinates of all tiles with their corresponding attention weights (**Figures 5K and 5L; Figures S9A, S9C, and S9D**).

For each whole-slide image (WSI), the attention scores were min-max normalized and contrast-enhanced using a power-law transformation to emphasize regions with high discriminative importance. A continuous attention field was then generated by interpolating the normalized tile scores across the entire tissue section using a KD-tree-based nearest-neighbor interpolation strategy, which preserves spatial locality while maintaining smooth transitions between adjacent tiles. Each tile was modeled as a 224-pixel square region, and overlapping tiles were averaged to ensure continuity. Regions with insufficient tile coverage were masked to suppress boundary artifacts, and mild Gaussian smoothing was applied to further enhance spatial coherence.

The resulting attention field was blended with the original H&E image in RGBA color space with an opacity of 0.6. Regions outside the valid tissue coverage were rendered as white background. This visualization provides intuitive interpretability by highlighting morphologically relevant tissue regions that most strongly influenced the model’s predictions.

Additionally, 20× field images were randomly sampled within regions guided by the model-generated attention heatmaps. Each field was obtained at the same magnification and from comparable tissue regions. Mitotic figures were quantified and pathological structures were annotated using QuPath, with all measurements independently reviewed and adjudicated by two pathologists (**Figure 5K; Figures S9E and S9F**).

### OVER-REPRESENTATION ENRICHMENT ANALYSIS

Functional enrichment of the gene sets was performed using both Gene Ontology (GO) and MSigDB Hallmark gene sets. For GO, the Biological Process (BP) ontology for *Homo sapiens* was used as the reference annotation. For MSigDB, curated Hallmark pathways were employed to evaluate enrichment of core transcriptional programs. Over-representation significance was evaluated using a hypergeometric model, and multiple testing correction was performed with the Benjamini–Hochberg procedure. Enrichment results were ranked according to adjusted *P* values and enrichment ratios and visualized as bar plots displaying − log_10_ adjusted *P* values for selected pathways. The same procedure was applied to analyze genes correlated with the prognostic-associated trajectory in breast cancer. For the enrichment analysis of top differentially expressed genes in the TAIE region, only GO Biological Process enrichment was performed. For all analyses, only the top enriched nonredundant terms were shown.

### QUANTIFICATION AND STATISTICAL ANALYSIS

Comparisons of matched model performance were performed using two-sided paired Wilcoxon signed-rank tests. Two-group comparisons used two-sided Mann–Whitney U tests, whereas comparisons involving more than two groups used Kruskal–Wallis tests. Categorical enrichment was assessed using two-sided Fisher’s exact tests. Continuous-variable associations were evaluated using Pearson’s or Spearman’s correlation, as appropriate. Differential expression was assessed using Wilcoxon rank-sum tests. Survival analyses used Cox proportional-hazards models, Kaplan–Meier curves, Wald tests, and log-rank tests. Gene-set over-representation and ligand–receptor significance were evaluated using hypergeometric and permutation tests, respectively. Multiple testing was corrected using the Benjamini–Hochberg procedure. Unless otherwise specified, tests were two-sided and *P* < 0.05 was considered statistically significant.

## Notes

### Summary of Updates

Updated benchmarking. All figures and text revised; author affiliations updated; Supplemental files updated.

