## Supplemental Tables for "A multimodal foundation model to integrate histology with spatial molecular programs"

### Supplementary Tables

| Res. | Data source | Hosting country | Num. study | Num. section | Num. sample |
|---|---|---|---|---|---|
| Low | Gene Expression Omnibus | U.S. | 235 | 2,228 | 1,388 |
| | ArrayExpress | U.S. | 16 | 218 | 102 |
| | Zenodo | Switzerland | 15 | 162 | 100 |
| | Mendeley | Netherlands | 9 | 205 | 56 |
| | SOAR | U.S. | 9 | 69 | 61 |
| | CELLxGENE | U.S. | 7 | 147 | 91 |
| | National Genomics Data Center | China | 4 | 37 | 17 |
| | Figshare | U.K. | 3 | 38 | 29 |
| | Single Cell Portal | U.S. | 3 | 25 | 12 |
| | Dryad | U.S. | 1 | 28 | 28 |
| | Spatial Research | Sweden | 1 | 16 | 2 |
| | Human Cell Atlas | U.S. | 1 | 8 | 8 |
| | KU Leuven RDR | Belgium | 1 | 6 | 6 |
| | STOmics | China | 1 | 2 | 2 |
| | 10X Genomics | U.S. | - | 54 | 50 |
| High | STAGE | China | 1 | 99 | 99 |
| | In house | China | - | 95 | 76 |
| | SPATCH (OV, COAD) | China | 1 | 4 | 2 |
| | 10X Genomics | U.S. | - | 5 | 4 |
| Total | Low resolution data | | - | 3,243 | 1,952 |
| | High resolution data | | - | 203 | 181 |
| | Total | | | 3,446 | 2,133 |

**Table S1**: **Dataset source characteristics of *histMol* corpus.** The *histMol* corpus consists of both low-resolution (*histMol*-low, with histology images acquired at $< 20\times$ magnification) and high-resolution (*histMol*-high, with histology images acquired at $20\times$ magnification) data for a two-stage pretraining strategy. All included sections contain paired hematoxylin and eosin (H&E)-stained pathology images and corresponding spatial transcriptomic profiles. We combined in-house data with datasets systematically collected from 15 publicly accessible repositories, covering diverse tissue types, cohorts, and geographical sources, to construct the pretraining corpus.

| Res. | Platform | Array pattern | Spot diameter | Inter-Cen Dist. | Num. sections |
|---|---|---|---|---|---|
| Low | Visium | Hexagonal | 55 µm | 100 µm | 2,555 |
| | Ex-ST | Hexagonal | 20 µm | 37 µm | 4 |
| | Spatial Transcriptomics | Square | 100 µm | 200 µm | 606 |
| | SM-Omics | Square | 100 µm | 100 µm | 15 |
| | CBSST-seq | Square | 50 µm | 100 µm | 1 |
| | Decoder-seq | Square | 50 µm/25 µm/15 µm | 50 µm/25 µm/15 µm | 22 |
| Low* | HDST | Hexagonal | 2 µm | 3.464 µm | 6 |
| | OpenST | Hexagonal | 0.3 µm | 0.625 µm | 24 |
| | NovaST | Hexagonal | 0.3 µm | 0.625 µm | 7 |
| | Seq-Scope | Hexagonal | 0.3 µm | 0.625 µm | 1 |
| | Stereo-seq | Square | 0.22 µm | 0.5 µm | 2 |
| High | Visium HD | Square | 2 µm | 2 µm | 203 |

**Table S2**: **Technical characteristics of the platforms represented in the *histMol* corpus.** The *histMol* corpus consists of data sourced from various technical platforms with varying sequencing resolutions and spot arrangement patterns. For the low-resolution asterisk (*) subset, although the sequencing resolution is higher than that of the low-resolution data without asterisk, it either lacks high-quality pathological images or is limited in data size. Therefore, we first downsampled the histological images in this subset while preserving aspect ratio to align with the image dimensions typical of the non-asterisked subset (i.e., a maximum of 2,000 pixels on the longest side). Next, we applied grid binning to achieve the target token resolution, where one binned sequencing unit corresponds to a $4\times 4$ image pixel area, without using pseudolabeling. Low and Low* are collectively termed *histMol*-low. The high-resolution subset consists solely of Visium HD data, selected for its high-quality histology images and fine-grained sequencing resolution.

| Resolution | Species | Num. sections | Num. samples |
|---|---|---|---|
| Low | Homo sapiens | 1,926 | 1,198 |
| | Mus musculus | 1,282 | 721 |
| | Homo sapiens/Mus musculus xenograft | 35 | 33 |
| High | Homo sapiens | 203 | 181 |

**Table S3**: **Species characteristics of *histMol* corpus.** The *histMol* corpus comprises samples from Homo sapiens, Mus musculus, and xenograft models. For each species, only genes with nonzero expression in at least 1 sample were retained. The final gene list was then intersected with the Visium HD probe list to obtain a unified gene list for the *histMol* corpus. In total, 15,757 genes were obtained.

| Species | # Sections | # Samples | Histology median resolution | Sequencing median resolution |
|---|---|---|---|---|
| Human | 1,926 | 1,198 | 3.56 $\mu$m/pixel | 55 $\mu$m |
| Mouse | 1,282 | 721 | 3.92 $\mu$m/pixel | 55 $\mu$m |
| Xenografts | 35 | 33 | 3.80 $\mu$m/pixel | 55 $\mu$m |

**Table S4**: **Species composition and resolution characteristics of *histMol*-Low corpus.** H&E image resolutions were summarized across collected datasets. Spatial transcriptomic resolutions are primarily determined by the experimental platform and are summarized here for reference; detailed platform-level resolution information is provided in **Table S2**. All newly generated high-resolution spatial transcriptomic data were profiled using the Visium HD platform with 2-$\mu$m binning, and corresponding H&E images were standardized to 0.5 $\mu$m/pixel for SQUALL pretraining.

| Resolution | Preservation method | Num. sections | Num. samples |
|---|---|---|---|
| Low | Fresh Frozen | 2,818 | 1,628 |
| | FFPE | 385 | 308 |
| | Fixed Frozen | 40 | 23 |
| High | Fresh Frozen | 2 | 2 |
| | FFPE | 201 | 179 |

**Table S5**: **Characteristics of tissue preservation methods of the *histMol* corpus.** Tissue preservation methods were confirmed based on the experimental conditions or the original protocol utilized in the corresponding publication, and were categorized according to the specific protocol details. Here, 'Fixed Frozen' refers to samples cryosectioned from PFA-fixed, OCT-embedded tissue blocks. Please note that, for a given patient or subject, different tissue blocks may have undergone different preservation methods before sequencing.

| Major tissue type | Num. slides (Train: Test) |
|---|---|
| Brain | 622 (560: 62) |
| Spinal cord | 384 (346: 38) |
| Skin | 272 (245: 27) |
| Liver | 254 (229: 25) |
| Breast | 246 (221: 25) |
| Lung | 214 (193: 21) |
| Intestine | 187 (168: 19) |
| Kidney | 181 (163: 18) |
| Heart | 179 (161: 18) |
| Prostate | 123 (111: 12) |
| Head and neck | 115 (103: 12) |
| Ovary | 89 (80: 9) |
| Pancreas | 87 (78: 9) |
| Lymph node | 45 (40: 5) |
| Spleen | 39 (35: 4) |
| Bone | 33 (30: 3) |
| Embryo | 32 (29: 3) |
| Adipose tissue | 31 (28: 3) |
| Placenta | 24 (22: 2) |
| Thymus | 19 (17: 2) |
| Soft tissue | 18 (16: 2) |
| Stomach | 15 (13: 2) |
| Endometrium | 10 (10: 0) |
| Muscle | 10 (10: 0) |
| Uterus | 9 (9: 0) |
| Cervix | 9 (9: 0) |
| Tonsil | 8 (8: 0) |
| Blood vessel | 7 (7: 0) |
| Peripheral nerve | 6 (6: 0) |
| Bladder | 4 (4: 0) |
| Esophagus | 4 (4: 0) |
| Tendon | 4 (4: 0) |
| Testis | 3 (3: 0) |
| Total | 3,283 (2,962: 321) |

**Table S6**: **Tissue anatomical site distribution of *histMol*-low.** Each section was systematically assigned a slide label for histology images representing a major anatomical region, informed by the original metadata and consensus of three experienced pathologists. These anatomical labels are used in a classification task aimed at evaluating pretraining effectiveness and facilitating ablation studies of key model components during Stage 1 training. For this task, sections are stratified by species, tissue embedding method, and technical platform, followed by a 90:10 (2,962: 321) train-test split. We split tissue microarray (TMA) sections into different sections for classification (n = 3). Tissues represented by fewer than 15 sections are included exclusively in the training set.

| Cancer by system | Cancer by location | Num. slides (Train: Test) |
|---|---|---|
| Breast | Breast Cancer | 233 (210: 23) |
| Neurologic | Brain Tumor–Adult | 83 (75: 8) |
| | Brain Tumor–Childhood | 30 (27: 3) |
| Genitourinary | Kidney (Renal Cell) Cancer | 67 (60: 7) |
| | Prostate Cancer | 66 (59: 7) |
| | Bladder Cancer | 4 (4: 0) |
| | Testicular Cancer | 3 (3: 0) |
| Gynecologic | Ovarian Epithelial Cancer | 51 (46: 5) |
| | Cervical Cancer | 9 (9: 0) |
| | Endometrial Cancer | 6 (6: 0) |
| Digestive/Gastrointestinal | Pancreatic Cancer | 54 (49: 5) |
| | Colon Cancer | 40 (36: 4) |
| | Liver Cancer–Adult Primary | 29 (26: 3) |
| | Liver Cancer–Childhood | 3 (3: 0) |
| | Stomach (Gastric) Cancer | 12 (12: 0) |
| | Location not specified | 1 (1: 0) |
| Respiratory/Thoracic | Lung Cancer–Non-Small Cell | 40 (36: 4) |
| Head and Neck | Lip and Oral Cavity Cancer | 35 (31: 4) |
| | Throat Cancer | 8 (8: 0) |
| | Rare | 4 (4: 0) |
| | Nasopharyngeal Cancer | 2 (2: 0) |
| | Laryngeal Cancer | 1 (1: 0) |
| Skin | Skin Cancer | 28 (25: 3) |
| | Melanoma | 1 (1: 0) |
| Musculoskeletal | Soft Tissue Sarcoma | 14 (14: 0) |
| Hematologic/Blood | Primary Central Nervous System Lymphoma | 8 (8: 0) |
| Eye | Rare | 1 (1: 0) |
| Endocrine and Neuroendocrine | Thyroid Cancer | 4 (4: 0) |
| | Pheochromocytoma | 2 (2: 0) |
| | Adrenocortical Carcinoma | 1 (1: 0) |
| | Total | 840 (764: 76) |

**Table S7**: **Cancer type distribution of *histMol*-low.** Each section was systematically assigned a slide label for histology images representing a cancer type, based on the original metadata and consensus from three experienced pathologists. Label hierarchy was inspired by the NCI cancer categories with minor modifications. These cancer type labels are subsequently used in a classification task to evaluate the impact of pretraining data composition on model performance during Stage 1 training. For this task, sections are stratified by cancer type, and a 90:10 (764: 76) train-test split is applied. Cancer types represented by fewer than 20 sections are excluded from the test set.

| Cancer type | Num. patients | Num. slides | Num. tiles |
|---|---|---|---|
| Ovarian cancer | 78 | 93 | 268,745 |
| Cervical cancer | 23 | 23 | 79,767 |
| Endometrial cancer | 20 | 20 | 68,949 |
| Liver cancer | 41 | 46 | 148,786 |
| Stomach cancer | 13 | 13 | 27,316 |
| Colon cancer | 6 | 8 | 22,204 |
| Total | 181 | 203 | 615,767 |

**Table S8**: **Cancer type distribution of *histMol*-high.** Each section was systematically assigned a cancer type label based on the original pathological reports corresponding to the patients. Only the anatomical sites of the primary tumors are reported. Tiles with less than 50% tissue area were discarded.

| Downstream application | SQUALL setting | Representation/output |
|---|---|---|
| Virtual gene expression profiling (Figure 2) | Frozen H&E encoder and ST decoder | Predicted gene expressions |
| Spatially aware clustering (Figure 3) | Frozen H&E and ST encoders | H&E–ST multimodal embeddings |
| Progression trajectory modeling (Figure 4) | Frozen H&E and ST encoders | H&E–ST multimodal embeddings |
| Recurrence-associated region discovery (Figure 5) | Frozen H&E and ST encoders | H&E–ST multimodal embeddings |
| WSI-level outcome prediction (Figure 5) | Frozen H&E encoder only | H&E-only image embeddings |

**Table S9**: **Summary of SQUALL configurations and representations used for downstream applications.** Different downstream tasks utilized task-specific SQUALL components and representations or outputs. Frozen encoders indicate that pretrained SQUALL parameters were kept unchanged during downstream analysis.

| Config. | Tissue | Cancer |
|---|---|---|
| Num. of class | 33 | 12 |
| Attention type | Gated Attention | |
| Hidden dim. | 256 | |
| Optimization target | Cross Entropy | |
| Optimizer | AdamW | |
| Optimizer momentum | $\beta_1 = 0.9$, $\beta_2 = 0.999$ | |
| Peak learning rate | 1e-4 | |
| Learning rate schedule | Cosine annealing | |
| Epochs | 100 | 50 |
| Batch size | 1 | |
| Gradient accu. steps | 64 | |

**Table S10**: **Configuration for SQUALL slide classification probing task (on *histMol*-low corpus).** An NVIDIA H100 GPU was used for ABMIL evaluation. The batch size refers to the number of slides processed in one forward pass. For each slide, all associated tiles were forwarded through the model. To stabilize the gradients and simulate large-batch training, we employed gradient accumulation, allowing multiple sections to be processed before a single backward pass.

| Pos. Embd. | Learnable? | w/o Resolution | w/ Resolution | Δ |
|---|---|---|---|---|
| PE | ✗ | 0.434 (0.368-0.501) | **0.478 (0.410-0.544)** | +4.4% |
| RPB | ✓ | 0.488 (0.419-0.553) | **0.542 (0.471-0.604)** | +5.4% |

**Table S11**: **Assessing the impact of injecting resolution as an additional parameter into the positional encoding within attention in SQUALL (on *histMol*-low corpus).** We randomly sampled 30% of *histMol*-low data tiles without replacement and employed a 100 epochs training schedule for this assessment. For position encoding (PE), we used sinusoidal positional encoding. For relative position bias (RPB), we took the relative distance between 2 tokens $(\Delta x, \Delta y)$ as input to a two layers MLP. To inject resolution, for PE, the token before it was passed to the attention blocks were modeled as $(X_0 = X + PE \times Resolution)$; for RPB, the q-k dot product within attention between token $X_i$ and $X_j$ was calculated as $\frac{1}{\sqrt{d}}(U_i W^q)(V_j W^k) + B_{ij} \times Resolution$. Pre-extracted tile features from each section were used to train a shallow ABMIL aggregation and classification head on curated tissue classification train-test folds (2,962: 321). Test performance was reported using top-1 accuracy. The best-performing metrics are highlighted in bold, with 95% confidence intervals provided in parentheses. Baseline settings are marked in grey. Improvements in performance are indicated in the Δ column. All experiments were performed under SQUALL-large settings.

| Cat. | Top1-ACC | Top3-ACC | Top5-ACC | Weighted F1 | AUROC |
|---|---|---|---|---|---|
| No | 0.595 (0.523-0.655) | 0.776 (0.710-0.823) | 0.859 (0.803-0.898) | 0.578 (0.509-0.641) | **0.907 (0.858-0.938)** |
| PE | 0.659 (0.591-0.718) | 0.805 (0.746-0.853) | **0.902 (0.852-0.934)** | 0.632 (0.561-0.691) | 0.914 (0.863-0.941) |
| RPB | 0.542 (0.471-0.604) | 0.732 (0.665-0.784) | 0.810 (0.751-0.857) | 0.500 (0.433-0.567) | 0.872 (0.819-0.910) |
| SQUALL | **0.663 (0.596-0.723)** | **0.815 (0.756-0.861)** | 0.873 (0.819-0.910) | **0.635 (0.566-0.696)** | 0.901 (0.852-0.934) |

**Table S12**: **Assessing the impact of position encoding strategy of SQUALL (on *histMol*-low corpus).** We randomly sampled 30% of *histMol*-low data tiles without replacement and employed a 100 epochs training schedule for this assessment. We evaluated various position encoding strategies. 'SQUALL' here refers to the combination of position embedding and relative position bias. Pre-extracted tile features from each section were used to train a shallow ABMIL aggregation and classification head on curated tissue classification train-test folds (2,962: 321). Experiments were performed under SQUALL-large settings. Test performance was reported using top-1, top-3, and top-5 accuracy, weighted F1 score, and AUROC. The best and second-best metrics are bolded and underlined, with the 95% confidence interval provided in parentheses. Baseline settings are marked in grey. Final position encoding strategy was selected based on top-1 accuracy in the presented study.

| P.E. | Cat. | Top1-ACC | Top3-ACC | Top5-ACC | Weighted F1 | AUROC |
|---|---|---|---|---|---|---|
| No | – | .595 (.523-.655) | **.776(.710-.823)** | **.859 (.803-.898)** | .578 (.509-.641) | .907 (.858-.938) |
| Sinusoid | Abs. | .478 (.410-.544) | .683(.615-.740) | .790 (.725-.836) | .405 (.341-.472) | .829 (.767-.869) |
| RoPE-2D | Hyb. | .546 (.475-.609) | .746(.680-.797) | .839 (.782-.882) | .494 (.424-.558) | .893 (.841-.926) |
| Parameters | Rel. | .498 (.428-.562) | .702(.635-.758) | .795 (.731-.840) | .433 (.363-.496) | .878 (.825-.914) |
| Buckets (32) | Rel. | .590 (.518-.650) | .751(.685-.802) | .849 (.793-.890) | .553 (.485-.618) | **.910 (.863-.941)** |
| Buckets (16) | Rel. | **.634 (.566-.696)** | .761(.695-.810) | .849 (.793-.890) | **.605 (.537-.668)** | .907 (.858-.938) |

**Table S13**: **Assessing the impact of position embedding type of SQUALL (on *histMol*-low corpus).** We randomly sampled 30% of *histMol*-low data tiles without replacement and employed a 100 epochs training schedule for this assessment. We systematically evaluated various type of positional encoding. All position encoding presented here were all injected with resolution. 'Parameters' here refers to for all possible relative position values between tokens, we set their corresponding relative position bias as the model parameters. The 'bucket method' refers to uniformly partitioning the set of all possible relative distances into equal-frequency intervals—a strategy we found can improve model performance compared with direction partition (data not shown). Experiments were performed under SQUALL-large settings. Pre-extracted tile features from each section were used to train a shallow ABMIL aggregation and classification head on curated tissue classification train-test folds (2,962: 321). Test performance was reported using top-1, top-3, and top-5 accuracy, weighted F1 score, and AUROC. The best and second-best metrics are bolded and underlined, with the 95% confidence interval provided in parentheses. Leading zeros before decimal points were omitted to enhance readability. Baseline setting are marked in grey.

| Arch. | w/ Stain Norm | w/o Stain Norm | Δ |
|---|---|---|---|
| SQUALL-base | 0.571 (0.499-0.632) | **0.639 (0.571-0.700)** | −6.8% |
| SQUALL-large | 0.654 (0.586-0.714) | **0.663 (0.596-0.723)** | −0.9% |

**Table S14**: **Assessing the impact of stain normalization of data (on *histMol*-low corpus).** We randomly sampled 30% of *histMol*-low data tiles without replacement and employed a 100 epochs training schedule for this assessment. Comparison of performance with and without stain normalization for tissue label classification on *histMol*-low slides. For stain normalization, we utilized Macenko's method, using the median color vector of the entire *histMol*-low histology slides as the target stain vector. Each tissue section was then normalized to this target vector. The models were trained on the entire training set. Pre-extracted tile features from each section were used to train a shallow ABMIL aggregation and classification head on curated tissue classification train-test folds (2,962: 321). Test performance was reported using top-1 accuracy, with the 95% confidence interval provided in parentheses. Changes in performance are indicated in the Δ column.

| %Mask | Top1-ACC | Top3-ACC | Top5-ACC | Weighted F1 | AUROC |
|---|---|---|---|---|---|
| 25% | 0.624 (0.557-0.687) | 0.771 (0.705-0.819) | 0.839 (0.782-0.882) | 0.592 (0.523-0.655) | 0.897 (0.847-0.930) |
| 50% | **0.663 (0.596-0.723)** | **0.815 (0.756-0.861)** | **0.873 (0.819-0.910)** | **0.635 (0.566-0.696)** | **0.901 (0.852-0.934)** |
| 75% | 0.639 (0.571-0.700) | 0.751 (0.685-0.802) | 0.844 (0.787-0.886) | 0.593 (0.523-0.655) | 0.872 (0.819-0.910) |

**Table S15**: **Assessing the impact of data masking ratio of SQUALL pretraining (on *histMol*-low corpus).** Evaluation of data masking ratio impact. We randomly sampled 30% of *histMol*-low data tiles without replacement and employed a 100 epochs training schedule for this assessment. We opted to use complementary masking scheme to ensure training efficiency. Mask ratio here refers to the proportion of image regions that were masked during training. Pre-extracted tile features from each section were used to train a shallow ABMIL aggregation and classification head on curated tissue classification train-test folds (2,962: 321). Test performance was reported using top-1, top-3, and top-5 accuracy, weighted F1 score, and AUROC. The best and second best metrics are bolded and underlined, with the 95% confidence interval provided in parentheses.

| Stage | Par. | Val. | Top1-ACC | Top3-ACC | Top5-ACC | Weighted F1 | AUROC |
|---|---|---|---|---|---|---|---|
| Stage1 | lr | 5e-4 | .532 (.461-.595) | .751 (.685-.802) | .800 (.741-.849) | .488 (.419-.553) | .858 (.803-.898) |
| | | 2e-4 | **.683 (.615-.740)** | **.824 (.767-.869)** | **.893 (.841-.926)** | **.654 (.586-.714)** | **.918 (.869-.945)** |
| | | 1e-4 | .659 (.591-.718) | .824 (.767-.869) | .888 (.836-.922) | .618 (.547-.677) | .910 (.863-.941) |
| | | 5e-5 | .556 (.485-.618) | .746 (.680-.797) | .844 (.787-.886) | .511 (.442-.576) | .876 (.819-.910) |
| | bs | 64 | .649 (.581-.709) | **.829 (.761-.865)** | .878 (.825-.914) | .613 (.542-.673) | .913 (.863-.941) |
| | | 128 | **.683 (.615-.740)** | .824 (.767-.869) | **.893 (.841-.926)** | **.654 (.586-.714)** | **.918 (.869-.945)** |
| Stage2 | lr | 1e-4 | .432 (.363-.496) | .820 (.761-.865) | .896 (.847-.930) | .260 (.203-.320) | .506 (.433-.567) |
| | | 5e-5 | .470 (.400-.534) | .820 (.761-.865) | .924 (.880-.953) | .379 (.313-.443) | .568 (.499-.632) |
| | | 2e-5 | **.618 (.547-.677)** | **.874 (.819-.910)** | **.954 (.915-.974)** | **.568 (.499-.632)** | **.797 (.736-.844)** |
| | | 1e-5 | .432 (.363-.496) | .820 (.761-.865) | .895 (.847-.930) | .260 (.203-.320) | .495 (.428-.562) |
| | bs | 64 | .537 (.466-.600) | .840 (.782-.882) | .942 (.897-.964) | .459 (.391-.525) | .722 (.655-.775) |
| | | 128 | .618 (.547-.677) | .874 (.819-.910) | .954 (.915-.974) | .568 (.499-.632) | .797 (.736-.844) |
| | | 256 | **.636 (.566-.696)** | **.886 (.836-.922)** | **.956 (.915-.974)** | .575 (.504-.636) | **.813 (.751-.857)** |
| | | 512 | .630 (.561-.691) | .877 (.825-.914) | .955 (.915-.974) | **.577 (.509-.641)** | .806 (.746-.853) |
| | | 1,024 | .626 (.557-.687) | .876 (.819-.910) | .955 (.915-.974) | .566 (.494-.627) | .785 (.720-.832) |
| Full | lr | 1e-4 | .760 (.695-.810) | .943 (.903-.967) | .986 (.952-.993) | .741 (.675-.793) | .884 (.830-.918) |
| | | 5e-5 | **.778 (.715-.827)** | **.946 (.903-.967)** | .986 (.959-.995) | **.762 (.700-.815)** | **.890 (.836-.922)** |
| | | 2e-5 | .776 (.710-.823) | .946 (.903-.967) | **.987 (.959-.995)** | .760 (.695-.810) | .898 (.847-.930) |
| | | 1e-5 | .772 (.710-.823) | .946 (.903-.967) | .986 (.959-.995) | .755 (.690-.806) | .888 (.836-.922) |
| | | 5e-6 | .757 (.695-.810) | .941 (.897-.964) | .984 (.952-.993) | .738 (.670-.789) | .862 (.809-.902) |
| | bs | 256 | **.776 (.710-.823)** | .946 (.903-.967) | **.987 (.959-.995)** | **.760 (.695-.810)** | **.898 (.847-.930)** |
| | | 1,024 | .774 (.710-.823) | **.950 (.909-.971)** | .986 (.959-.995) | .758 (.695-.810) | .883 (.830-.918) |

**Table S16**: **Assessing the pretraining hyperparameters of SQUALL.** The hyperparameter grid search results obtained during pretraining are reported, providing a reference for future model development. We randomly sampled 30% of *histMol*-low data tiles without replacement and employed a 100-epoch training schedule. At each training stage, we first identified the optimal learning rate using a fixed batch size, and subsequently determined the optimal batch size while holding the learning rate constant. Full denotes continue Stage2 training with Stage 1 pretrained parameters. Experiments were performed under SQUALL-large settings. Pre-extracted tile features from each section were used to train a shallow ABMIL aggregation and classification head on curated tissue classification train-test folds (2,962: 321). Test performance was reported using top-1, top-3, and top-5 accuracy, weighted F1 score, and AUROC. The best and second best metrics are bolded and underlined for each parameter, with the 95% confidence interval provided in parentheses. Final optimal parameters are marked in grey. Leading zeros before decimal points were omitted to enhance readability. lr: learning rate, bs: batch size.

| SQUALL Arch. | Img. Arch. | #Tot. Para. | #Img. Para. | #Lyrs | #Fus. Lyrs | Tkn Dim. |
|---|---|---|---|---|---|---|
| SQUALL-base | ViT-base | 152M | 83M | 12 | 4 | 768 |
| SQUALL-large | ViT-large | 555M | 303M | 24 | 8 | 1,024 |

**Table S17**: **SQUALL architecture details.** We report the architectural details of the SQUALL encoders. In this study, we implement two model sizes, and the weights of all pretrained models of are freely available to the public for research purposes only.

| Arch. | Top1-ACC | Top3-ACC | Top5-ACC | Weighted F1 | AUROC |
|---|---|---|---|---|---|
| SQUALL-base | .639 (.571-.700) | .771 (.705-.819) | **.907 (.858-.938)** | .609 (.537-.668) | **.911 (.863-.941)** |
| SQUALL-large | **.663 (.596-.723)** | **.815 (.756-.861)** | .873 (.819-.910) | **.635 (.566-.696)** | .901 (.852-.934) |

**Table S18**: **Assessing the model scale of SQUALL (on *histMol*-low corpus).** We randomly sampled 30% of *histMol*-low data tiles without replacement and employed a 100 epochs training schedule for this assessment. Pre-extracted tile features from each section were used to train a shallow ABMIL aggregation and classification head on curated tissue classification train-test folds (2,962: 321). Test performance was reported using top-1, top-3, and top-5 accuracy, weighted F1 score, and AUROC. The best-performing metrics are highlighted in bold, with 95% confidence intervals provided in parentheses. Leading zeros before decimal points were omitted to enhance readability. Final model architectures were selected based on top-1 accuracy in the presented study.

| Epoch | Top1-ACC | Top3-ACC | Top5-ACC | Weighted F1 | AUROC |
|---|---|---|---|---|---|
| 100 | 0.683 (0.615-0.740) | 0.824 (0.767-0.869) | 0.893 (0.841-0.926) | 0.654 (0.586-0.714) | 0.918 (0.869-0.945) |
| 200 | 0.673 (0.605-0.731) | 0.839 (0.782-0.882) | 0.907 (0.858-0.938) | 0.657 (0.591-0.718) | 0.921 (0.874-0.949) |
| 400 | **0.751 (0.685-0.802)** | **0.893 (0.841-0.926)** | **0.937 (0.891-0.960)** | **0.727 (0.660-0.780)** | **0.926 (0.880-0.953)** |

**Table S19**: **Assessing the pretraining length of SQUALL (Stage 1).** Evaluation of pretraining length of SQUALL across training durations ranging from 100 to 400 epochs. SQUALL-large was pretrained on the entire *histMol*-low corpus. Pre-extracted tile features from each section were used to train a shallow ABMIL aggregation and classification head on curated tissue classification train-test folds (2,962: 321).. Test performance was reported using top-1, top-3, and top-5 accuracy, weighted F1 score, and AUROC. The best and second-best metrics are bolded and underlined, with the 95% confidence interval provided in parentheses.

| %Data | Top1-ACC | Top3-ACC | Top5-ACC | Weighted F1 | AUROC |
|---|---|---|---|---|---|
| 30% | 0.663 (0.596-0.723) | 0.815 (0.756-0.861) | 0.873 (0.819-0.910) | 0.635 (0.566-0.696) | 0.901 (0.852-0.934) |
| 100% | **0.683 (0.615-0.740)** | **0.824 (0.767-0.869)** | **0.893 (0.841-0.926)** | **0.654 (0.586-0.714)** | **0.918 (0.869-0.945)** |

**Table S20**: **Assessing the pretraining data scale of SQUALL (on *histMol*-low corpus).** Evaluation of pretraining data scale of SQUALL across different pretraining data fractions. SQUALL-large models were pretrained on different amount of the *histMol*-low corpus using a 100 epochs schedule. Pre-extracted tile features from each section were used to train a shallow ABMIL aggregation and classification head on curated tissue classification train-test folds (2,962: 321). Test performance was reported using top-1, top-3, and top-5 accuracy, weighted F1 score, and AUROC. The best metrics are bolded, with the 95% confidence provided in parentheses.

| Data comp. | Top1-ACC | Top3-ACC | Top5-ACC | Weighted F1 | AUROC |
|---|---|---|---|---|---|
| *histMol*-low | **.479 (.410-.544)** | **.813 (.751-.857)** | **.917 (.869-.945)** | **.468 (.400-.534)** | .697 (.630-.753) |
| w/o mus. | .458 (.391-.525) | .792 (.731-.840) | .917 (.869-.945) | .421 (.354-.487) | .684 (.615-.740) |
| Visium only | .438 (.368-.501) | .771 (.705-.819) | .917 (.869-.945) | .405 (.336-.468) | **.698 (.630-.753)** |

**Table S21**: **Assessing the pretraining data composition of SQUALL (on *histMol*-low corpus).** Evaluation of pretraining data composition during SQUALL pretraining. We randomly sampled a 30% subset of the total training tiles used for SQUALL-large pretraining. For 'w/o mus.', we removed all mouse tiles from this subset. For 'Visium only', we removed all data tiles obtained from other platforms. For all experiments, we ensured that the data passed through the model with an equivalent number of iterations (about 60,600 times). Pre-extracted tile features from each section were used to train a shallow ABMIL aggregation and classification head on curated human cancer classification train-test folds (764: 76). Test performance was reported using top-1, top-3, and top-5 accuracy, weighted F1 score, and AUROC. The best and second-best metrics are bolded and underlined, with the 95% confidence interval provided in parentheses. Leading zeros before decimal points were omitted to enhance readability.

| Major class | Minor class | #Poly. | #Tile (Train: Test) | Occur. |
|---|---|---|---|---|
| Normal parenchyma | Ovary parenchyma | 13 | 10,515 (7,360: 3,155) | OC |
| | Myometrium | 3 | 2,993 (2,095: 898) | CC/EC |
| | Normal | 3 | 2,988 (2,091: 897) | CC/OC |
| | Muscular layer | 1 | 391 (273: 118) | OC |
| | Corpus albicans | 15 | 53 (37: 16) | OC |
| | Mucosal glands | 51 | 37 (25: 12) | CC/EC |
| | Endometrium | 23 | 16 (11: 5) | EC/OC |
| | Fallopian tube | 3 | - | OC |
| | Glandular epithelium | 2 | - | OC |
| | Squamous epithelium | 1 | - | CC |
| Tumor parenchyma | Tumor | 2,983 | 42,087 (29,460: 12,627) | CC/EC/OC |
| | Sarcoma component | 10 | 3,080 (2,156: 924) | EC |
| | Carcinoma component | 50 | 622 (435: 187) | EC |
| Interstitial | Adipose tissue | 31 | 1,861 (1,302: 559) | CC/OC |
| | Immune cells | 209 | 777 (543: 234) | CC/EC/OC |
| | Blood vessel | 28 | - | CC |
| Stroma | Simple stroma | 926 | 27,367 (19,156: 8,211) | CC/EC/OC |
| | Complex stroma | 95 | 351 (245: 106) | CC/EC/OC |
| Others | Unknown | 96 | - | CC/EC/OC |
| | Other (Ambiguous) | 48 | - | CC/OC |
| | Background | 2 | - | OC |
| Pathological regions | Debris | 474 | 4,277 (2,993: 1,284) | CC/EC/OC |
| | Necrosis | 25 | 15 (10: 5) | CC/OC |
| | Calcification | 13 | 12 (8: 4) | OC |
| | Sq. metaplasia (glandular epi.) | 12 | 5 (3: 2) | CC |
| | Ectopic endometrium | 8 | - | OC |
| | Benign cyst | 6 | - | OC |
| | L-Grade IN (epi. neopl.) | 3 | - | OC |
| | Paramesonephric duct cyst | 3 | - | OC |
| Total | | 5,137 | 97,447 (68,203: 29,244) | |

**Table S22**: **Annotation hierarchy and summary for gynecological cancer subset of *histMol*-high.** Three experienced pathologists were involved in the tissue annotation process. Categories were merged based on their consensus. All annotations were formatted as polygon or multi-polygon objects in GeoJSON. Each tile (224 × 224 pixel image region) was assigned a label if at least 80% of its area intersected with a polygon. Tiles containing multiple annotations were discarded. The tile labels were subsequently used in a classification task to evaluate the model behavior after different stages of pretraining. For this task, a 70: 30 (68,203: 29,244) train-test split is applied. Labels with less than 5 tiles are excluded from the test set. This annotation set was used to assess *histMol*-high pretraining and provides labels for ovarian cancer analysis.

| Config. | Value |
|---|---|
| Num. of class | 18 |
| Layers | 1 |
| Layers type | Linear |
| Optimization target | Cross Entropy |
| Optimizer | AdamW |
| Optimizer momentum | $\beta_1 = 0.9$, $\beta_2 = 0.999$ |
| Peak learning rate | 1e-4 |
| Learning rate schedule | Cosine Annealing |
| Epochs | 50 |
| Batch size | 128 |

**Table S23**: **Configuration for SQUALL tile classification probing task (on *histMol*-high corpus).** An NVIDIA H100 GPU was used for classification evaluation. To assess the quality of the extracted embeddings, we employed a simple linear classifier on the annotated tiles.

| Stage | Ep. | Top1-ACC | Top3-ACC | Top5-ACC | Weighted F1 | AUROC |
|---|---|---|---|---|---|---|
| Stage1 | 400 | .706 (.640-.762) | .930 (.885-.956) | .979 (.946-.990) | .672 (.605-.731) | .831 (.772-.874) |
| Stage2 | 400 | .759 (.695-.810) | .941 (.897-.964) | .983 (.952-.993) | .741 (.675-.793) | **.904 (.852-.934)** |
| Full | 100 | **.778 (.715-.827)** | **.946 (.903-.967)** | **.987 (.959-.995)** | **.762 (.700-.815)** | .890 (.836-.922) |

**Table S24**: **Assessing the pretraining stage of SQUALL.** SQUALL was initially trained on the *histMol*-low corpus (Stage 1), followed by training on the *histMol*-high corpus (Stage 2). We also included a baseline model that was pretrained only on the *histMol*-high corpus with random parameter initialization. The results indicated that after Stage 1 training, a 100-epochs training schedule on the *histMol*-high corpus already outperformed Stage 2 training on tile classification, even with a longer training schedule (100 vs. 400 epochs). Pre-extracted tile features from each section were used to train a shallow MLP classification head on curated tissue annotation classification train-test folds (68,203: 29,244) of gynecological cancer annotated tiles. Test performance was evaluated using top-1, top-3, and top-5 accuracy, weighted F1 score, and AUROC. The best and second-best metrics are highlighted in bold and underlined, with the 95% confidence interval provided in parentheses. Leading zeros before decimal points were omitted to enhance readability.

| Epoch | Top1-ACC | Top3-ACC | Top5-ACC | Weighted F1 | AUROC |
|---|---|---|---|---|---|
| 100 | 0.710 (0.645-0.767) | 0.925 (0.880-0.953) | 0.976 (0.939-0.987) | 0.689 (0.620-0.745) | 0.850 (0.793-0.890) |
| 200 | 0.755 (0.690-0.806) | 0.940 (0.897-0.964) | 0.982 (0.952-0.993) | 0.738 (0.670-0.789) | **0.912 (0.863-0.941)** |
| 400 | **0.759 (0.695-0.810)** | **0.941 (0.897-0.964)** | **0.983 (0.952-0.993)** | **0.741 (0.675-0.793)** | 0.904 (0.852-0.934) |

**Table S25**: **Assessing the pretraining length of SQUALL (Stage 2).** Evaluation of pretraining length of SQUALL across different training durations ranging from 100 to 400 epochs. SQUALL-large was pretrained on the entire *histMol*-high corpus with random parameter initialization. Pre-extracted tile features from each section were used to train a shallow MLP classification head on curated tissue annotation classification train-test folds (68,203: 29,244) of gynecological cancer annotated tiles. Test performance was reported using top-1, top-3, and top-5 accuracy, weighted F1 score, and AUROC. The best and second-best metrics are bolded and underlined, with the 95% confidence interval provided in parentheses.

**Panel A. SQUALL design-choice ablations across downstream tasks**

| Data | Cat | Setting | Virtual expression profiling (All sections) | | | Spatially aware clustering | | Clinical-related prediction | | Top-3 metric proportion |
|---|---|---|---|---|---|---|---|---|---|---|
| | | | PCC | AUC | SSIM | ARI | AMI | C-index (%) | F1 (%) | |
| 30% | alignment | W/o ST | – | – | – | 0.30 (-0.02–0.62) | 0.28 (0.03–0.53) | 54.50 (36.56–72.44) | 65.56 (62.42–68.69) | – |
| | alignment | Direct-prediction | 0.02 (-0.04–0.08) | 0.50 (0.50–0.51) | 0.11 (-0.10–0.33) | **0.30 (-0.03–0.62)** | 0.26 (0.00–0.51) | 55.45 (46.95–63.95) | 66.79 (62.42–71.17) | 2/7 (28.6%) |
| | data | W/o mouse | 0.03 (-0.15–0.21) | 0.51 (0.44–0.57) | 0.13 (-0.03–0.29) | 0.24 (-0.01–0.49) | 0.24 (0.03–0.45) | 65.53 (52.68–78.38) | 66.79 (62.42–71.17) | 2/7 (28.6%) |
| | pre-norm | Rank | 0.13 (-0.19–0.45) | 0.55 (0.43–0.66) | 0.22 (-0.01–0.45) | 0.20 (-0.03–0.43) | 0.23 (0.01–0.46) | 65.53 (52.68–78.38) | 66.79 (62.42–71.17) | 2/7 (28.6%) |
| | pre-norm | Log1p | *0.24 (-0.21–0.69)* | *0.60 (0.44–0.76)* | 0.24 (0.11–0.37) | 0.30 (0.06–0.53) | **0.31 (0.08–0.53)** | 54.89 (40.72–69.06) | 66.20 (61.39–71.00) | 4/7 (57.1%) |
| | pre-image | UNI init | 0.21 (-0.27–0.70) | 0.59 (0.41–0.77) | *0.23 (0.10–0.36)* | 0.11 (0.07–0.15) | 0.14 (0.11–0.17) | 56.76 (37.01–76.51) | 65.52 (61.64–69.40) | 1/7 (14.3%) |
| | pre-interp | Cubic | 0.06 (-0.35–0.47) | 0.51 (0.38–0.64) | 0.16 (0.00–0.32) | 0.25 (-0.03–0.53) | 0.23 (0.00–0.47) | **66.27 (60.09–72.45)** | 66.79 (62.42–71.17) | 2/7 (28.6%) |
| | pre-interp | Linear | **0.28 (-0.16–0.73)** | **0.63 (0.46–0.80)** | 0.27 (0.19–0.36) | 0.27 (0.03–0.50) | 0.29 (0.07–0.51) | 54.83 (47.44–62.22) | 66.20 (61.39–71.00) | 4/7 (57.1%) |
| | SQUALL | SQUALL | 0.27 (-0.18–0.72) | 0.62 (0.46–0.78) | **0.28 (0.18–0.37)** | *0.29 (-0.02–0.59)* | *0.26 (0.03–0.49)* | 57.20 (47.54–66.87) | **66.79 (62.42–71.17)** | **6/7 (85.7%)** |
| 100% | SQUALL | SQUALL | 0.33 (-0.07–0.74) | 0.58 (0.46–0.70) | 0.23 (0.08–0.38) | 0.26 (0.09–0.44) | 0.33 (0.16–0.49) | 62.10 (51.73–72.49) | 69.00 (58.34–79.61) | – |

**Panel B. Matched image-only versus multimodal pretraining**

| Setting | Virtual expression profiling (Xenium 5K) | | |
|---|---|---|---|
| | PCC | AUC | SSIM |
| Image only | 0.13 (0.12–0.14) | 0.52 (0.51–0.52) | 0.20 (0.20–0.21) |
| SQUALL | **0.48 (0.23–0.74)** | **0.70 (0.59–0.80)** | **0.23 (0.08–0.38)** |

**Table S26**: **Analysis of SQUALL design choices and matched comparison of image-only and multimodal pretraining. Panel A.** Alternative SQUALL design choices were evaluated using a randomly sampled 30% subset of the tiles used for pretraining. Downstream performance was assessed using the Pearson correlation coefficient (PCC), area under the receiver operating characteristic curve (AUC), and structural similarity index measure (SSIM) for virtual spatial gene expression profiling; adjusted Rand index (ARI) and adjusted mutual information (AMI) for representative spatially aware clustering tasks; concordance index (C-index) for survival prediction; and F1 score for drug-response prediction. Both clinical tasks were evaluated on the CGODC-OV cohort. Values are reported as the mean, with the lower and upper bounds of the 95% confidence interval shown in parentheses. C-index and F1 are reported as percentages. The best, second-best, and third-best results are shown in bold, underlined, and italics, respectively. The Top-3 metric proportion indicates the number and percentage of the seven metrics for which each setting ranked among the top three. The W/o ST setting and the complete SQUALL model were included as reference models and excluded from the Top-3 ranking. W/o ST denotes an MAE trained exclusively on histology images without spatial transcriptomic data. Direct-prediction denotes direct prediction of gene expression from histology images, corresponding to masking 100% of the expression input in SQUALL. W/o mouse denotes removal of all mouse data from the pretraining dataset. UNI init denotes the use of UNI-derived histology features during multimodal pretraining. Pre-norm compares alternative normalization methods for preprocessing spatial transcriptomic data, whereas pre-interp compares interpolation methods used for pseudo-label construction. For each ablation experiment, only the indicated component was changed, while all other components were kept identical to the corresponding SQUALL configuration. **Panel B.** The image-only model and multimodal SQUALL model were trained on the same randomly sampled 30% subset of pretraining tiles, matching histology exposure while differing in whether spatial transcriptomic supervision was used. Because the image-only model does not contain a molecular decoder, its predictions were generated using the same prediction-head framework applied to pathology foundation-model representations. Performance was evaluated on Xenium 5K sections using PCC, AUC, and SSIM. Values are reported as means, with 95% confidence intervals in parentheses.

| Config. | base | large |
|---|---|---|
| Encoder layers | 12 | 24 |
| Decoder layers | 4 | 4 |
| Fusion layers | 4 | 8 |
| Heads | 12 | 16 |
| Drop path rate | 0.1 | |
| Position encoding | Relative + Absolute | |
| FFN layer | MLP | |
| Head activation | GELU | |
| Embedding Dim. | 768 | 1,024 |
| MLP Dim. | 768 | 1,024 |
| Normalize last layer | ✓ | |
| Image input size | 224 × 224 × 3 | |
| Expression input size | 56 × 56 × 15,757 | |
| Mask ratio | 0.5 | |
| Token size per tile | 14×14 | |
| Total epochs | 100 | 400 |
| Warm-up epochs | 5 | |
| Batch size | 128 | |
| Optimizer | AdamW | |
| Optimizer momentum | $\beta_1 = 0.9$, $\beta_2 = 0.999$ | |
| Learning rate | 2e-4 | |
| Weight decay | 0.05 | |
| Weight decay scheduler | Cosine | |
| Automatic mixed precision | ✓ | |

**Table S27**: **Configuration for SQUALL pretraining (Stage 1, on *histMol*-low corpus).** Four nodes, each with 8 × NVIDIA H100 GPUs linked with **InfiniBand** were used for pretraining. The batch size refers to the total batch size across GPUs, with each node utilizing **NVLink** for intra-node GPU communication. 'Token size per tile' refers to the spatial size of the SQUALL feature map after the projection layer. We opted to keep it fixed across both the base and large model variants to ensure resolution consistency, which resulted in a slight decrease in the number of parameters at the large setting.

| Config. | From scratch | Stage 1 init. |
|---|---|---|
| Encoder layers | 24 | |
| Decoder layers | 4 | |
| Fusion layers | 8 | |
| Heads | 16 | |
| Drop path rate | 0.1 | |
| Position encoding type | Relative + Absolute | |
| FFN layer | MLP | |
| Head activation | GELU | |
| Embedding Dim. | 1,024 | |
| MLP Dim. | 1,024 | |
| Normalize last layer | ✓ | |
| Image input size | 224 × 224 × 3 | |
| Expression input size | 56 × 56 × 15,757 | |
| Mask ratio | 0.5 | |
| Token size per tile | 14× 14 | |
| Embedding Dim. | 1,024 | |
| Total epochs | 400 | 100 |
| Warm up epochs | 5 | |
| Batch size | 256 | |
| Optimizer | AdamW | |
| Optimizer momentum | $\beta_1 = 0.9$, $\beta_2 = 0.999$ | |
| Learning rate | 2e-5 | |
| Weight decay | 0.05 | |
| Weight decay scheduler | Cosine | |
| Automatic mixed precision | ✓ | |

**Table S28**: **Configuration for SQUALL pretraining (Stage 2, on *histMol*-high corpus).** 4 nodes, each with 8 × NVIDIA H100 GPUs linked with **InfiniBand** were used for pretraining. The batch size refers to the total batch size across GPUs, with each node utilizing **NVLink** for intra-node GPU communication.

| Setting | Dataset | Platform | Tissue | Preservation | Metadata |
|---|---|---|---|---|---|
| Training | SPATCH-HCC | Visium HD | Liver | FFPE | Hepatocellular carcinoma: II(T2N0M0) |
| | SPATCH-OV | Visium HD | Ovary | FFPE | High-grade serous ovarian carcinoma: III-C (T3cN0M0) |
| Evaluation | SPATCH-HCC | Xenium 5K | Liver | FFPE | Hepatocellular carcinoma: II(T2N0M0), adjacency section |
| | 10×-OV | Xenium 5K | Ovary | FFPE | Ovarian papillary serous carcinoma: III-B(T3bN0MX) |

**Table S29**: **Data characteristics of the virtual spatial gene expression profiling dataset for comparative evaluation.** To provide paired training data for the compared models, we collected Visium HD and Xenium 5K sections following the cross-platform correspondence established in previous spatial transcriptomics benchmark study. All datasets summarized here were not included in SQUALL pretraining at either the tissue-block or patient level, and SQUALL parameters were kept frozen during inference. The corresponding compared models were trained using Visium HD section and evaluated on Xenium 5K sections within the same tumor category.

| Cat. | Model | Encoder Arch. | PT data | Pred. head Arch. | #Para. | Pred. Res. |
|---|---|---|---|---|---|---|
| TSM | ST-Net | DenseNet-121 | ImageNet | FCN (1×, rand. init.) | 7.98M | 112 $\mu$m |
| TSM | iStar | ViT-S/256 | HIPT (TCGA) | MLP (4×, rand. init.) | 9.6M | 2 $\mu$m |
| TSM | DeepPT | ResNet-50 | ImageNet | AE + MLP (3×) | 30.4M | 112 $\mu$m |
| TSM | Hist2ST | Hybridize | - | Linear + ZINB heads | 92.4M | 112 $\mu$m |
| TSM | EGN | ResNet-50 | ImageNet | Exemplar regression head | 149.5M | 112 $\mu$m |
| TSM | Path2Space | CTransPath | TCGA | MLP (2×, rand. init.) | 31.8M | 112 $\mu$m |
| PFM | UNI | ViT-L | BWH, MGH, GTEx | MLP (4×, rand. init.) | 307M | 112 $\mu$m |
| PFM | PLIP | ViT-B | OpenPath (WIT400M) | MLP (4×, rand. init.) | 86M | 112 $\mu$m |
| PFM | Virchow | ViT-H | MSKCC | MLP (4×, rand. init.) | 632M | 112 $\mu$m |
| MM | OmiCLIP | ViT | ST-bank & LAION-5B | LokiPredEX | 306.7M | 112 $\mu$m |
| MM | Multi-Embed | UNI + Alignment | TCGA | MLP (4×, rand. init.) | 338.2M | 112 $\mu$m |
| MM | SQUALL | ViT-L/224 | *histMol* | Trans. (4×, SQUALL decoder) | 354M | 2 $\mu$m |

**Table S30**: **Architectural details of models included in the comparative evaluation of virtual spatial gene expression profiling.** We summarize the architectural configurations of all evaluated models based on their original publications or publicly available implementations. All models were evaluated on Xenium 5K sections following their respective evaluation protocols, while models requiring task-specific optimization were trained on Visium HD sections. Specifically, task-specific models were trained on Visium HD sections and evaluated on either spatially adjacent Xenium 5K sections or independent Xenium 5K sections from the same tumor type. For pathology foundation model benchmarking, cancer-specific models were trained using a 9:1 training-validation split of the Visium HD sections, with model selection based on validation performance. The selected models were subsequently applied to the corresponding Xenium 5K prediction tasks. All training, model selection, and evaluation procedures were performed within the same tumor type. None of the evaluation datasets were included in SQUALL pretraining at either the tissue-block or patient level. SQUALL predictions were generated directly using its pretrained decoder, with all parameters kept frozen during inference. OmiCLIP and Multi-Embed were additionally included as multimodal embedding baselines. OmiCLIP predictions were generated directly from its pretrained image-spatial transcriptomics similarity matrix, whereas Multi-Embed predictions were generated following its original training protocol. For a unified comparison, all predictions were evaluated at a common spatial resolution using 112 $\mu$m × 112 $\mu$m image tiles. All training and inference procedures performed for this comparsion used a single NVIDIA L40 GPU or A100 GPU. PT denotes the pretraining data used for representation learning, while TSM, PFM, and MM denote task-specific models, pathology foundation models, and multimodal models, respectively.

| Config. | Virtual expression prediction |
|---|---|
| Backbone | UNI/Virchow/PLIP |
| Maximum epochs | 50 |
| Early-stopping patience | 10 epochs |
| Minimum improvement | 0.0 |
| Batch size | 32 |
| Dropout | 0.2 |
| Learning rate | $1 \times 10^{-4}$ |

**Table S31**: **Configuration for virtual expression prediction using pathology foundation models.** Gene expression targets were defined using either the *histMol* gene set or the intersection gene set. Models were trained for up to 50 epochs, with early stopping applied when validation performance did not improve for 10 consecutive epochs. All experiments were conducted using NVIDIA A100 GPUs.

| Dataset | Model | mPCC | mAUC | mSSIM |
|---|---|---|---|---|
| HCC | ST-Net | 0.333 (0.168–0.499) | 0.516 (0.498–0.535) | 0.150 (0.131–0.169) |
| | iStar | 0.383 (0.191–0.574) | 0.528 (0.498–0.557) | 0.181 (0.128–0.234) |
| | DeepPT | <u>0.397 (0.182–0.613)</u> | 0.532 (0.480–0.584) | 0.204 (0.090–0.319) |
| | Hist2ST | 0.335 (0.159–0.512) | 0.500 (0.461–0.540) | 0.140 (0.065–0.215) |
| | EGN | 0.305 (0.152–0.459) | 0.479 (0.428–0.530) | 0.117 (0.038–0.196) |
| | Path2Space | 0.362 (0.117–0.608) | <u>0.539 (0.490–0.587)</u> | 0.244 (0.129–0.360) |
| | UNI | 0.238 (-0.012–0.489) | 0.536 (0.486–0.585) | <u>0.267 (0.076–0.458)</u> |
| | PLIP | 0.124 (-0.068–0.315) | 0.504 (0.471–0.536) | 0.205 (0.022–0.389) |
| | Virchow | 0.267 (0.025–0.510) | 0.533 (0.494–0.571) | **0.283 (0.107–0.460)** |
| | OmiCLIP | 0.367 (0.181–0.552) | 0.511 (0.477–0.545) | 0.167 (0.099–0.234) |
| | Multi-Embed | 0.306 (0.152–0.459) | 0.481 (0.437–0.526) | 0.092 (0.010–0.175) |
| | SQUALL | **0.421 (0.199–0.644)** | **0.545 (0.500–0.590)** | 0.211 (0.132–0.289) |
| OC | ST-Net | 0.237 (0.098–0.376) | 0.513 (0.453–0.573) | 0.083 (0.064–0.102) |
| | iStar | <u>0.481 (0.308–0.654)</u> | <u>0.660 (0.602–0.718)</u> | 0.108 (0.070–0.146) |
| | DeepPT | 0.397 (0.133–0.661) | 0.590 (0.460–0.719) | 0.160 (0.048–0.272) |
| | Hist2ST | 0.236 (0.042–0.429) | 0.501 (0.397–0.606) | 0.077 (0.009–0.145) |
| | EGN | 0.438 (0.144–0.732) | 0.624 (0.470–0.778) | <u>0.184 (0.058–0.309)</u> |
| | Path2Space | 0.390 (0.106–0.675) | 0.608 (0.511–0.706) | **0.208 (0.111–0.304)** |
| | UNI | 0.238 (-0.015–0.491) | 0.565 (0.478–0.652) | 0.160 (0.071–0.249) |
| | PLIP | 0.258 (-0.018–0.534) | 0.560 (0.443–0.678) | 0.161 (0.075–0.248) |
| | Virchow | 0.274 (-0.007–0.555) | 0.581 (0.482–0.680) | 0.175 (0.077–0.272) |
| | OmiCLIP | 0.353 (0.089–0.617) | 0.562 (0.426–0.698) | 0.144 (0.038–0.251) |
| | Multi-Embed | 0.277 (0.081–0.474) | 0.530 (0.430–0.630) | 0.085 (0.016–0.153) |
| | SQUALL | **0.569 (0.383–0.754)** | **0.719 (0.660–0.778)** | 0.156 (0.089–0.223) |
| Overall | ST-Net | 0.285 (0.125–0.446) | 0.515 (0.470–0.559) | 0.117 (0.078–0.155) |
| | iStar | <u>0.432 (0.243–0.621)</u> | <u>0.594 (0.513–0.675)</u> | 0.144 (0.086–0.203) |
| | DeepPT | 0.397 (0.156–0.639) | 0.561 (0.458–0.664) | 0.182 (0.067–0.297) |
| | Hist2ST | 0.286 (0.094–0.478) | 0.501 (0.422–0.580) | 0.109 (0.030–0.187) |
| | EGN | 0.372 (0.128–0.616) | 0.552 (0.416–0.687) | 0.150 (0.040–0.260) |
| | Path2Space | 0.376 (0.110–0.643) | 0.574 (0.489–0.658) | <u>0.226 (0.118–0.334)</u> |
| | UNI | 0.238 (-0.014–0.490) | 0.550 (0.478–0.622) | 0.214 (0.055–0.372) |
| | PLIP | 0.191 (-0.056–0.438) | 0.532 (0.441–0.623) | 0.183 (0.038–0.328) |
| | Virchow | 0.271 (0.008–0.534) | 0.557 (0.478–0.636) | **0.229 (0.077–0.381)** |
| | OmiCLIP | 0.360 (0.132–0.588) | 0.537 (0.434–0.639) | 0.155 (0.065–0.246) |
| | Multi-Embed | 0.291 (0.115–0.468) | 0.506 (0.424–0.587) | 0.089 (0.013–0.165) |
| | SQUALL | **0.495 (0.277–0.713)** | **0.632 (0.530–0.734)** | 0.183 (0.106–0.261) |

**Table S32**: **Virtual spatial gene expression profiling performance of comparative evaluation.** Predictions generated at each model's native spatial resolution were subsequently aggregated into 224 × 224-pixel tiles to ensure evaluation at a common spatial resolution. The Pearson correlation coefficient (PCC), area under the curve (AUC) and structural similarity index measure (SSIM) were computed independently for each of the 4,421 overlapping genes on Xenium 5K sections, and the resulting gene-wise scores were averaged across genes to obtain mPCC, mAUC, and mSSIM, respectively. The best and second-best values for each metric are shown in bold and underlined, respectively. Values in parentheses indicate the lower and upper bounds corresponding to one standard deviation from the mean.

| Gene set | Sources | #Genes | Note |
|---|---|---|---|
| Transcription factors | PMID: 33257861; 36608654 | 599 | Provides insight on regional regulations. |
| Ligands and receptors | PMID: 32103204 | 639 | Provides insight on cellular communication and signaling. |
| Cytokine and chemokine | PMID: 30184180 | 74 | Provides insight on immune response. |
| Membrane proteins | PMID: 35121907 | 1,146 | Rich source of immune and targeted therapies. |

**Table S33**: **Gene sets used for virtual spatial biomarker profiling in the comparative evaluation.** We report the gene lists defining each biomarker category. To evaluate virtual spatial expression profiling for clinically and biologically relevant biomarkers.

| Set | Model | HCC | | | OV | | | Overall | | |
|---|---|---|---|---|---|---|---|---|---|---|
| | | mPCC | mAUC | mSSIM | mPCC | mAUC | mSSIM | mPCC | mAUC | mSSIM |
| TFs | ST-Net | 0.33 | 0.51 | 0.15 | 0.23 | 0.51 | 0.08 | 0.28 (0.17–0.39) | 0.51 (0.48–0.54) | 0.12 (0.10–0.13) |
| | iStar | 0.38 | 0.53 | 0.18 | 0.49 | 0.66 | 0.11 | 0.43 (0.30–0.56) | 0.59 (0.56–0.63) | 0.14 (0.11–0.17) |
| | DeepPT | 0.38 | 0.52 | 0.18 | 0.42 | 0.60 | 0.17 | 0.40 (0.23–0.56) | 0.56 (0.49–0.63) | 0.18 (0.10–0.25) |
| | Hist2ST | 0.33 | 0.50 | 0.14 | 0.24 | 0.50 | 0.08 | 0.29 (0.15–0.42) | 0.50 (0.44–0.56) | 0.11 (0.06–0.16) |
| | EGN | 0.31 | 0.48 | 0.12 | 0.47 | 0.64 | 0.20 | 0.39 (0.22–0.55) | 0.56 (0.49–0.64) | 0.16 (0.08–0.23) |
| | Path2Space | 0.34 | 0.53 | 0.22 | 0.41 | 0.62 | **0.22** | 0.37 (0.19–0.56) | 0.57 (0.52–0.62) | 0.22 (0.15–0.29) |
| | UNI | 0.20 | 0.53 | 0.26 | 0.25 | 0.57 | 0.17 | 0.23 (0.06–0.40) | 0.55 (0.50–0.60) | 0.21 (0.10–0.33) |
| | PLIP | 0.11 | 0.50 | 0.22 | 0.28 | 0.57 | 0.16 | 0.20 (0.03–0.36) | 0.54 (0.47–0.60) | 0.19 (0.08–0.30) |
| | Virchow | 0.24 | 0.53 | **0.28** | 0.28 | 0.58 | 0.18 | 0.26 (0.08–0.45) | 0.56 (0.50–0.61) | **0.23 (0.12–0.34)** |
| | OmiCLIP | 0.37 | 0.52 | 0.17 | 0.36 | 0.56 | 0.15 | 0.37 (0.20–0.53) | 0.54 (0.47–0.61) | 0.16 (0.10–0.23) |
| | Multi-Embed | 0.31 | 0.49 | 0.10 | 0.29 | 0.53 | 0.09 | 0.30 (0.17–0.42) | 0.51 (0.46–0.56) | 0.10 (0.05–0.14) |
| | SQUALL | **0.41** | **0.54** | 0.20 | **0.58** | **0.72** | 0.16 | **0.49 (0.35–0.64)** | **0.63 (0.59–0.67)** | 0.18 (0.13–0.23) |
| LRs | ST-Net | 0.26 | 0.51 | 0.14 | 0.19 | 0.51 | 0.08 | 0.22 (0.12–0.33) | 0.51 (0.48–0.54) | 0.11 (0.10–0.12) |
| | iStar | 0.29 | 0.52 | 0.16 | 0.38 | 0.63 | 0.09 | 0.34 (0.21–0.47) | 0.58 (0.55–0.61) | 0.12 (0.09–0.15) |
| | DeepPT | 0.30 | 0.52 | 0.17 | 0.25 | 0.53 | 0.11 | 0.28 (0.12–0.44) | 0.53 (0.46–0.59) | 0.14 (0.07–0.21) |
| | Hist2ST | 0.26 | 0.50 | 0.12 | 0.19 | 0.50 | 0.06 | 0.23 (0.10–0.35) | 0.50 (0.45–0.55) | 0.09 (0.05–0.14) |
| | EGN | 0.25 | 0.49 | 0.12 | 0.28 | 0.56 | 0.12 | 0.27 (0.10–0.43) | 0.53 (0.44–0.61) | 0.12 (0.05–0.19) |
| | Path2Space | 0.26 | 0.53 | 0.22 | 0.23 | 0.55 | 0.17 | 0.24 (0.06–0.42) | 0.54 (0.49–0.59) | 0.19 (0.12–0.27) |
| | UNI | 0.15 | 0.52 | 0.29 | 0.12 | 0.53 | 0.17 | 0.14 (-0.02–0.29) | 0.53 (0.49–0.57) | 0.23 (0.11–0.35) |
| | PLIP | 0.09 | 0.50 | 0.25 | 0.12 | 0.52 | 0.16 | 0.10 (-0.03–0.24) | 0.51 (0.46–0.56) | 0.20 (0.08–0.32) |
| | Virchow | 0.16 | 0.52 | **0.29** | 0.15 | 0.54 | **0.17** | 0.16 (-0.01–0.32) | 0.53 (0.49–0.58) | **0.23 (0.12–0.35)** |
| | OmiCLIP | 0.28 | 0.51 | 0.14 | 0.23 | 0.51 | 0.09 | 0.25 (0.11–0.40) | 0.51 (0.45–0.58) | 0.12 (0.06–0.17) |
| | Multi-Embed | 0.25 | 0.49 | 0.10 | 0.22 | 0.52 | 0.07 | 0.23 (0.11–0.36) | 0.51 (0.45–0.56) | 0.09 (0.04–0.14) |
| | SQUALL | **0.32** | **0.53** | 0.17 | **0.47** | **0.69** | 0.12 | **0.39 (0.25–0.54)** | **0.61 (0.57–0.65)** | 0.15 (0.10–0.20) |
| CCs | ST-Net | 0.19 | 0.51 | 0.14 | 0.15 | 0.50 | 0.07 | 0.17 (0.08–0.27) | 0.50 (0.48–0.53) | 0.10 (0.09–0.11) |
| | iStar | 0.23 | 0.52 | 0.14 | 0.33 | 0.63 | 0.08 | 0.28 (0.16–0.39) | 0.58 (0.54–0.61) | 0.11 (0.08–0.13) |
| | DeepPT | 0.22 | 0.52 | 0.13 | 0.20 | 0.52 | 0.08 | 0.21 (0.07–0.35) | 0.52 (0.46–0.58) | 0.11 (0.05–0.17) |
| | Hist2ST | 0.19 | 0.50 | 0.10 | 0.13 | 0.48 | 0.05 | 0.16 (0.05–0.27) | 0.49 (0.44–0.54) | 0.07 (0.04–0.11) |
| | EGN | 0.19 | 0.49 | 0.11 | 0.22 | 0.55 | 0.09 | 0.21 (0.07–0.35) | 0.52 (0.44–0.60) | 0.10 (0.05–0.14) |
| | Path2Space | 0.17 | 0.52 | 0.19 | 0.14 | 0.53 | 0.15 | 0.16 (0.01–0.30) | 0.52 (0.48–0.57) | 0.17 (0.11–0.23) |
| | UNI | 0.06 | 0.51 | **0.31** | 0.06 | 0.51 | 0.17 | 0.06 (-0.04–0.16) | 0.51 (0.49–0.53) | 0.24 (0.11–0.37) |
| | PLIP | 0.05 | 0.50 | 0.29 | 0.09 | 0.52 | 0.17 | 0.07 (-0.05–0.19) | 0.51 (0.46–0.56) | 0.23 (0.10–0.36) |
| | Virchow | 0.08 | 0.51 | 0.30 | 0.09 | 0.52 | **0.17** | 0.08 (-0.04–0.20) | 0.52 (0.49–0.55) | **0.24 (0.12–0.36)** |
| | OmiCLIP | 0.22 | 0.52 | 0.14 | 0.17 | 0.50 | 0.07 | 0.20 (0.07–0.32) | 0.51 (0.45–0.56) | 0.10 (0.06–0.14) |
| | Multi-Embed | 0.20 | 0.50 | 0.11 | 0.17 | 0.51 | 0.06 | 0.19 (0.07–0.31) | 0.50 (0.45–0.56) | 0.08 (0.04–0.13) |
| | SQUALL | **0.23** | **0.52** | 0.15 | **0.39** | **0.67** | 0.10 | **0.31 (0.18–0.44)** | **0.60 (0.56–0.63)** | 0.12 (0.09–0.16) |
| MPs | ST-Net | 0.28 | 0.52 | 0.14 | 0.21 | 0.51 | 0.08 | 0.24 (0.13–0.35) | 0.51 (0.48–0.54) | 0.11 (0.10–0.12) |
| | iStar | 0.32 | 0.52 | 0.16 | 0.40 | 0.64 | 0.09 | 0.36 (0.23–0.49) | 0.58 (0.55–0.61) | 0.13 (0.10–0.16) |
| | DeepPT | 0.33 | 0.53 | 0.18 | 0.29 | 0.54 | 0.12 | 0.31 (0.15–0.47) | 0.54 (0.47–0.60) | 0.15 (0.08–0.22) |
| | Hist2ST | 0.28 | 0.50 | 0.13 | 0.21 | 0.50 | 0.07 | 0.24 (0.11–0.37) | 0.50 (0.45–0.55) | 0.10 (0.05–0.14) |
| | EGN | 0.27 | 0.49 | 0.12 | 0.30 | 0.57 | 0.13 | 0.29 (0.12–0.45) | 0.53 (0.44–0.62) | 0.12 (0.06–0.19) |
| | Path2Space | 0.28 | 0.53 | 0.22 | 0.28 | 0.57 | **0.18** | 0.28 (0.10–0.47) | 0.55 (0.50–0.60) | 0.20 (0.13–0.27) |
| | UNI | 0.17 | 0.53 | 0.28 | 0.16 | 0.54 | 0.17 | 0.17 (-0.00–0.33) | 0.53 (0.49–0.58) | 0.22 (0.11–0.34) |
| | PLIP | 0.10 | 0.50 | 0.23 | 0.15 | 0.52 | 0.16 | 0.13 (-0.02–0.27) | 0.51 (0.46–0.56) | 0.20 (0.08–0.31) |
| | Virchow | 0.20 | 0.52 | **0.29** | 0.19 | 0.56 | 0.18 | 0.19 (0.02–0.37) | 0.54 (0.49–0.59) | **0.23 (0.12–0.35)** |
| | OmiCLIP | 0.30 | 0.51 | 0.15 | 0.26 | 0.52 | 0.10 | 0.28 (0.12–0.43) | 0.52 (0.45–0.59) | 0.12 (0.07–0.18) |
| | Multi-Embed | 0.26 | 0.49 | 0.10 | 0.23 | 0.52 | 0.08 | 0.25 (0.12–0.37) | 0.50 (0.45–0.56) | 0.09 (0.04–0.14) |
| | SQUALL | **0.34** | **0.53** | 0.18 | **0.49** | **0.70** | 0.13 | **0.42 (0.27–0.56)** | **0.62 (0.58–0.65)** | 0.16 (0.11–0.21) |

**Table S34**: **Virtual spatial gene expression profiling performance across biomarker categories in the comparative evaluation.** Predictions generated at each model's native spatial resolution were subsequently aggregated into 224 × 224-pixel tiles to ensure evaluation at a common spatial resolution. The Pearson correlation coefficient (PCC), area under the curve (AUC), and structural similarity index measure (SSIM) were computed independently for each of the categories, and the resulting gene-wise scores were averaged across genes to obtain mPCC, mAUC, and mSSIM, respectively. The best and second-best values for each metric are shown in bold and underlined, respectively. Values in parentheses indicate the lower and upper bounds corresponding to one standard deviation from the mean. TFs: Transcription factors, LRs: Ligand and receptors, CCs: Cytokine and chemokine, MPs: Membrane proteins.

| Dataset | # Sections | Tissue | Cancer type | Preservation | Metadata |
|---|---|---|---|---|---|
| GSE274051 | 10 | Lymph node | HIV-associated NHL | FFPE | $EBV^+$ (n=5), $EBV^-$ (n=5); biopsies |
| GSE276542 | 9 | Lymph node/Tonsil | DLBCL | FFPE | Tonsil (n=2), LN (2), DLBCL (5); biopsies |
| GSE301720 | 8 | Head and neck | HNSCC | FFPE | Oropharynx(6),nasal cavity(1),oral cavity(1) |
| GSE300445 | 4 | Skin | Melanoma | FFPE | Primary cutaneous melanoma |
| GSE320041 | 15 | Skin | Melanoma | FFPE | Metastatic melanoma |
| GSE322553 | 7 | Colon/Liver/Lung | CRC/HCC/NSCLC | FFPE | CRC (n=4), HCC (n=1), NSCLC (n=2) |
| GSE318562 | 22 | Brain | GBM | FF/FFPE | IDH-wildtype high-grade glioma |
| GSE313694 | 42 | Brain | Meningioma | FF/FFPE | WHO grade 1–3 meningioma |
| **Total** | 117 | 8 tissues | 9 cancer types | FF / FFPE | – |

**Table S35**: **Data characteristics of the virtual spatial gene expression profiling dataset for pretrained-model evaluation (Visium).** To assess the performance of pretrained models in virtual spatial gene expression profiling, we collected newly released Visium sections that became publicly available only after SQUALL pretraining had been completed and the evaluated model weights had been released, thereby minimizing potential information leakage. None of these datasets were included in SQUALL pretraining at either the tissue-block or patient level. All model parameters remained frozen during inference.

| Model | Encoder | Pretraining recipe | PT data | # Para. | Date of release |
|---|---|---|---|---|---|
| OmiCLIP | ViT | Contrastive | ST-bank | 306.7M | 18-Mar-2025 |
| STPath | GigaPath | Generative | HEST-1K + STImage-1k4m | 1.1B | 15-Jun-2025 |
| SQUALL | SQUALL | Generative | *histMol* | 354M | – |

**Table S36**: **Comparison of pretrained histology-spatial transcriptomics foundation models.** We summarize the encoder architecture, pretraining strategy, pretraining data, number of parameters, and release date of each model. Predictions were generated using the respective officially released codebases and following the protocols described in the original publications.

| Cancer type | Dataset | Model | mPCC | mAUC | mSSIM |
|---|---|---|---|---|---|
| HIV-NHL | GSE274051 | OmiCLIP | 0.04 (-0.09–0.18) | 0.51 (0.47–0.55) | 0.24 (0.07–0.41) |
| | | STPath | 0.04 (-0.09–0.18) | 0.51 (0.47–0.56) | 0.25 (0.10–0.39) |
| | | SQUALL | **0.12 (-0.01–0.26)** | **0.54 (0.50–0.58)** | **0.30 (0.15–0.45)** |
| DLBCL | GSE276542 | OmiCLIP | 0.07 (-0.01–0.14) | 0.53 (0.49–0.56) | 0.24 (0.04–0.43) |
| | | STPath | 0.06 (-0.04–0.16) | 0.52 (0.48–0.55) | 0.22 (0.01–0.43) |
| | | SQUALL | **0.17 (0.06–0.27)** | **0.56 (0.52–0.61)** | **0.30 (0.09–0.52)** |
| Head and neck cancer | GSE301720 | OmiCLIP | 0.11 (0.05–0.18) | 0.55 (0.52–0.58) | 0.28 (0.16–0.41) |
| | | STPath | 0.00 (-0.09–0.10) | 0.50 (0.46–0.55) | 0.22 (0.09–0.35) |
| | | SQUALL | **0.13 (0.08–0.19)** | **0.56 (0.53–0.59)** | **0.29 (0.18–0.41)** |
| Primary melanoma | GSE300445 | OmiCLIP | 0.11 (0.00–0.22) | 0.54 (0.51–0.56) | 0.26 (0.19–0.33) |
| | | STPath | 0.24 (0.03–0.45) | 0.60 (0.51–0.69) | 0.37 (0.24–0.51) |
| | | SQUALL | **0.31 (0.16–0.46)** | **0.61 (0.55–0.67)** | **0.40 (0.27–0.53)** |
| Metastatic melanoma | GSE320041 | OmiCLIP | 0.12 (-0.12–0.37) | **0.56 (0.44–0.68)** | 0.15 (-0.06–0.36) |
| | | STPath | 0.10 (-0.15–0.34) | 0.54 (0.44–0.64) | 0.13 (-0.09–0.35) |
| | | SQUALL | **0.12 (-0.10–0.35)** | 0.56 (0.46–0.66) | **0.15 (-0.05–0.35)** |
| NSCLC / HCC / CRC | GSE322553 | OmiCLIP | 0.12 (0.02–0.23) | 0.55 (0.50–0.59) | 0.26 (0.12–0.41) |
| | | STPath | 0.20 (0.12–0.27) | 0.58 (0.55–0.61) | 0.33 (0.25–0.42) |
| | | SQUALL | **0.22 (0.17–0.27)** | **0.59 (0.56–0.61)** | **0.34 (0.23–0.45)** |
| GBM | GSE318562 | OmiCLIP | 0.16 (-0.04–0.35) | 0.56 (0.48–0.65) | 0.33 (0.12–0.55) |
| | | STPath | 0.07 (-0.08–0.22) | 0.53 (0.47–0.59) | 0.30 (0.17–0.43) |
| | | SQUALL | **0.20 (0.05–0.34)** | **0.58 (0.51–0.64)** | **0.37 (0.22–0.52)** |
| Meningioma | GSE313694 | OmiCLIP | 0.12 (-0.00–0.24) | 0.55 (0.49–0.60) | 0.25 (0.10–0.40) |
| | | STPath | 0.03 (-0.09–0.16) | 0.51 (0.45–0.56) | 0.20 (0.07–0.33) |
| | | SQUALL | **0.18 (0.07–0.28)** | **0.57 (0.52–0.61)** | **0.29 (0.15–0.44)** |
| Overall | | OmiCLIP | 0.11 (-0.04–0.27) | 0.55 (0.48–0.62) | 0.26 (0.08–0.43) |
| | | STPath | 0.07 (-0.09–0.22) | 0.52 (0.46–0.59) | 0.23 (0.07–0.39) |
| | | SQUALL | **0.17 (0.04–0.31)** | **0.57 (0.51–0.63)** | **0.30 (0.13–0.46)** |

**Table S37: Virtual spatial gene expression profiling performance of pretrained histology-spatial transcriptomics foundation models (Visium).** OmiCLIP, STPath, and SQUALL were evaluated across diverse cancer types. Predictions generated at each model's native spatial resolution were subsequently aggregated into 224 × 224-pixel tiles to ensure evaluation at a common spatial resolution. The Pearson correlation coefficient (PCC), area under the curve (AUC), and structural similarity index measure (SSIM) were computed independently for each of the 14,766 overlapping genes on Visium sections, and the resulting gene-wise scores were averaged across genes to obtain mPCC, mAUC, and mSSIM, respectively. The best values for each metric are shown in bold. Values in parentheses indicate the lower and upper bounds corresponding to one standard deviation from the mean.

| Dataset | Platform | Tissue | Preservation | Metadata |
|---|---|---|---|---|
| OV-H1 | Visium HD | Ovary | FF | Sex: female; Age: 39 years |
| OV-H2 | Visium HD | Ovary | FF | Sex: female; Age: 65 years |
| Pancreatic cancer | Visium HD | Pancreatic | FF | Sex: female |

**Table S38: Data characteristics of the virtual spatial gene expression profiling dataset for pretrained models (Visium HD).** To assess the performance of pretrained models in virtual spatial gene expression profiling, collected newly released Visium HD sections from 10X Genomics. None of these datasets were included in SQUALL pretraining at either the tissue-block or patient level. All pretrained model parameters remained frozen during inference.

| Model | OV-H1 mPCC | OV-H1 mAUC | OV-H1 mSSIM | OV-H2 mPCC | OV-H2 mAUC | OV-H2 mSSIM | PC mPCC | PC mAUC | PC mSSIM | Overall mPCC | Overall mAUC | Overall mSSIM |
|---|---|---|---|---|---|---|---|---|---|---|---|---|
| UNI | 0.11 | 0.49 | - | 0.03 | 0.47 | - | 0.04 | 0.48 | - | 0.06 (-0.10–0.22) | 0.48 (0.35–0.61) | - |
| PLIP | 0.08 | 0.49 | - | 0.00 | 0.47 | - | 0.00 | 0.47 | - | 0.03 (-0.07–0.13) | 0.47 (0.35–0.60) | - |
| Virchow | 0.14 | 0.48 | - | 0.02 | 0.46 | - | 0.08 | 0.49 | - | 0.08 (-0.08–0.24) | 0.48 (0.35–0.61) | - |
| STPath | 0.32 | 0.49 | 0.12 | 0.33 | 0.53 | 0.29 | 0.34 | 0.58 | 0.18 | 0.33 (0.11–0.55) | 0.53 (0.36–0.70) | 0.20 (0.07–0.32) |
| OmiCLIP | 0.33 | 0.48 | **0.14** | 0.30 | 0.49 | 0.28 | 0.26 | 0.51 | 0.16 | 0.30 (0.08–0.52) | 0.49 (0.31–0.68) | 0.19 (0.06–0.32) |
| SQUALL | **0.34** | **0.50** | 0.13 | **0.34** | **0.53** | **0.30** | **0.35** | **0.59** | **0.19** | **0.34 (0.12–0.57)** | **0.54 (0.37–0.72)** | **0.21 (0.08–0.33)** |

**Table S39**: **Virtual spatial gene expression profiling performance of pretrained foundation models (Visium HD).** SQUALL and STPath were applied directly to the H&E images for inference. For pathology foundation model-based predictors, five-fold cross-validation was performed, and out-of-fold predictions from all held-out test folds were aggregated to summarize overall performance. Predictions were aggregated into 224 × 224-pixel tiles to ensure evaluation at a common spatial resolution. The Pearson correlation coefficient (PCC), area under the curve (AUC), and structural similarity index measure (SSIM) were computed independently for each of the 15,757 overlapping genes on Visium HD sections, and the resulting gene-wise scores were averaged across genes to obtain mPCC, mAUC, and mSSIM, respectively. We do not report mSSIM for pathology foundation model due to its prediction sparseness. The best values for each metric are shown in bold. Values in parentheses indicate the lower and upper bounds corresponding to one standard deviation from the mean.

| Model | Arch. | #Para. | Tech. | Channel | Pretrain data size |
|---|---|---|---|---|---|
| GigaTIME | NestedUNet | 50M | COMET | 21 | 10M cells (21 slides) |
| ROSIE | ConvNet | 50.2M | CODEX | 50 | 16M cells (1,342 slides) |

**Table S40**: **Architecture and pretraining details of spatial proteomics models.** We summarize the implementation details of each pretrained model. All spatial-biomarker prediction evaluations were performed according to the official protocols and preprocessing procedures.

| Category | SQUALL gene | ROSIE marker | SQUALL gene | GigaTIME marker |
|---|---|---|---|---|
| Cell state | *ATM*<br>*BCL2*<br>*MKI67*<br>*PCNA* | ATM<br>BCL2<br>Ki67<br>PCNA | *ACTB*<br>*MKI67* | Actin<br>Ki67 |
| Myeloid / APC | *CD14*<br>*CD163*<br>*ITGAX*<br>*MPO* | CD14<br>CD163<br>CD11c<br>MPO | *CD14*<br>*ITGAX* | CD14<br>CD11c |
| Stromal / Mesenchymal | *ACTA2*<br>*CD34*<br>*PDPN*<br>*VIM* | aSMA<br>CD34<br>Podoplanin<br>Vimentin | *CD34*<br>*TAGLN* | CD34<br>Transgelin |
| T cell | *CD3E*<br>*CD4*<br>*GATA3* | CD3e<br>CD4<br>GATA3 | *CD4*<br>*TBX21* | CD4<br>T-bet |
| B / Plasma cell | *CD38*<br>*CD79A*<br>*CR2*<br>*MS4A1* | CD38<br>CD79a<br>CD21<br>CD20 | *MS4A1*<br>*SDC1* | CD20<br>CD138 |
| Immune state | *CD40*<br>*CD44*<br>*ENTPD1*<br>*ICOS*<br>*IDO1*<br>*LAG3*<br>*LGALS3*<br>*TIGIT*<br>*VSIR* | CD40<br>CD44<br>CD39<br>ICOS<br>IDO1<br>LAG3<br>Gal3<br>TIGIT<br>VISTA | - | - |
| Epithelial | *CDH1*<br>*EPCAM*<br>*TP63* | ECad<br>EpCAM<br>TP63 | - | - |
| Endothelial / Vascular | *PECAM1*<br>*THBD* | CD31<br>CD141 | - | - |
| Immune cell | *PTPRC* | CD45 | - | - |
| **# Total** | **34** | **34** | **10** | **10** |

**Table S41**: **Shared biomarker panels used for virtual biomarker profiling evaluation of SQUALL, ROSIE, and GigaTIME.** We report the genes used for SQUALL and the corresponding protein markers used for ROSIE and GigaTIME in virtual biomarker profiling evaluation. To enable reliable comparison across modalities, protein markers were converted to their corresponding gene symbols using one-to-one gene–protein mappings. Secreted proteins and markers without unambiguous gene correspondence were excluded to ensure structural comparability between gene- and protein-based biomarker panels.

**Panel A: SQUALL vs ROSIE**

| Dataset | Scale | SQUALL | | | ROSIE | | |
|---|---|---|---|---|---|---|---|
| | | mPCC | mAUC | mSSIM | mPCC | mAUC | mSSIM |
| OV-H1 | 224 | 0.33 | 0.50 | 0.19 | **0.34** | **0.52** | **0.19** |
| | 448 | **0.37** | **0.54** | **0.17** | 0.36 | 0.53 | 0.13 |
| OV-H2 | 224 | **0.33** | **0.55** | **0.32** | 0.31 | 0.52 | 0.29 |
| | 448 | **0.43** | **0.58** | **0.32** | 0.41 | 0.54 | 0.30 |
| PC | 224 | **0.35** | **0.58** | **0.19** | 0.24 | 0.46 | 0.12 |
| | 448 | **0.44** | **0.63** | **0.21** | 0.32 | 0.46 | 0.07 |
| **Overall** | 224 | **0.34 (0.29–0.38)** | **0.54 (0.52–0.57)** | **0.23 (0.21–0.26)** | 0.30 (0.26–0.33) | 0.50 (0.47–0.52) | 0.20 (0.18–0.22) |
| | 448 | **0.41 (0.36–0.46)** | **0.58 (0.55–0.61)** | **0.23 (0.20–0.27)** | 0.36 (0.32–0.40) | 0.51 (0.48–0.54) | 0.17 (0.14–0.19) |

**Panel B: SQUALL vs GigaTIME**

| Dataset | Scale | SQUALL | | | GigaTIME | | |
|---|---|---|---|---|---|---|---|
| | | mPCC | mAUC | mSSIM | mPCC | mAUC | mSSIM |
| OV-H1 | 224 | **0.39** | **0.53** | **0.22** | 0.36 | 0.49 | 0.20 |
| | 448 | **0.48** | **0.58** | **0.23** | 0.38 | 0.49 | 0.16 |
| OV-H2 | 224 | **0.37** | **0.57** | **0.32** | 0.33 | 0.53 | 0.30 |
| | 448 | **0.48** | **0.60** | **0.34** | 0.42 | 0.52 | 0.29 |
| PC | 224 | **0.46** | **0.68** | **0.24** | 0.23 | 0.45 | 0.09 |
| | 448 | **0.63** | **0.75** | **0.34** | 0.26 | 0.40 | 0.00 |
| **Overall** | 224 | **0.40 (0.34–0.47)** | **0.59 (0.56–0.62)** | **0.26 (0.22–0.30)** | 0.31 (0.24–0.37) | 0.49 (0.46–0.52) | 0.20 (0.16–0.24) |
| | 448 | **0.53 (0.46–0.60)** | **0.64 (0.60–0.68)** | **0.31 (0.25–0.36)** | 0.35 (0.29–0.42) | 0.47 (0.43–0.52) | 0.15 (0.09–0.22) |

**Table S42**: **Virtual biomarker profiling performance across shared biomarkers.** SQUALL predictions were compared with ROSIE and GigaTIME using the corresponding shared biomarker panels. All models were evaluated at two image scales (224 and 448 pixels) to enable comparisons across spatial scales. Performance was assessed using the mean Pearson correlation coefficient (mPCC), mean area under the curve (mAUC), and mean structural similarity index measure (mSSIM). Confidence intervals are reported for the Overall results. Within each pairwise comparison, the better result is highlighted in bold. Overall performance was calculated by pooling all valid biomarker–dataset metric values, with each valid biomarker–dataset observation contributing equally.

**Panel A: SQUALL vs ROSIE**

| Category | Scale | SQUALL | | | ROSIE | | |
|---|---|---|---|---|---|---|---|
| | | mPCC | mAUC | mSSIM | mPCC | mAUC | mSSIM |
| Cell state | 224 | **0.33** | **0.56** | 0.22 | 0.33 | 0.54 | **0.22** |
| | 448 | **0.39** | **0.59** | **0.18** | 0.38 | 0.56 | 0.17 |
| Myeloid / APC | 224 | **0.37** | **0.55** | **0.25** | 0.30 | 0.48 | 0.19 |
| | 448 | **0.45** | **0.57** | **0.26** | 0.36 | 0.49 | 0.14 |
| Stromal / Mesenchymal | 224 | **0.35** | **0.58** | **0.24** | 0.28 | 0.51 | 0.18 |
| | 448 | **0.45** | **0.62** | **0.26** | 0.36 | 0.52 | 0.14 |
| T cell | 224 | **0.59** | **0.62** | **0.36** | 0.46 | 0.50 | 0.26 |
| | 448 | **0.65** | **0.67** | **0.41** | 0.49 | 0.52 | 0.20 |
| B / Plasma cell | 224 | 0.15 | 0.48 | 0.15 | **0.16** | **0.50** | **0.15** |
| | 448 | 0.20 | 0.49 | 0.10 | **0.23** | **0.51** | **0.11** |
| Immune state | 224 | **0.35** | **0.53** | **0.25** | 0.31 | 0.46 | 0.21 |
| | 448 | **0.40** | **0.59** | **0.23** | 0.34 | 0.47 | 0.16 |
| Epithelial | 224 | **0.36** | **0.55** | **0.22** | 0.35 | 0.53 | 0.22 |
| | 448 | **0.48** | **0.56** | **0.25** | 0.47 | 0.55 | 0.22 |
| Endothelial / Vascular | 224 | **0.12** | **0.43** | **0.13** | 0.12 | 0.42 | 0.13 |
| | 448 | **0.19** | **0.44** | **0.10** | 0.19 | 0.41 | 0.07 |
| Immune cell | 224 | **0.38** | **0.59** | **0.23** | 0.34 | 0.55 | 0.20 |
| | 448 | **0.50** | **0.60** | **0.26** | 0.45 | 0.56 | 0.18 |
| **Overall** | 224 | **0.33 (0.29–0.38)** | **0.54 (0.52–0.56)** | **0.23 (0.21–0.25)** | 0.29 (0.26–0.33) | 0.50 (0.49–0.51) | 0.19 (0.18–0.21) |
| | 448 | **0.41 (0.36–0.46)** | **0.57 (0.55–0.59)** | **0.23 (0.20–0.26)** | 0.36 (0.33–0.40) | 0.51 (0.49–0.53) | 0.15 (0.14–0.17) |

**Panel B: SQUALL vs GigaTIME**

| Category | Scale | SQUALL | | | GigaTIME | | |
|---|---|---|---|---|---|---|---|
| | | mPCC | mAUC | mSSIM | mPCC | mAUC | mSSIM |
| Cell state | 224 | **0.29** | **0.60** | **0.19** | 0.16 | 0.44 | 0.14 |
| | 448 | **0.46** | **0.66** | **0.24** | 0.24 | 0.42 | 0.06 |
| Myeloid / APC | 224 | **0.38** | **0.57** | **0.25** | 0.35 | 0.56 | 0.23 |
| | 448 | **0.48** | **0.61** | **0.26** | 0.40 | 0.55 | 0.21 |
| Stromal / Mesenchymal | 224 | **0.53** | **0.59** | **0.31** | 0.37 | 0.44 | 0.22 |
| | 448 | **0.62** | **0.65** | **0.35** | 0.37 | 0.40 | 0.12 |
| T cell | 224 | **0.57** | **0.61** | **0.34** | 0.44 | 0.48 | 0.25 |
| | 448 | **0.65** | **0.65** | **0.40** | 0.43 | 0.46 | 0.19 |
| B / Plasma cell | 224 | **0.27** | **0.58** | **0.19** | 0.23 | 0.52 | 0.17 |
| | 448 | **0.43** | **0.63** | **0.23** | 0.33 | 0.54 | 0.17 |
| **Overall** | 224 | **0.41 (0.34–0.47)** | **0.59 (0.58–0.60)** | **0.26 (0.23–0.29)** | 0.31 (0.26–0.36) | 0.49 (0.46–0.51) | 0.20 (0.18–0.22) |
| | 448 | **0.53 (0.48–0.57)** | **0.64 (0.63–0.65)** | **0.29 (0.26–0.33)** | 0.35 (0.32–0.39) | 0.47 (0.44–0.50) | 0.15 (0.12–0.18) |

**Table S43**: **Virtual biomarker profiling performance across shared biomarker categories.** SQUALL predictions were compared with those of ROSIE and GigaTIME using the corresponding shared biomarker panels. All models were evaluated using two image-tile sizes ($224 \times 224$ and $448 \times 448$ pixels) to enable comparisons across spatial scales. Performance within each biomarker category was assessed using the mean Pearson correlation coefficient (mPCC), mean area under the curve (mAUC), and mean structural similarity index measure (mSSIM). Confidence intervals are reported for the Overall results. Within each pairwise comparison, the better-performing result is shown in bold. Overall performance was calculated as an unweighted macro-average across category-level means, with each biomarker category contributing equally regardless of its number of biomarkers.

| Cat. | Tissue | Species | Dataset | Preser. | #Samp. | #Sect. | Path. cond. |
|---|---|---|---|---|---|---|---|
| Ann | Liver | H. sapiens | GSE238264 | FF | 2 | 2 | HCC neoadjuvant responder, HCC neoadjuvant nonresponder |
| | Breast | H. sapiens | GSE195665 | FF | 10 | 10 | Contra(2), Left(3), Right(5) |
| | Lung | H. sapiens | E-MTAB-13530 | FF | 8 | 16 | LUAD(8), LUSC(5), NSCLC(2), Normal(1) |
| | Head and neck | H. sapiens | GSE200310 | FF | 2 | 2 | Nasopharyngeal carcinoma(2) |
| | Cervix | H. sapiens | GSE208654 | FF | 7 | 7 | HGSIL HPV+, CESC HPV+, Normal HPV+ |
| | Soft tissue | H. sapiens | GSE212526 | FF | 4 | 4 | Undiff. pleomorphic sarcoma (3) Leiomyosarcoma(1) |
| | Multiple | H. sapiens | GSE224411 | FFPE/FF | 3 | 3 | PDAC lymphnode(1,FF) PaIN (1,FFPE) HCC(1,FF) |
| Ex | Thyroid (head and neck) | H. sapiens | GSE250521 | FF | 12 | 12 | Normal(2), LPTC(2), PT(4), PTC(4) |
| | Breast | M. musculus | GSE230098 | FFPE | 4 | 4 | 4T1 tumors were orthotopically implanted into BALB/C mice. |

**Table S44**: **Clinical characteristics of the spatially aware clustering dataset.** Both annotated sections and external Visium sections were used to evaluate model performance on the spatially aware clustering task across various tumor and normal conditions. Annotated (Ann) sections refer to those used during SQUALL pretraining and were annotated by pathologists using histology images or supplemented with marker gene expressions to provide reference labels for evaluation. External (Ex) sections are from patients whose data were not seen by SQUALL during pretraining, allowing us to assess SQUALL's clinical and biological relevant hypothesis generation ability.

| Method | Modality | Preprocessing | Feature extractor | Fusion method | Clustering |
|---|---|---|---|---|---|
| UNI<br>Scanpy | Image<br>ST | ImageNet normalization<br>Log-normalization | UNI<br>PCA | -<br>- | k-means<br>Leiden |
| MISO | Image+ST | Log-normalization<br>Z-score normalization | HIPT (TCGA)<br>SpectralNet | Kronecker product | k-means |
| SpatialGlue | Image+ST | Log-normalization<br>Z-score normalization | GCN | Gated attention | mclust |
| OmiCLIP | Pretrained multimodal embedding | Scanpy/Seurat pipeline | Loki (ST-BANK) | Contrastive alignment | k-means |
| STPath | Pretrained multimodal embedding | Aggregating into pseudo-spots | GigaPath | Generative alignment | Leiden |
| SQUALL | Image+ST | Pseudo-labeling<br>Background correction | SQUALL Encoder | Cross-attention | k-means |

**Table S45**: **Summary of implementation details of spatially aware clustering models.** We present the structural differences among spatially aware clustering models. UNI and Scanpy were included as single-modality baselines, whereas OmiCLIP and STPath were included as multimodal embedding methods for evaluation. All protocols were adopted from the original codebases without modifications. For SQUALL, the final normalization layer was removed to obtain spot-level representations for downstream clustering.

| Tissue | Dataset | Model | ARI | AMI | NMI | CHAOS |
|---|---|---|---|---|---|---|
| Liver | GSE238264 | Scanpy | 0.10 (0.01–0.17) | 0.19 (0.09–0.30) | 0.19 (0.10–0.30) | 0.07 (0.07–0.07) |
| | | UNI | 0.13 (0.04–0.19) | 0.22 (0.15–0.28) | 0.22 (0.16–0.29) | 0.09 (0.09–0.09) |
| | | MISO | 0.17 (0.09–0.25) | 0.21 (0.17–0.23) | 0.21 (0.17–0.23) | 0.08 (0.08–0.08) |
| | | SpatialGlue | 0.10 (0.02–0.20) | 0.15 (0.08–0.24) | 0.16 (0.08–0.24) | **0.06 (0.06–0.07)** |
| | | OmiCLIP | 0.16 (0.04–0.23) | 0.26 (0.12–0.35) | 0.27 (0.13–0.36) | 0.07 (0.07–0.08) |
| | | STPath | 0.08 (0.03–0.12) | **0.29 (0.25–0.37)** | **0.30 (0.25–0.38)** | **0.06 (0.05–0.06)** |
| | | SQUALL | **0.20 (0.05–0.33)** | 0.28 (0.12–0.37) | 0.29 (0.12–0.37) | 0.07 (0.06–0.07) |
| Breast | GSE195665 | Scanpy | 0.06 (0.02–0.09) | 0.16 (0.09–0.20) | 0.16 (0.10–0.20) | 0.10 (0.08–0.11) |
| | | UNI | 0.11 (0.07–0.15) | 0.23 (0.18–0.29) | 0.23 (0.19–0.29) | 0.09 (0.08–0.11) |
| | | MISO | 0.08 (0.02–0.17) | 0.15 (0.08–0.23) | 0.16 (0.08–0.23) | 0.07 (0.06–0.09) |
| | | SpatialGlue | 0.12 (0.03–0.25) | 0.16 (0.04–0.28) | 0.17 (0.05–0.28) | 0.07 (0.06–0.08) |
| | | OmiCLIP | 0.16 (0.11–0.20) | **0.25 (0.22–0.28)** | **0.26 (0.22–0.28)** | 0.08 (0.07–0.10) |
| | | STPath | 0.02 (0.01–0.03) | 0.09 (0.06–0.16) | 0.10 (0.07–0.17) | 0.06 (0.06–0.08) |
| | | SQUALL | **0.16 (0.07–0.35)** | 0.20 (0.12–0.39) | 0.20 (0.12–0.39) | **0.06 (0.06–0.06)** |
| Lung | E-MTAB-13530 | Scanpy | 0.06 (0.01–0.18) | 0.13 (0.03–0.30) | 0.13 (0.04–0.31) | 0.10 (0.08–0.13) |
| | | UNI | 0.05 (0.0–0.10) | 0.10 (0.01–0.16) | 0.10 (0.01–0.17) | 0.11 (0.08–0.14) |
| | | MISO | 0.07 (-0.01–0.20) | 0.11 (0.0–0.22) | 0.12 (0.01–0.23) | 0.09 (0.07–0.13) |
| | | SpatialGlue | 0.08 (-0.02–0.23) | 0.13 (0.02–0.33) | 0.14 (0.02–0.33) | **0.08 (0.06–0.12)** |
| | | OmiCLIP | 0.08 (0.01–0.15) | 0.12 (0.02–0.20) | 0.12 (0.03–0.21) | 0.10 (0.08–0.14) |
| | | STPath | 0.04 (0.00–0.10) | **0.18 (0.07–0.30)** | **0.18 (0.08–0.31)** | **0.08 (0.06–0.12)** |
| | | SQUALL | **0.08 (0.02–0.20)** | 0.14 (0.03–0.27) | 0.14 (0.03–0.28) | 0.10 (0.07–0.14) |
| Head and neck | GSE200310 | Scanpy | 0.01 (0.00–0.03) | 0.07 (0.02–0.11) | 0.07 (0.03–0.12) | 0.11 (0.10–0.11) |
| | | UNI | 0.03 (0.01–0.04) | 0.09 (0.06–0.11) | 0.09 (0.07–0.11) | 0.13 (0.13–0.14) |
| | | MISO | 0.02 (0.02–0.03) | 0.07 (0.05–0.08) | 0.07 (0.06–0.09) | 0.11 (0.11–0.12) |
| | | SpatialGlue | 0.04 (0.01–0.07) | 0.08 (0.03–0.13) | 0.08 (0.03–0.13) | **0.09 (0.09–0.10)** |
| | | OmiCLIP | 0.04 (0.02–0.06) | 0.09 (0.06–0.12) | 0.10 (0.07–0.13) | 0.12 (0.11–0.13) |
| | | STPath | 0.01 (0.00–0.01) | 0.06 (0.03–0.10) | 0.07 (0.03–0.11) | **0.09 (0.09–0.10)** |
| | | SQUALL | **0.05 (0.02–0.08)** | **0.10 (0.07–0.14)** | **0.11 (0.07–0.15)** | 0.11 (0.10–0.12) |
| Cervix | GSE208654 | Scanpy | 0.05 (0.01–0.12) | 0.14 (0.05–0.28) | 0.15 (0.07–0.28) | 0.13 (0.09–0.18) |
| | | UNI | 0.10 (0.02–0.18) | 0.23 (0.09–0.36) | 0.24 (0.11–0.36) | 0.15 (0.11–0.24) |
| | | MISO | 0.08 (0.00–0.20) | 0.15 (0.04–0.29) | 0.16 (0.06–0.30) | 0.13 (0.10–0.20) |
| | | SpatialGlue | 0.07 (0.01–0.16) | 0.12 (0.04–0.24) | 0.13 (0.05–0.24) | **0.12 (0.09–0.17)** |
| | | OmiCLIP | **0.17 (0.06–0.31)** | **0.25 (0.13–0.39)** | **0.26 (0.14–0.39)** | 0.14 (0.10–0.21) |
| | | STPath | 0.04 (0.01–0.08) | 0.18 (0.08–0.29) | 0.19 (0.11–0.30) | **0.12 (0.09–0.18)** |
| | | SQUALL | 0.12 (0.03–0.24) | 0.24 (0.12–0.37) | 0.25 (0.14–0.37) | 0.13 (0.10–0.19) |
| Soft tissue | GSE212526 | Scanpy | 0.08 (0.00–0.22) | 0.14 (0.00–0.37) | 0.14 (0.00–0.37) | 0.10 (0.08–0.11) |
| | | UNI | 0.09 (0.01–0.26) | 0.19 (0.04–0.42) | 0.19 (0.05–0.43) | 0.11 (0.10–0.13) |
| | | MISO | 0.16 (0.02–0.30) | 0.16 (0.05–0.30) | 0.16 (0.05–0.31) | 0.10 (0.09–0.10) |
| | | SpatialGlue | 0.14 (0.00–0.39) | 0.13 (0.00–0.35) | 0.14 (0.01–0.35) | 0.09 (0.07–0.11) |
| | | OmiCLIP | 0.11 (0.02–0.24) | 0.22 (0.08–0.41) | 0.22 (0.08–0.41) | 0.10 (0.09–0.12) |
| | | STPath | 0.03 (0.00–0.09) | 0.15 (0.02–0.29) | 0.15 (0.03–0.30) | 0.08 (0.07–0.10) |
| | | SQUALL | **0.27 (0.09–0.45)** | **0.27 (0.09–0.51)** | **0.27 (0.09–0.51)** | **0.05 (0.05–0.05)** |
| Multiple | GSE224411 | Scanpy | 0.11 (0.05–0.18) | 0.21 (0.08–0.32) | 0.22 (0.09–0.33) | 0.09 (0.07–0.11) |
| | | UNI | 0.07 (0.01–0.17) | 0.10 (0.02–0.21) | 0.11 (0.03–0.21) | 0.11 (0.09–0.12) |
| | | MISO | 0.08 (0.01–0.19) | 0.13 (0.05–0.22) | 0.14 (0.06–0.23) | 0.09 (0.07–0.11) |
| | | SpatialGlue | 0.11 (0.05–0.21) | 0.15 (0.08–0.27) | 0.15 (0.09–0.27) | 0.08 (0.07–0.10) |
| | | OmiCLIP | 0.11 (0.04–0.23) | 0.16 (0.08–0.30) | 0.17 (0.09–0.30) | 0.09 (0.07–0.11) |
| | | STPath | 0.10 (0.08–0.11) | **0.31 (0.26–0.36)** | **0.32 (0.27–0.37)** | 0.08 (0.06–0.09) |
| | | SQUALL | **0.16 (0.06–0.33)** | 0.21 (0.11–0.38) | 0.22 (0.12–0.38) | **0.08 (0.07–0.10)** |
| **Overall** | All cohorts | Scanpy | 0.07 (0.01–0.11) | 0.15 (0.07–0.21) | 0.15 (0.07–0.22) | 0.10 (0.07–0.13) |
| | | UNI | 0.08 (0.03–0.13) | 0.16 (0.09–0.23) | 0.17 (0.09–0.24) | 0.11 (0.09–0.15) |
| | | MISO | 0.10 (0.02–0.17) | 0.14 (0.07–0.21) | 0.15 (0.07–0.21) | 0.10 (0.07–0.13) |
| | | SpatialGlue | 0.09 (0.04–0.14) | 0.13 (0.08–0.16) | 0.14 (0.08–0.17) | 0.09 (0.06–0.12) |
| | | OmiCLIP | 0.12 (0.04–0.17) | 0.19 (0.09–0.26) | 0.20 (0.10–0.27) | 0.10 (0.07–0.14) |
| | | STPath | 0.04 (0.01–0.10) | 0.18 (0.06–0.31) | 0.19 (0.07–0.32) | **0.08 (0.06–0.12)** |
| | | SQUALL | **0.15 (0.05–0.27)** | **0.21 (0.10–0.28)** | **0.21 (0.11–0.29)** | 0.09 (0.05–0.13) |

**Table S46**: **Spatially aware clustering performance for annotated dataset.**

**Legend for Table S46 (continued).** For each section, spatial clusters were derived by the corresponding methods. With reference annotations, model performance was evaluated using reference-based clustering metrics: ARI (adjusted Rand index), AMI (adjusted mutual information), NMI (normalized mutual information), and unsupervised metrics, CHAOS; however, spatial clusters in cancer sections are not necessarily spatially smooth. Therefore, CHAOS should be interpreted as a complementary metric rather than a definitive measure of clustering performance. The best and second-best metrics are highlighted in bold and underlined, respectively, with the minimum and maximum values within each cohort or across datasets provided in parentheses.

| Dataset | Tissue | SQUALL vs. | Win (%) | Loss (%) |
|---|---|---|---|---|
| GSE250521 | Thyroid | Scanpy | 80.00 (68.2–88.2) | 20.00 (11.8–31.8) |
| | | UNI | 95.00 (86.3–98.3) | 5.00 (1.7–13.7) |
| | | MISO | 96.67 (88.6–99.1) | 3.33 (0.9–11.4) |
| | | SpatialGlue | 98.33 (91.1–99.7) | 1.67 (0.3–8.9) |
| | | OmiCLIP | 96.67 (88.6–99.1) | 3.33 (0.9–11.4) |
| | | STPath | 100.00 (94.0–100.0) | 0.00 (0.0–6.0) |
| GSE230098 | Breast | Scanpy | 100.00 (83.9–100.0) | 0.00 (0.0–16.1) |
| | | UNI | 90.00 (69.9–97.2) | 10.00 (2.8–30.1) |
| | | MISO | 100.00 (83.9–100.0) | 0.00 (0.0–16.1) |
| | | SpatialGlue | 100.00 (83.9–100.0) | 0.00 (0.0–16.1) |
| | | OmiCLIP | 70.00 (48.1–85.5) | 30.00 (14.5–51.9) |
| | | STPath | 100.00 (83.9–100.0) | 0.00 (0.0–16.1) |

**Table S47**: **Spatially aware clustering for external datasets.** Five pathologists independently ranked annotation outputs without knowledge of the underlying methods; rank 1 was best. A win means SQUALL received a better rank than the comparison method. A loss includes worse ranks. Values are percentages; 95% confidence intervals are shown in parentheses. Baseline settings are indicated in gray.

| Pathologist | SQUALL vs. | Win (%) | Loss (%) |
|---|---|---|---|
| 1 | Scanpy | 75.00 (50.5–89.8) | 25.00 (10.2–49.5) |
| | UNI | 93.75 (71.7–98.9) | 6.25 (1.1–28.3) |
| | MISO | 100.00 (80.6–100.0) | 0.00 (0.0–19.4) |
| | SpatialGlue | 93.75 (71.7–98.9) | 6.25 (1.1–28.3) |
| | OmiCLIP | 87.50 (64.0–96.5) | 12.50 (3.5–36.0) |
| | STPath | 100.00 (80.6–100.0) | 0.00 (0.0–19.4) |
| 2 | Scanpy | 87.50 (64.0–96.5) | 12.50 (3.5–36.0) |
| | UNI | 93.75 (71.7–98.9) | 6.25 (1.1–28.3) |
| | MISO | 93.75 (71.7–98.9) | 6.25 (1.1–28.3) |
| | SpatialGlue | 100.00 (80.6–100.0) | 0.00 (0.0–19.4) |
| | OmiCLIP | 87.50 (64.0–96.5) | 12.50 (3.5–36.0) |
| | STPath | 100.00 (80.6–100.0) | 0.00 (0.0–19.4) |
| 3 | Scanpy | 68.75 (44.4–85.8) | 31.25 (14.2–55.6) |
| | UNI | 100.00 (80.6–100.0) | 0.00 (0.0–19.4) |
| | MISO | 93.75 (71.7–98.9) | 6.25 (1.1–28.3) |
| | SpatialGlue | 100.00 (80.6–100.0) | 0.00 (0.0–19.4) |
| | OmiCLIP | 93.75 (71.7–98.9) | 6.25 (1.1–28.3) |
| | STPath | 100.00 (80.6–100.0) | 0.00 (0.0–19.4) |
| 4 | Scanpy | 93.75 (71.7–98.9) | 6.25 (1.1–28.3) |
| | UNI | 93.75 (71.7–98.9) | 6.25 (1.1–28.3) |
| | MISO | 100.00 (80.6–100.0) | 0.00 (0.0–19.4) |
| | SpatialGlue | 100.00 (80.6–100.0) | 0.00 (0.0–19.4) |
| | OmiCLIP | 93.75 (71.7–98.9) | 6.25 (1.1–28.3) |
| | STPath | 100.00 (80.6–100.0) | 0.00 (0.0–19.4) |
| 5 | Scanpy | 100.00 (80.6–100.0) | 0.00 (0.0–19.4) |
| | UNI | 87.50 (64.0–96.5) | 12.50 (3.5–36.0) |
| | MISO | 100.00 (80.6–100.0) | 0.00 (0.0–19.4) |
| | SpatialGlue | 100.00 (80.6–100.0) | 0.00 (0.0–19.4) |
| | OmiCLIP | 87.50 (64.0–96.5) | 12.50 (3.5–36.0) |
| | STPath | 100.00 (80.6–100.0) | 0.00 (0.0–19.4) |

**Table S48**: **Spatially aware clustering for external datasets as evaluated by five individual pathologists.** Each pathologist ranked 16 sections. A win means SQUALL received a better rank than the comparison method; a loss includes worse ranks. Values are percentages with 95% confidence intervals in parentheses. Baseline settings are indicated in gray.

| *histMol*-low: Breast cancer (N=103, S=198) | | | | |
|---|---|---|---|---|
| **Pathological type** | **N** | **N (%)** | **S** | **S (%)** |
| Ductal Carcinoma In Situ | 6 | 5.8% | 11 | 5.6% |
| Invasive Ductal Carcinoma | 38 | 36.9% | 72 | 36.4% |
| Invasive Lobular Carcinoma | 4 | 3.9% | 10 | 5.1% |
| Invasive Micropapillary Carcinoma | 7 | 6.8% | 16 | 8.1% |
| Metaplastic Breast Carcinoma | 16 | 15.5% | 18 | 9.1% |
| Mixed Carcinoma | 1 | 1.0% | 1 | 0.5% |
| Medullary Carcinoma of Breast | 1 | 1.0% | 3 | 1.5% |
| No Special Type | 30 | 29.1% | 67 | 33.8% |
| **Technical platform** | | | | |
| Visium | 70 | 68.0% | 111 | 56.1% |
| Spatial Transcriptomics | 33 | 32.0% | 87 | 43.9% |
| **Curated molecular subtype** | | | | |
| Luminal A | 9 | 8.7% | 15 | 7.6% |
| Luminal B | 21 | 20.4% | 46 | 23.2% |
| HER2-enriched | 12 | 11.7% | 32 | 16.2% |
| TNBC | 49 | 47.6% | 77 | 38.9% |
| NA | 12 | 11.7% | 28 | 14.1% |
| **Predicted PAM50 subtype*** | | | | |
| Luminal A | 12 | 11.7% | 22 | 11.1% |
| Luminal B | 22 | 21.4% | 36 | 18.2% |
| HER2-enriched | 18 | 17.5% | 42 | 21.2% |
| Basal-like | 49 | 47.6% | 75 | 37.9% |
| Normal-like | 8 | 7.8% | 16 | 8.1% |
| NA | 7 | 6.8% | 7 | 3.5% |

**Table S49**: **Cohort characteristics of the *histMol*-low breast cancer corpus used for analysis.** All metadata were curated and harmonized from the original publications. Pathological subtype annotations were reviewed by three clinical pathologists based on available pathological images and clinical records. Final labels were assigned through consensus review. Cases without sufficient evidence for further pathological subclassification were retained as No Special Type (NST) when applicable. For PAM50 subtype assignment, spatial transcriptomic raw count matrices were aggregated into pseudo-bulk expression profiles, and subtype classification was performed using the `genefu` R package. Patient-level percentages were calculated using the total number of patients as the denominator ($N = 103$), whereas section-level percentages were calculated using the total number of tissue sections as the denominator ($S = 198$). Because multiple sections from the same patient may receive different PAM50 subtype assignments, an individual patient may be counted in more than one PAM50 subtype row. Consequently, the PAM50 patient counts and their corresponding $N$-based percentages are not mutually exclusive and should not be summed vertically. In contrast, each section contributes to only one PAM50 subtype category, including NA; therefore, the PAM50 section counts and their corresponding $S$-based percentages sum to $S = 198$ and 100%, respectively.

| **Instruction** | **Rationale** |
|---|---|
| Emphasize the importance of accurate alignment with human-recognizable features identified by pathologists | Pathologist-defined annotations may not always correspond one-to-one with detailed subtyping based on molecular features. One goal of multimodal modeling is to provide an unsupervised and efficient method to recovery spatial tissue feature difference in a unsupervised manner as well as subsequently reveals these differences to identified novel molecular-pathology subtype or clusters. Ensuring alignment with human-recognizable features can help with model evaluations, as well as give more consistent and reliable association analysis. |
| Provide resolution information about H&E images as well as sequencing resolution. | This enables participants to determine the appropriate level of annotation granularity, facilitates accurate identification and categorization of pathological features, enhancing the precision and reliability of analysis. |
| Provide H&E stained images with background correction | H&E images from public sources often show varying imaging conditions and exposure intensities, which can impede feature recognition. Background correction ensures uniformity, improves image clarity, and enhances the identification of critical pathological structures. |
| Provide detailed metadata, key marker genes spatial expression as much as possible. Include original annotations from authors if available. | Comprehensive metadata aids pathologists in selecting appropriate annotation classes and locating specific features, particularly in low-magnification images. Original annotations offer valuable context and further improve the accuracy of analysis. Also provides key marker gene expression overlay with pathological images if needed. |

**Table S50**: **Spatial cluster and tissue annotation guidelines.** Our experiences during collaboration with pathologists in annotating *histMol* data are outlined. We detail 4 instructions and the corresponding rationales that we found to facilitate a smooth annotation process. We believe these guidelines can make communication more seamless during future collaborations with pathologists.

| Major class | Minor class | Polygon counts |
|---|---|---|
| Normal parenchyma | Alveolus | 394 |
| | Interlobular duct | 226 |
| | Glandular duct | 80 |
| Tumor parenchyma | Tumor cells | 1,524 |
| | Infiltrating carcinoma | 1,188 |
| | Carcinoma in situ | 161 |
| | Necrotic and receding cancer cells | 113 |
| | Necrotic tumor cells | 23 |
| | Scattered tumor cells with immune cells infiltration | 19 |
| Stroma | Generic stroma | 2,208 |
| | Stroma with high immune infiltration | 299 |
| | Fibrotic stroma | 17 |
| Interstitial | Adipocyte | 1,211 |
| | Immune cells | 102 |
| | Blood vessel | 43 |
| | Vascular endothelial cells | 17 |
| | Loose interstitial tissue | 7 |
| | Endothelial cells | 5 |
| Tumor-stroma mixture | Interstitial and infiltrating cancer cells | 767 |
| | Scattered cancer cells in stroma | 87 |
| Tumor-interstitial mixture | Adipose tissue and infiltrating cancer cells | 127 |
| Others | Unknown | 84 |
| | Calcification | 34 |
| | Tissue lamination | 6 |
| Total | | 8,742 |

**Table S51**: **Annotation hierarchy and summary for breast cancer subset of *histMol*-low.** Three to five experienced pathologists were involved in the tissue annotation process. Categories were merged based on their consensus. All annotations were formatted as polygon or multi-polygon objects in GeoJSON. Each spot was assigned a label based on its intersection with the polygon. Spots located on the boundary or lacking distinguishable features were assigned to the "Others/NA" category.

| ***histMol*-high: Ovarian tumor (N=52)** | |
|---|---|
| **Pathological type** | |
| High-Grade Serous Ovarian Carcinoma | 34 (65.4%) |
| Ovarian Clear Cell Carcinoma | 4 (7.7%) |
| Ovarian Endometrioid Carcinoma | 5 (9.6%) |
| Ovarian Mucinous Cystadenoma | 3 (5.8%) |
| High-Grade Ovarian Squamous Cell Carcinoma | 1 (1.9%) |
| Ovarian Endometrioid Adenocarcinoma | 1 (1.9%) |
| Ovarian Granulosa Cell Tumor | 1 (1.9%) |
| Mixed Germ Cell Tumor | 1 (1.9%) |
| Others | 2 (3.8%) |
| **RECIST response** | |
| Complete Response | 30 (57.7%) |
| Partial Response | 8 (15.4%) |
| NA | 14 (26.9%) |
| **Recurrence status** | |
| Yes | 29 (55.8%) |
| No | 23 (44.2%) |
| **Survival status** | |
| Deceased | 12 (23.1%) |
| Alive | 40 (76.9%) |

**Table S52**: **Clinical characteristics of the *histMol*-high primary ovarian tumor component used for analysis.** All clinical metadata were curated and cleaned from the original patients' medical records. Paired histological images were scanned and digitized using a Leica Aperio GT 450 scanner at 40× magnification prior to sequencing.

| Config. | Survival prediction | Platinum resistance prediction |
|---|---|---|
| Hidden dim. | 256 | |
| Attention type | Gated Attention | |
| Dropout rate | 0.25 | |
| Optimization target | DeepSurv | Cross-entropy |
| Optimizer | AdamW | |
| Optimizer momentum | $\beta_1 = 0.9$, $\beta_2 = 0.999$ | |
| Learning rate | 2e-4 | |
| Learning rate schedule | Constant | |
| Maximum epochs | 50 | |
| Batch size | 1 | |
| Gradient accu. steps | 64 | |
| Early stopping | ✓ | |
| Patience | 30 | |
| Model-selection metric | C-index | Accuracy |

**Table S53**: **Configuration for clinically relevant downstream prediction tasks.** An NVIDIA H100 GPU was used for all downstream clinical-task evaluations. All models were trained using the same training recipe to ensure fair, head-to-head comparisons. For each slide, all associated tiles were forwarded through the model. To stabilize the gradients and simulate large-batch training, we employed gradient accumulation, allowing multiple sections to be processed before a single backward pass. Early stopping was applied to prevent overfitting.

| Model | Arch. | Embd. level | SSL recipe | Mag. | Input size | Embd. size | #Para. | Modality |
|---|---|---|---|---|---|---|---|---|
| UNI | ViT-large | Tile | DINOv2 | 20× | 256×256×3 | 16×16×1,024 | 307M | Vision |
| PLIP | ViT-base | Tile | CLIP | various | 224×224×3 | 32×32×512 | 86M | Vision-Language |
| Virchow | ViT-huge | Tile (Slide) | DINOv2 | 20× | 224×224×3 | 1×1×2,560 | 632M | Vision |
| SQUALL | ViT-large | Tile | MAE* | 20× | 224×224×3 | 14×14×1,024 | 303M | Vision-ST |

**Table S54**: **Comparison of implementation details of pretrained vision encoders.** We present implementation details of each pretrained encoder. All publicly available model weights were accessed at Nov. 15th, 2024. All downstream clinically relevant tasks were evaluated following their official protocols and whole slide image (WSI) preprocessing procedures. For UNI, WSI preprocessing was performed using the CLAM toolbox. The total number of parameters (#Para.) refers only to those in the vision encoder. For Virchow, tile-level encoder were used to obtain the final WSI embedding. For SQUALL, the self-supervised learning (SSL) recipe was within the MAE family.

| Model | Pretrain source | Data scale (Tile, Slides) | Epoch | Batch size | Time | Resources |
|---|---|---|---|---|---|---|
| UNI | BWH, MGH, GTEx | 100.1M, 100.1K | 3.85 | 3,072 | 32 hrs | 32× A100s |
| PLIP | OpenPath (WIT400M) | 0.2M (400M) Text-Image pairs | 10 | 128 | <10 hrs | Multiple L40Ss |
| Virchow | MSKCC | 2B, 1.5M | 2.3 | 3,072 | ∼22.5 hrs | 16×V100s |
| SQUALL | *histMol* | 0.7M, 3.4K | 400/100 | 128/256 | 177.3/26.8 hrs | 32× H100s |

**Table S55**: **Comparison of training details of pretrained vision encoders.** We present the training details of each pretrained encoder. All information was curated from the original publications or official codebases. The number of training epochs was estimated based on the reported batch size and iteration count. For PLIP, we also included the WIT-400M dataset, as the original CLIP was trained on it; the data scale is reported in parentheses. For SQUALL, training resources are reported in the format 'stage 1 / stage 2', indicating different training phases. Please note that SQUALL employed a MAE-like training recipe, which requires a longer training schedule than DINOv2 due to its reconstruction-based objective and multi-modal input. Despite the higher training cost, this approach enables more comprehensive representation learning across both histology and transcriptomic modalities. As paired histology image–spatial transcriptomics datasets are still being accumulated and have not yet reached full coverage, the pretrained weights of SQUALL can serve as effective starting points for further model development.

| CGODC-OV (N=85) | |
|---|---|
| **Pathological type** | |
| High-Grade Serous Ovarian Carcinoma | 68 (80.0%) |
| Ovarian Clear Cell Carcinoma | 4 (4.7%) |
| Low-Grade Serous Ovarian Carcinoma | 3 (3.5%) |
| Mucinous Ovarian Carcinoma | 2 (2.4%) |
| High-Grade Mucinous Ovarian Carcinoma | 2 (2.4%) |
| Granulosa Cell Tumor | 2 (2.4%) |
| Mixed Germ Cell Tumor | 1 (1.2%) |
| Endometrioid Ovarian Carcinoma | 2 (2.4%) |
| Small Cell Carcinoma | 1 (1.2%) |
| **FIGO stage** | |
| Stage I | 6 (7.1%) |
| Stage II | 3 (3.5%) |
| Stage III | 50 (58.8%) |
| Stage IV | 13 (15.3%) |
| NA | 13 (15.3%) |
| **Platinum status** | |
| Sensitive | 67 (78.8%) |
| Resistance | 18 (21.2%) |
| **Survival status** | |
| Deceased | 35 (41.2%) |
| Alive | 50 (58.8%) |

**Table S56**: **Clinical characteristics of the CGODC-OV dataset.** Diagnostic H&E-stained tissue sections from the patients were collected and digitized using an Olympus SLIDEVIEW VS200 scanner. Detailed clinical metadata were curated from the original medical records. This cohort was subsequently used to evaluate the performance of pretrained models on survival prediction and platinum-based chemotherapy resistance prediction tasks.

| TCGA-CESC (N=269) | |
|---|---|
| **Pathological type** | |
| Squamous Cell Neoplasms | 219 (81.4%) |
| Adenomas and Adenocarcinomas | 27 (10.0%) |
| Cystic, Mucinous and Serous Neoplasms | 17 (6.3%) |
| Complex Epithelial Neoplasms | 6 (2.2%) |
| **FIGO stage** | |
| Stage I | 140 (52.0%) |
| Stage II | 61 (22.7%) |
| Stage III | 42 (15.6%) |
| Stage IV | 20 (7.4%) |
| NA | 6 (2.2%) |
| **HPV status** | |
| Positive | 138 (51.3%) |
| Negative | 8 (3.0%) |
| NA | 123 (45.7%) |
| **Survival status** | |
| Deceased | 65 (24.2%) |
| Alive | 204 (75.8%) |

**Table S57**: **Clinical characteristics of the TCGA-CESC dataset.** Only slides labeled as diagnostic (DX) were used for model training and evaluation. For patients with more than one DX slide, one was randomly selected for use in training and evaluation.

| TCGA-STAD (N=416) | |
|---|---|
| **AJCC stage** | |
| Stage I | 53 (12.7%) |
| Stage II | 121 (29.1%) |
| Stage III | 176 (42.3%) |
| Stage IV | 43 (10.3%) |
| NA | 23 (5.5%) |
| **Survival status** | |
| Deceased | 164 (39.4%) |
| Alive | 247 (59.4%) |
| Unknown | 5 (1.2%) |

**Table S58**: **Clinical characteristics of the TCGA-STAD dataset.** Only slides labeled as diagnostic (DX) were used for model training and evaluation. For patients with more than one DX slide, one was randomly selected for use in training and evaluation.

| TCGA-OV (N=106) | |
|---|---|
| **FIGO stage** | |
| Stage I | 2 (1.9%) |
| Stage II | 4 (3.8%) |
| Stage III | 74 (69.8%) |
| Stage IV | 24 (22.6%) |
| NA | 2 (1.9%) |
| **Platinum status** | |
| Sensitive | 27 (25.5%) |
| Resistance | 15 (14.2%) |
| Too early | 6 (5.7%) |
| Missing | 58 (54.7%) |
| **Survival status** | |
| Deceased | 72 (67.9%) |
| Alive | 33 (31.1%) |
| Unknown | 1 (0.9%) |

**Table S59**: **Clinical characteristics of the TCGA-OV dataset.** Of note, all patients within this cohort are patients with high-grade serous ovarian carcinomas. Only slides labeled as diagnostic (DX) were used for model training and evaluation. For patients with more than one DX slide, one was randomly selected for use in training and evaluation.

| Cohort | SQUALL | UNI | PLIP | Virchow |
|---|---|---|---|---|
| CGODC-OV | **0.621 (0.492–0.714)** | 0.542 (0.429–0.654) | 0.576 (0.373–0.700) | <u>0.603 (0.407–0.740)</u> |
| TCGA-CESC | **0.597 (0.554–0.671)** | 0.515 (0.370–0.627) | <u>0.543 (0.477–0.589)</u> | 0.519 (0.381–0.629) |
| TCGA-STAD | **0.545 (0.465–0.610)** | 0.479 (0.410–0.534) | 0.471 (0.438–0.495) | <u>0.511 (0.468–0.560)</u> |

**Table S60**: **Survival prediction for individual cohorts.** For each cohort, we evaluated model performance using a five-fold cross-validation scheme. Patients within each cohort were first split into five folds. In each iteration, the model was trained on four folds and evaluated on the remaining fold, with the process repeated until every fold had served once as the evaluation set. Pre-extracted tile- or slide- level features from each section were used to train a shallow survival prediction network separately for each repetition. Test performance was reported using mean C-index across all folds. The best and second-best metrics are highlighted in bold and underlined, with the minimum and maximum values across all fold provided in parentheses. This table reports cohort-specific results for all evaluated cohorts, whereas the main text highlights the principal overall findings.

| Cohort | N patients | Median tissue area | CIELAB color variance | mC-index |
|---|---|---|---|---|
| TCGA-CESC | 269 | 117.0 $mm^2$ | 110.8 | 0.60 |
| CGODC-OV | 85 | 121.4 $mm^2$ | 35.2 | 0.62 |
| TCGA-OV | 106 | 203.1 $mm^2$ | 166.0 | 0.58 |
| TCGA-STAD | 416 | 149.5 $mm^2$ | 153.3 | 0.55 |

**Table S61**: **Evaluation of SQUALL generalizability across clinical cohorts with diverse tissue-scale and staining characteristics.** TCGA-OV was additionally included to evaluate the effects of whole-slide tissue scale and staining heterogeneity on model performance. Mean five-fold cross-validation results were shown. Compared with CGODC-OV, TCGA-OV exhibited a larger median tissue area and greater staining variation, as quantified by the total variance in CIELAB color space. Similar challenges were observed in TCGA-STAD. These results suggest that model performance may be influenced by tissue scale and morphological or staining variations.

| | | **Platinum-based chemotherapy resistance** | | | **Time to recurrence** |
|---|---|---|---|---|---|
| **Cohort** | **Model** | F1 | mACC | AUROC | C-index |
| CGODC-OV | SQUALL | **0.690 (0.585–0.814)** | 0.795 (0.750–0.882) | 0.574 (0.417–0.865) | **0.558 (0.429–0.778)** |
| | UNI | 0.633 (0.566–0.734) | 0.796 (0.750–0.875) | 0.534 (0.385–0.635) | 0.418 (0.357–0.464) |
| | PLIP | 0.632 (0.554–0.690) | 0.793 (0.750–0.824) | 0.443 (0.188–0.654) | 0.526 (0.500–0.593) |
| | Virchow | 0.629 (0.489–0.803) | **0.817 (0.750–0.875)** | **0.588 (0.344–0.698)** | 0.429 (0.333–0.556) |
| TCGA-OV | SQUALL | **0.668 (0.417-0.778)** | **0.786 (0.750-0.875)** | **0.627 (0.400-0.778)** | - |
| | UNI | 0.604 (0.385-0.750) | 0.689 (0.500-0.875) | 0.446 (0.267-0.667) | - |
| | PLIP | 0.575 (0.417-0.750) | 0.692 (0.625-0.750) | 0.453 (0.278-0.667) | - |
| | Virchow | 0.557 (0.350-0.750) | 0.742 (0.667-0.875) | 0.543 (0.278-0.867) | - |
| **Overall** | SQUALL | **0.679 (0.417–0.814)** | **0.791 (0.750–0.882)** | **0.601 (0.400–0.865)** | – |
| | UNI | 0.619 (0.385–0.750) | 0.743 (0.500–0.875) | 0.490 (0.267–0.667) | – |
| | PLIP | 0.604 (0.417–0.750) | 0.743 (0.625–0.824) | 0.448 (0.188–0.667) | – |
| | Virchow | 0.593 (0.350–0.803) | 0.780 (0.667–0.875) | 0.566 (0.278–0.867) | – |

**Table S62**: **Platinum-based chemotherapy resistance prediction for individual cohorts.** Patients with platinum status were included in this prediction task. For each cohort, we evaluated model performance using a five-fold cross-validation scheme. Patients within each cohort were first split into five folds. In each iteration, the model was trained on four folds and evaluated on the remaining fold, with the process repeated until every fold had served once as the evaluation set. Pre-extracted tile or slide level features from each section were used to train a shallow prediction network separately for each repetition. Test performance was reported using F1 score, mean accuracy (mACC.), AUROC, and C-index across all fold. The best and second-best metrics are highlighted in bold and underlined, with the min and max value across all fold provided in parentheses.
